# Electrophilic Multi-Kinase Probes Identify Degradation and Aspartate Conjugation of ULK1

**DOI:** 10.64898/2026.04.30.722011

**Authors:** Nur Mehpare Kocaturk, Adam L. Pinto, Matylda Izert-Nowakowska, Lea P. Wilhelm, Gajanan Sathe, Qasim Ashraf, Ian G. Ganley, Adrien Rousseau, William Farnaby

**Affiliations:** Centre for Targeted Protein Degradation, Faculty of Life Sciences, University of Dundee, Dundee, DD1 5JJ, U.K.; MRC Protein Phosphorylation and Ubiquitylation Unit, Faculty of Life Sciences, University of Dundee, Dundee DD1 5EH, Scotland, U.K.; Molecular Cell and Developmental Biology, Faculty of Life Sciences, University of Dundee, Dundee, DD1 5EH, U.K.

## Abstract

Kinases have proven to be one of the most fertile target classes for new drug approvals, however, classical reversible inhibitors may not be capable of the levels of specificity or target modulation required across a broad spectrum of disease areas. Approaches that chemically modify kinase inhibitors in solvent exposed regions are unveiling a swathe of mechanisms to address kinase function in new ways. For example, by either covalently recruiting nucleophilic residues outside of the ATP-binding pocket to inhibit, or by recruiting secondary effector proteins to degrade. Here, we established a systematic workflow to assess the cellular impact of minimal electrophilic modifications to ATP-site binding scaffolds. Amongst multiple potential starting points for monovalent kinase degradation that were revealed, we identified that one of our scout molecules was able to control the abundance of the master regulator of autophagy, Unc-51-like autophagy activating kinase 1 (ULK1) and that a scaffold-hop could be performed to generate a targeted monovalent ULK1 degrader chemical tool, we named ALP-1. Remarkably, further investigation of how ALP-1 engages ULK1 unveiled an atypical conjugation event between the chloroacetamide warhead of ALP1 and an aspartate residue on the lip of the ULK1 ATP-binding site. These discoveries reveal new understanding of the potential for non-cysteine reactivity of the commonly employed chloroacetamide warhead, as well as new opportunities for the targeting and covalent engagement of kinases.

## Introduction

Since the approval of imatinib in 2001 for chronic myeloid leukaemia there have been over 70 further approvals of kinase inhibitors, underlining the importance of this target class^1^. These have featured an ever-expanding collection of approved targeted covalent inhibitors to address, for example, cancer resistance mutations that increase ATP affinity and render reversible inhibitors less effective^2, 3^. More recently, there has been a growing presence in early clinical phases of kinase targeted protein degraders, which can leverage the advantages of non-occupancy-based mechanisms of action, high levels of target specificity and removal of non-catalytic protein function^4, 5^. Degrader molecules are typically classified as either heterobifunctionals or molecular glues. Heterobifunctionals usually feature a moderate-to-high affinity ligand for the target protein, which is chemically linked to a recognised ligand for an E3 ligase. Molecular glues, on the other hand, bind with measurable affinity to only one, or neither, of the target protein or E3 ligase. Their function therefore relies on facilitating protein-protein interactions that ultimately drive target protein degradation. These approaches offer the potential to drive a new wave of kinase targeted therapies. For example, to address challenges with isoform specificity that is beyond the capability of classical inhibitors, or to penetrate into underrepresented disease areas for kinase targeted therapies, such as the central nervous system^4, 6^ and inflammation linked conditions^7^.

Approaches to kinase targeted molecular glue discovery have been rapidly evolving. The immunomodulatory drug Lenalidomide drives Casein Kinase 1α degradation^8, 9^ via recruitment of the Cul4 Cereblon (CRBN) E3 ligase recognition sub-unit, inspiring approaches to harness CRBN to degrade other kinases, including the WEE1 degrader BMS-986463^10^, currently in clinical trials. Alternatively, it has also been shown that kinase inhibitors can be functionalised either directly^11, 12^, or via short linkers^13^, with electrophilic handles to enact targeted protein degradation. Some trends have emerged here, including repeated identification of DCAF16^12^ and FBOX22^13^ as being engaged to drive target degradation, consistent with electrophilic targeted molecular glue approaches for other protein classes^14–17^. These studies primarily focus on the potential for *trans*-recruitment of proteins, such as E3 ligases, associated with cellular degradation machinery. In the context of targeted inhibition, solvent exposed reactive handles have also been used to *cis*-label, conjugating to nucleophilic amino acid residues near to the lip of an ATP-binding site. This has included development of chemoproteomic tools for profiling kinase inhibitor specificity^18^ and inhibitors that target residues outside of the catalytic pocket for potency and specificity gains^19^.

Unc-51-like autophagy activating kinase 1 (ULK1) is a serine/threonine kinase that is critical for autophagy initiation^20^, functioning within the ULK1 complex with ATG13, focal adhesion kinase family interacting protein (FIP200) and ATG101. ULK1 is believed to act as a signalling hub, regulated by the energy and nutrient sensing mTOR/AMPK axis. Understanding ULK1 disease relevant signalling and therapeutic prosecution is a highly active area. This includes clinical trials investigating the utility of ULK1 inhibitors in patients with advanced or metastatic solid tumours in non-small cell lung, pancreatic and colorectal cancers^21^. To date most attention has continued to focus on reversible ULK1 inhibitors^22, 23^.

In this study, we utilised multi-kinase targeting scout molecules for assessing functional labelling events in cells. Using both focussed and chemoproteomic kinase abundance profiling, we identified a series of starting points and revealed ULK1 as a degradation ‘hit’, before successfully scaffold hopping the reactive chloroacetamide handle to an ULK1 targeted binding motif. Compound induced ULK1 degradation was shown to be E1 ligase and lysosome, rather than proteasome, dependent. Additionally, and unexpectedly, we found that our lead molecule, ALP1, covalently engages the ULK1 kinase domain, rather than recruiting an effector protein via its electrophile. Further investigation revealed that instead of engaging a cysteine residue, as expected of a chloroacetamide warhead, this electrophile engages an aspartate residue on the D-helix of the ULK1 kinase domain. This study therefore reveals a previously unappreciated reactivity of this commonly employed electrophile and highlights new opportunities for covalent engagement of kinases via acidic amino acids residues. In addition, our lead degrader ALP-1 offers a differentiated starting point for generating ULK1 probes that control the abundance and function of the master autophagy regulator, ULK1.

## Results

### Design, Synthesis and Cellular Screening of Electrophilic Probes Identifies Modulated Kinases

To address bottlenecks in the discovery of molecular glues, we posited that by modifying a promiscuous kinase inhibitor with solvent-exposed electrophiles, we could identify kinases amenable to degradation by these molecules and use them as starting points for the development of targeted molecular glues. Critically, as covalent binding occurs at the effector, the prototypical catalytic mechanism of action of a molecular glue is retained. Alternatively, irreversible labelling of a solvent front residue of a kinase is a possibility, whereby *cis* modification may result in altered protein abundance or function (**Fig. 1A**).

**Figure 1.**
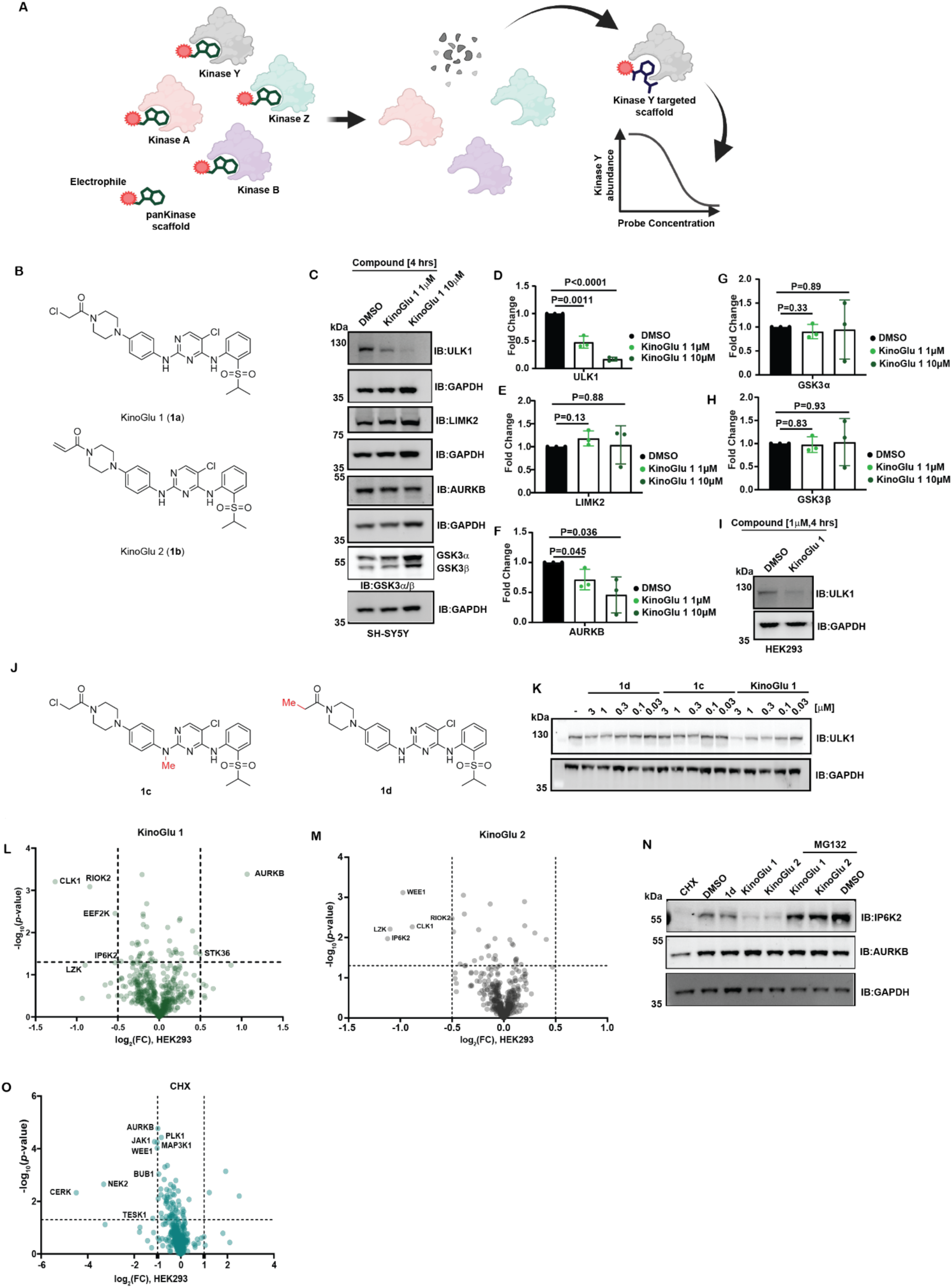
Initial scouting of kinases impacted by electrophilic probes. (**A**) Schematic representation of using promiscuous kinase scout molecules as a basis for targeted kinase degrader discovery. Created in BioRender. Farnaby, W. (2026) https://BioRender.com/mxro3i7 (**B**) Structures of KinoGlu 1 (**1a**) and KinoGlu 2 (**1b**). (**C**) Effect of KinoGlu 1 (1 and 10 µM, 4 h) on selected endogenous kinases in SH-SY5Y cells. Immunoblots are representative of n=3 biological replicates. Relative quantification of (**D**) ULK1, (**E**) LIMK2, (**F**) AURKB, (**G**) GSK3α and (**H**) GSK3β immunoblots normalised to internal controls in SH-SY5Y cells following treatments with KinoGlu 1 (n=3 biological replicates, mean, ±SD). (**I**) The effect of KinoGlu 1 (1 µM, 4 h) on endogenous ULK1 levels in HEK293 cells. Immunoblots are representative of n=3 biological replicates. (**J**) Structures of negative controls; **1c** and **1d**. (**K**) Dose-dependent effect of **1d**, **1c** and KinoGlu 1 (indicated doses, 4 h) on ULK1 levels in SH-SY5Y cells. Immunoblots are representative of n=3 biological replicates. Proteome-wide activity profiling of (**L**) KinoGlu 1 and (**M**) KinoGlu 2 normalised to negative control (**1d**) on kinase level in HEK293 cells (n=3 biological replicates, 1 µM, 4 h). (**N**) Effects of KinoGlu 1 (1 µM, 4 h), KinoGlu 2 (1 µM, 4 h) alone or in combination with MG132 (1 µM, 2 h pre-treatment then co-treatment with test compounds for additional 4 h), **1c** (1 µM, 4 h), cycloheximide (CHX, 100 µg/ml, 4 h) on identified kinase hits from TMT-based proteomics profiling (**L** and **M**). Immunoblots are representative of n=3 biological replicates. (**O**) Identification of short-lived kinases using label-free proteomics in HEK293 cells (n=3 biological replicates, CHX; 100 µg/ml, 4 h).

As a starting point, we synthesised molecules based on the TL13-87 kinase binding motif,^24^ with solvent exposed chloroacetamide or acrylamide moieties, which we here refer to as KinoGlu 1 (**1a**) and KinoGlu 2 (**1b**) respectively (**Fig. 1B**). We first screened KinoGlu 1 in SH-SY5Y cells at 1 μM for 4 h, a relatively low concentration and short timepoint chosen to reduce artefactual readouts associated with effects on transcription, rather than post-translational kinase modulation. As a pilot study we initially assessed the effect of treatment with KinoGlu 1 against a limited selection of kinases by western blotting, including those that had been observed to be most often degraded by Donovan et al. when using a set of multi-kinase bifunctional degraders^25^ (**Fig. 1C-H**). Of the kinases screened against, ULK1 stood out as significantly downregulated by KinoGlu 1 and this effect was further validated in HEK293 cells (**Fig. 1I**). To assess the dependency of ULK1 degradation on kinase engagement and electrophile presence, compounds **1c** and **1d** (**Fig. 1J**) were synthesised. In **1c** kinase binding was abrogated by N-methylation of the nitrogen which forms the key hinge-binding hydrogen-bonding interaction, and in **1d** the chloroacetamide electrophile was exchanged for an unreactive propanamide moiety. A 4 h treatment of KinoGlu 1 over a concentration range revealed dose-dependent degradation of ULK1 in SH-SY5Y cells, whilst the control compounds **1c** and **1d** were non-degraders, demonstrating the presence of both the electrophile and kinase hinge-binding moiety to be necessary for ULK1 degradation (**Fig. 1K**).

Next, we sought to establish a chemoproteomic pipeline to assess the effects of KinoGlus 1 and 2 on kinome-wide abundance. HEK293 or SH-SY5Y cells were treated with electrophilic compounds, inhibitor only control **1d** or DMSO for 4 h. We employed tandem mass tag (TMT) proteomics to enable us to quantify the effect of electrophilic compound treatment on kinase abundance compared to both DMSO (**Fig. S1A-C**) and the non-electrophilic control (**Fig. 1L, M)**. We included this control up-front due to the impact kinase inhibitors can impart on the stability of their targets^26^ and we were keen to understand the specific effects of the covalent modifications we had made. Using significance cut-offs of abs(log2[FC]) >0.5 and *p*-value <0.05, KinoGlu 1 treatment led to downregulation of CLK1, eEF-2K and RIOK2, whilst AURKB abundance was increased (**Fig. 1L**). Upon treatment with KinoGlu 2, five kinases were identified as decreased in abundance: CLK1, IP6K2, LZK, RIOK2 and WEE1 (**Fig. 1M**). KinoGlus 1 and 2 did not cause any drastic or unexpected changes in protein abundance across the total proteome after a 4 h treatment (**Fig. S1D-H**). We followed on by focusing on the validation of kinases that were modulated in abundance compared to both DMSO (**Fig. S1A-B**) and the inhibitor control (**Fig. 1L**). For example, CLK1 showed reduction as compared with the control compound but not compared with DMSO, suggesting the multi-kinase inhibitor scaffold alone increases CLK1 abundance.

We were interested to observe degradation of RIOK2, an atypical serine threonine kinase, by KinoGlu 2. Work of Ma et al.^27^ was recently published in which they develop CQ-211^28^, a selective RIOK2 inhibitor, into compound 9c, a degrader, by addition of a solvent exposed acrylamide. The 4 µM DC50 at 12 h seen in their study is well matched with our observation of modest downregulation upon 1 µM treatment of KinoGlu 2 for 4 h and gave us confidence that our approach could identify bona fide starting points for targeted kinase degrader discovery.

To further validate potential hits western blotting was carried out, and the effect of compound treatment was also compared to that of cycloheximide shut down of protein synthesis. Different to our observations by proteomics, AURKB levels were found not to significantly increase upon KinoGlu 1 treatment compared to the inhibitor control. We also noted that treatment of both KinoGlus 1 and 2 led to downregulation of IP6K2, however, this was also seen in the cycloheximide treatment arm (**Fig. 1N**). We therefore surmised that IP6K2 degradation was not necessarily a post-translational effect and it was therefore considered as a high risk hit kinase to continue with and deprioritised for follow-up.

LZK is a kinase associated with regulation of neuronal degeneration and axon growth and dual DLK/LZK inhibitors have been studied as potential amyotrophic lateral sclerosis (ALS) treatments^29, 30^. Selective LZK targeting has proven challenging and so we were interested to see that the abundance of this protein was lowered significantly versus both DMSO and control **1d**. However, commercially available antibodies could not be robustly validated to orthogonally quantify these effects and the low abundance of LZK limited us in follow-up MS based studies. Nevertheless, this remains an interesting putative hit.

Considering the relatively low abundance of cysteine residues in the human proteome, we extended the compound set to include four more electrophiles with reactivity towards other nucleophilic amino acids (**Fig. S2A**). To assay this larger compound set we moved to a label-free proteomics approach. Also included, was a cycloheximide treatment arm to enable quick identification of short-lived kinases (**Fig. 1O**), the importance of which was highlighted by the apparent IP6K2 degradation seen previously. Treatment with cycloheximide showed some overlap of modulated kinases upon treatment with KinoGlu 2, including WEE1, highlighting this as a potentially ‘high-risk’ hit for follow-up studies. Despite the extended compound set with differential amino acid reactivity, few kinases were identified as hits (Supplementary data file 1). Therefore, we decided to focus further efforts on the ULK1 degradation activity of KinoGlu 1.

### Scaffold Hopping to Identify an ULK1 Targeted Degrader

Encouraged by the activity of KinoGlu 1 against ULK1, we sought to generate electrophilic degraders based on ULK1/2 targeted scaffolds. Compounds were designed based on known ULK1 inhibitors SBP-7455^31^ and MRT68921^32^. Comparison of the crystal structures of MRT68921 with ULK1 and SBI-0206965 (an SBP-7455 related compound)^31^ with ULK2 to a close analogue of TL13-87 with FAK^33^, enabled identification of exit vectors to maintain the positioning of the chloroacetamide electrophile within the solvent exposed region (**Fig. 2A**). The solvent facing position of the tetrahydroisoquinoline (THIQ) group of MRT68921 and phenyl ring of SBP-7455 were noted as reasonable exit vectors. As the use of a secondary amine was preferred for appending the chloroacetamide, we incorporated or retained the THIQ to generate ALP-1 (**2a**) and **2b** (**Fig. 2B**). The activity of these compounds was assessed by western blotting in SH-SY5Y cells. ALP-1, which utilised the SBP-7455 core demonstrated similar degradation activity to KinoGlu 1, whilst the MRT68921 core bearing compounds did not consistently demonstrate ULK1 degradation at this concentration and time-point (**Fig. 2C, F**).

**Figure 2.**
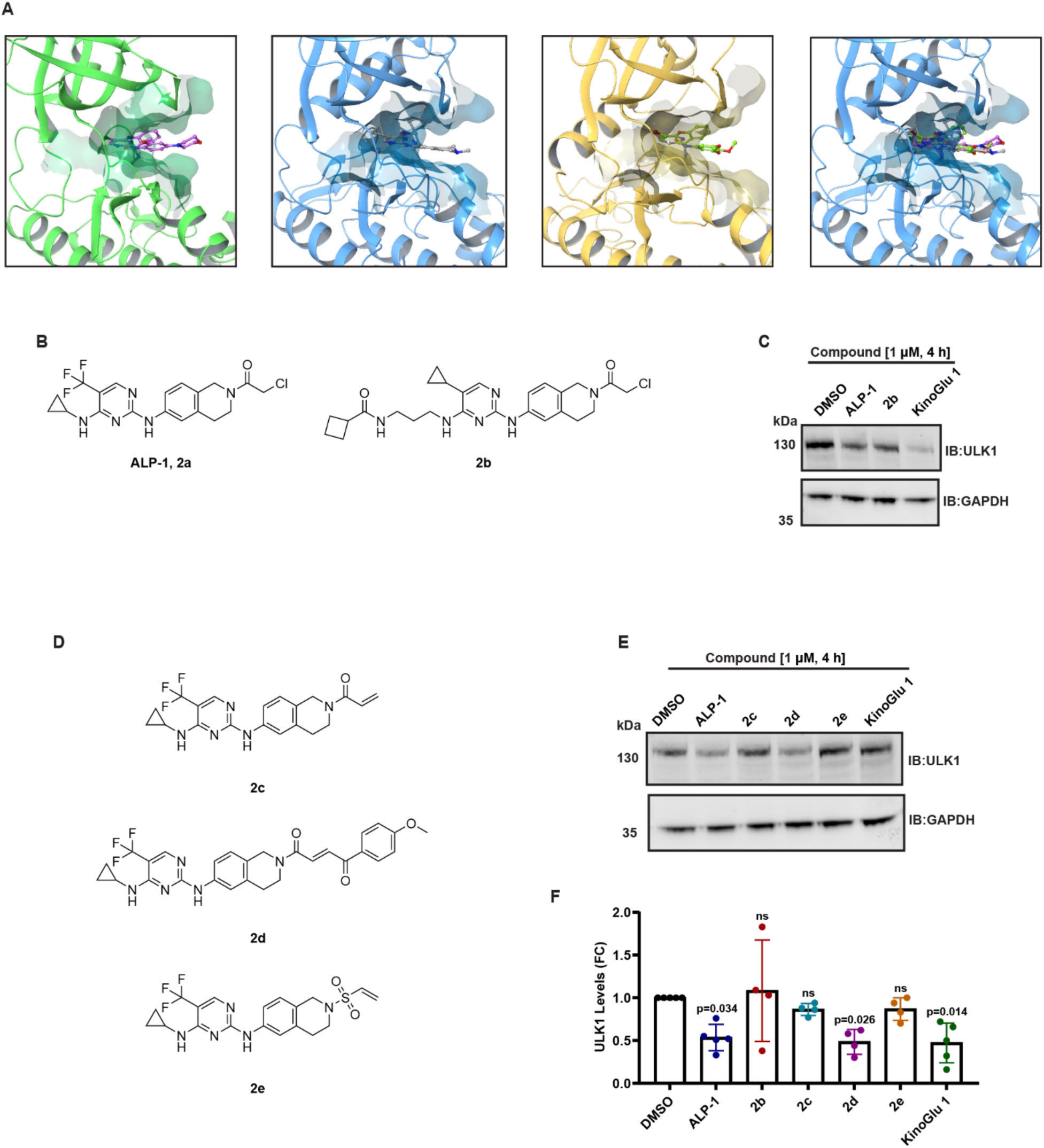
Design and cellular profiling of targeted ULK1 degraders. (**A**) Crystal structures of kinase domains of FAK in complex with TAE226 (PDB: 2JKK) (far left), ULK1 with MRT68921 (PDB: 8P5K) (centre left), ULK2 with SBI-0206965 (PDB: 6YID) (centre right), overlay of aligned binding sites (far right) (**B**) Structures of ALP-1 (**2a**) and **2b.** (**C**) The effect of ALP-1, **2b** and KinoGlu 1 (1 µM, 4 h) on ULK1 levels in SH-SY5Y cells. Images are representatives of n=4 biological replicates. (**D**) Structures of **2c**, **2d** and **2e.** (**E**) The effect of **2c**, **2d** and **2e** compared to ALP-1 and KinoGlu 1 (1 µM, 4 h) on ULK1 levels in SH-SY5Y cells. (F) Relative quantification of ULK1 immunoblots from panels C and E normalised to internal controls in SH-SY5Y cells upon treatment with ALP-1, **2b**, **2c**, **2d**, **2e** and KinoGlu 1 (1 µM, 4 h) (n≥3 biological replicates, mean, ±SD).

Next, we explored how modification of the electrophile on the ALP-1 core affected ULK1 degradation. The chloroacetamide was exchanged for an acrylamide (**2c**), β-ketoenamide (**2d**)^11^, or vinyl sulfonamide (**2e**) electrophile (**Fig. 2D**). In the case of the acrylamide and vinyl sulfonamide no obvious degradation of ULK1 was observed, however, the β-ketoenamide compound degraded ULK1 similarly to ALP-1 (**Fig. 2E, F**). As such, both ALP-1 and **2d** were taken forward for further characterisation.

### ALP-1 Requires Both Hinge Binding Interaction and Electrophile to Shut Down ULK1 Specific Signalling

In the knowledge that we could transplant the covalent handle to a targeted ULK1 scaffold and retain degradation activity, control compounds based on ALP-1 and **2d** were synthesised. As with KinoGlu 1, these control compounds consisted of kinase binding being abrogated by N-methylation of the hinge-binding nitrogen (**3a** and **3b**); or the electrophilic warhead being replaced with a propanamide (**3c**) (**Fig. 3A**). These compounds were characterised in a C-terminal HiBiT-tagged ULK1 Knock-In (KI) HEK293 cell line. The HiBiT tag enabled rapid detection of protein abundance via a luminescence-based readout^34^. Removal of the electrophile resulted in no degradation of ULK1-HiBiT up to 10 μM compound concentration. N-methylation of ALP-1 also significantly abolished ULK1 degradation (**Fig. 3B**), however, the same extent of abrogation of ULK1 degradation was not observed for **3b** (**Fig. 3C, S3A**). For this reason, we decided to prioritise ALP-1 for further study.

**Figure 3.**
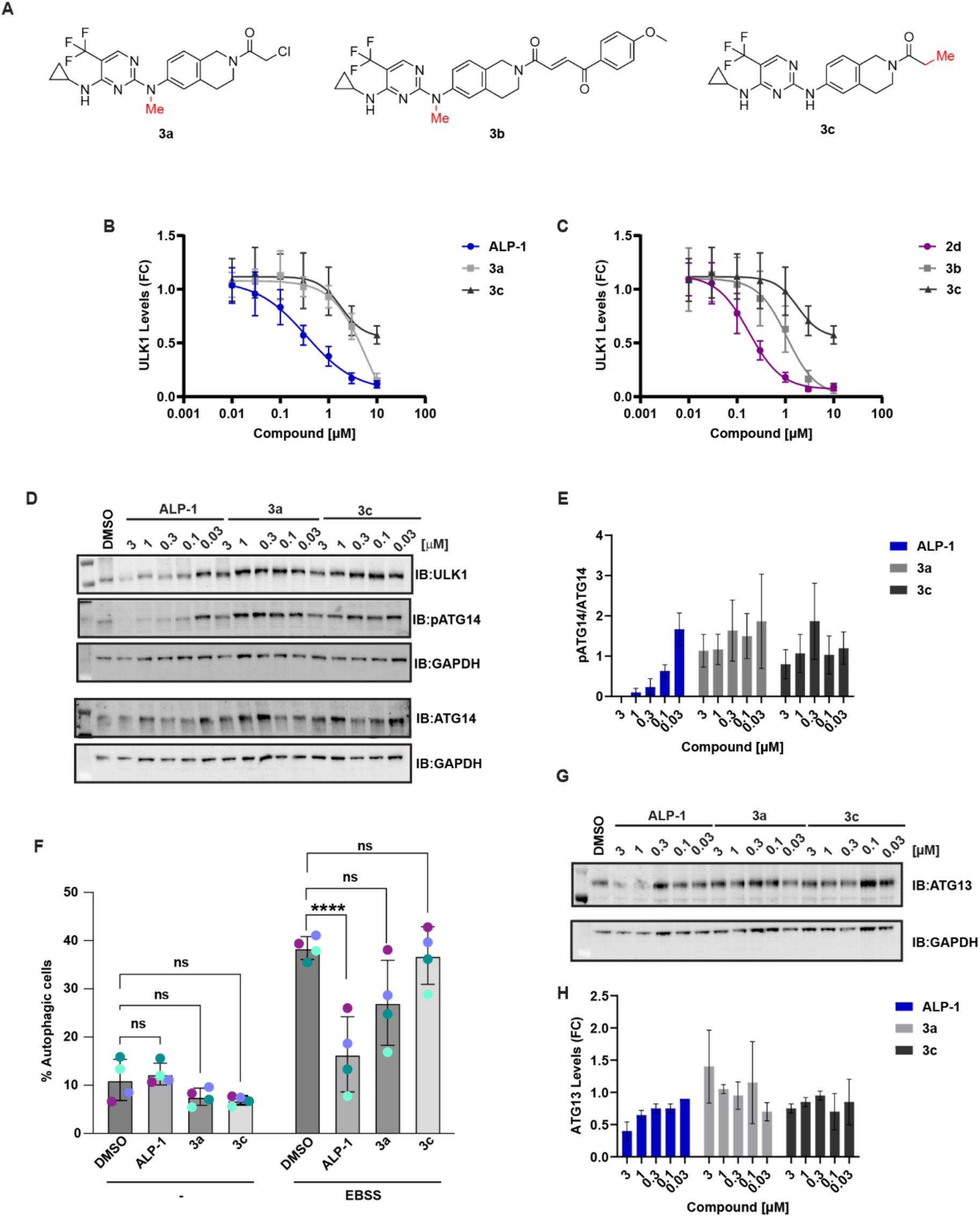
Cellular effects of ALP-1 on ULK1, ULK1 mediated signalling and autophagy induction. (**A**) Chemical structures of **3a**, **3b** and **3c**. (**B**) Dose-dependent degradation profile of ALP-1 and negative controls (**3a** and **3c**, 8 h) in ULK1-HiBiT KI HEK293 cells. Degradation curves were generated with non-linear 4 parameter fit correction (Dose response curves are means of at least n=3 biological replicates, ± SD). (**C**) Dose-dependent degradation profile of **2d** in comparison to negative controls (**3b** and **3c**, 8 h) in ULK1-HiBiT KI HEK293 cells. Degradation curves were generated with non-linear 4 parameter fit correction (Dose response curves are means of at least 3 biological replicates, ± SD). (**D**) Dose dependent effects of ALP-1, **3a** and **3c** (indicated doses, 8 h) on endogenous ULK1, basal and phosphorylated ATG14 levels in SH-SY5Y cells. Immunoblots are representative of n=3 biological replicates. (**E**) Relative quantification of pATG14/ATG14 ratios from immunoblots normalised to internal controls and DMSO upon treatment with ALP-1, **3a** and **3c** (indicated doses, 8 h) (n=3 biological replicates, mean, ±SD). (**F**) Quantification of autophagy levels using Auto-*QC* reporter by flow cytometry (n=4 biological replicates); statistics: two-way ANOVA with Tukey’s multiple-comparisons test. (**G**) Dose dependent effects of ALP-1, **3a** and **3c** (indicated doses, 8 h) on endogenous ATG13 levels in SH-SY5Y cells. (**H**) Relative quantification of ATG13 from immunoblots normalised to internal controls and DMSO upon treatment with ALP-1, **3a** and **3c** (indicated doses, 8 h) (n=2 biological replicates, mean, ±SD).

The potency of ALP-1 was measured in ULK1-HiBiT KI HEK293 cells to have a DC50 value of 340 nM and Dmax of 95% at 8 h. The effect on cell viability was also measured using a fluorescent GF-AFC substrate to provide a readout of protease activity as a surrogate for viability^35^. No cytotoxicity was observed for ALP-1 below a 10 µM treatment concentration (**Fig. S3B**). Dose courses of ALP-1 and controls in SH-SY5Y cells demonstrate that, similarly, endogenous ULK1 is robustly degraded by ALP-1 but no effect is observed with **3a** or **3c** controls (**Fig. 3D**). Likewise, using TMT-MS we observed degradation of ULK1 by ALP-1 (**Fig. S3C**). We also noted significant impact on some other kinases not previously observed with KinoGlu 1, which may be partly attributed to a combination of increased time and dose, as well as a different binding scaffold.

To understand the effect of ALP-1 on ULK1 signalling, we chose to quantify the effect of compound treatment on ser-29 phosphorylation of ATG14, an ULK1 specific substrate site.^36^ ULK1 mediated phosphorylation of ATG14 is upregulated under autophagy induction, and is required for autophagosome formation, therefore levels of pATG14 also act as an indicator for modulation of autophagy initiation. Treatment with ALP-1 led to robust reduction of pATG14 levels, with total inhibition of ATG14 phosphorylation at 1 µM, and significantly reduced levels even at 100 nM. This shut-down of ATG14 phosphorylation was not observed upon treatment with ULK1 inhibitor **3c** at matched concentrations (**Fig. 3D, E**), demonstrating that ALP-1 can reduce downstream ULK1 signalling to a much greater extent than its parent inhibitor (**3c**).

Building on this, we assessed the effect of ALP-1 on starvation-induced autophagy. To do so, we used ULK1/2 double-knockout MEF cells^22, 37^ stably expressing the autophagy reporter mCherry–GFP–LC3 (referred to as *Auto*-QC reporter) (**Fig. S3D**). Upon autophagy induction, autophagosomes form that are positive for both GFP and mCherry. Following fusion with lysosomes to generate autolysosomes, the acidic lysosomal environment quenches the GFP signal, while the mCherry signal remains stable. Consequently, autophagic flux can be quantified as an increase in the mCherry/GFP ratio (**Fig. S3D, E**). We stably reintroduced FLAG-tagged ULK1 into these cells and pretreated them with 1 µM ALP-1, **3a**, or **3b** for 4 h, followed by a further 2 h incubation in amino acid–free Earle’s Balanced Salt Solution (EBSS) in the continued presence of each compound. As previously observed in SH-SY5Y cells, ALP-1 decreased ULK1 levels under these conditions (**Fig. S3F**). We observed no effect of any compounds on basal autophagy, however, when combined with EBSS starvation, robust autophagy induction was observed in the presence of the negative control compounds **3a** and **3c**, or DMSO (**Fig. 3F, S3E**). In contrast, cells treated with ALP-1 exhibited a significant reduction in starvation-induced autophagy compared with the other conditions, consistent with its effects on ULK1 and pATG14 levels.

Having noted that ALP-1 impairs starvation induced autophagy, we sought to understand the effect of compound treatment on the ULK1 complex. Modest downregulation of ATG13, an ULK1 complex member, was observed upon high concentration ALP-1 treatment (**Fig 3G, H**). This was not observed to the same extent with inhibitor **3c** or negative control **3a**, suggesting that with increased clearance of ULK1 a decrease in stability of the entire ULK1 complex is seen, as may be expected.

Together, these data support that ALP-1 suppresses ULK1 signalling and autophagy induction beyond the level observed with the non-electrophilic inhibitor control.

### ALP-1 Mediated ULK1 Degradation is Dependent on E1 Enzyme and Lysosome Activity

Next, we investigated the mechanism by which ALP-1 degrades ULK1. We first sought to confirm the downregulation of ULK1 was post-translational. A 10 h time-course experiment was carried out, in which ULK1-HiBiT KI HEK293 cells were treated with either ALP-1, or cycloheximide (CHX) to shut down protein synthesis. This data shows that natural rate of decay of ULK1 levels in this setting is greater than 10 hours and as the onset of degradation was significantly faster upon ALP-1 treatment compared to CHX, we could confirm the effect of the molecule was truly post-translational (**Fig. 4A**).

**Figure 4.**
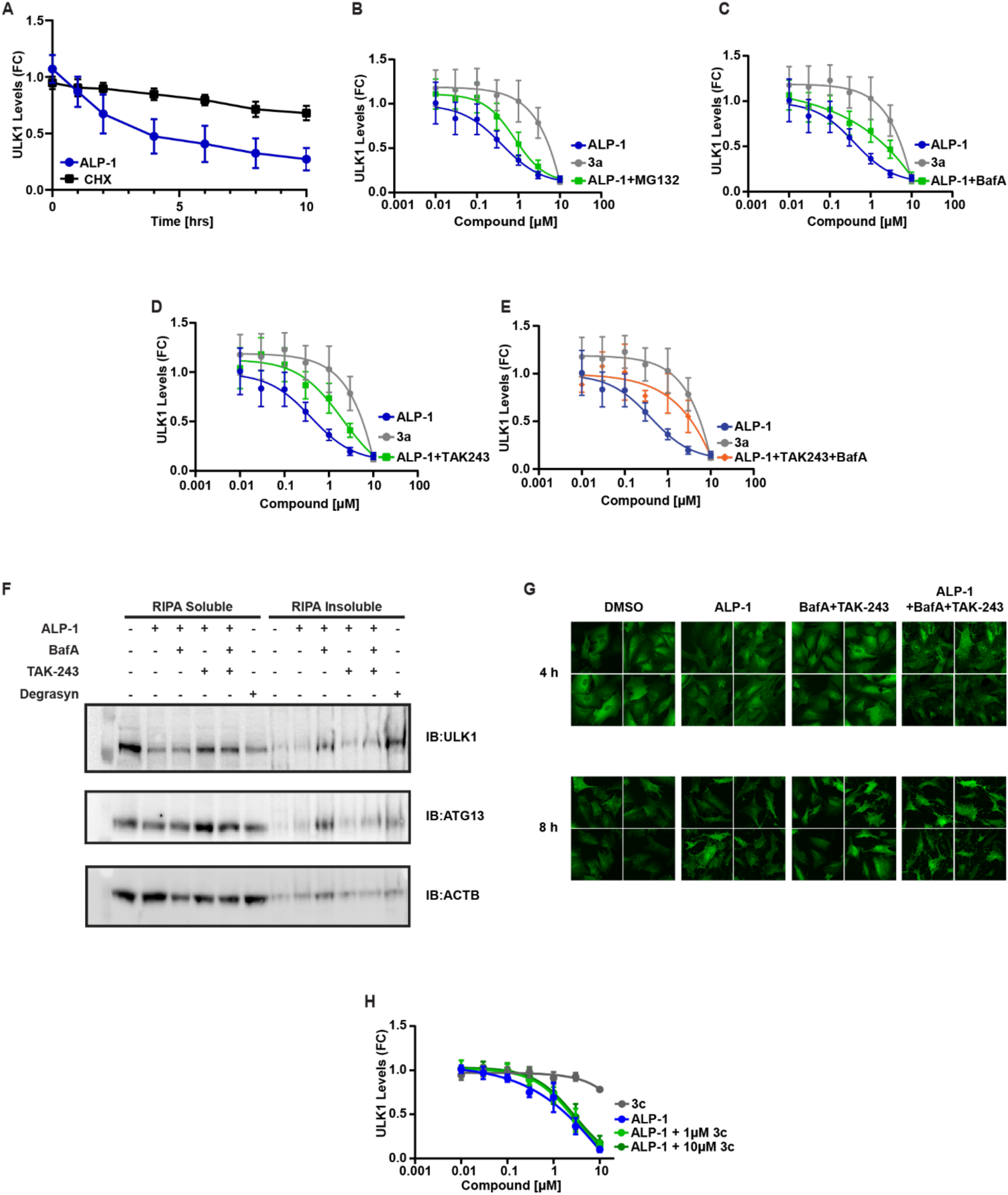
ALP-1 mechanism of action studies. (**A**) Degradation profile of ALP-1 (3 µM) in comparison with CHX (100 µg/ml) in ULK1-HiBiT KI HEK293 cells for 10 h. Representative graph is combined result of n=3 biological replicates. Mode-of-action studies of ALP-1 (10, 3, 1, 0.3 and 0.1 µM) and in combination with (**B**) MG132 (1 µM), (**C**) Bafilomycin A1 (BafA1, 100 nM), (**D**) TAK-243 (1 µM) and (**E**) TAK-243 (1 µM) with Bafilomycin A1 (100 nM) in ULK1-HiBiT KI HEK293 cells were co-treated with ALP-1 and inhibitors for 8 h. Dose-dependent degradation curves were generated with non-linear 4 parameter fit correction (Dose response curves are means of n=3 biological replicates, ± SD). (**F**) Western blot of ULK1 and ATG13 levels in the RIPA soluble and insoluble fractions of SH-SY5Y cells upon 8 h treatment with ALP-1 (3 µM), BafA1 (100 nM), TAK-243 (1 µM), combinations thereof, or Degrasyn (5 µM). Blots representative of n=3 biological replicates. (G) Representative microscopy images of ARPE19 cells stably expressing GFP-ULK1, treated with ALP-1 (1 μM) alone, BafA1 (100 nM) and TAK-243 (1 µM) in combination, or ALP-1 (1 μM) together with BafA1 (100 nM) and TAK-243 (1 µM). Insets show higher-magnification views. Scale bars, 10 μm. n = 3 biologically independent experiments. (**H**) Competition experiment of ALP-1 with parent inhibitor **3c** (1 µM and 10 µM), ULK1-HiBiT KI HEK293 cells were pre-treated with **3c** for 2 h, followed by 4 h co-treatment with ALP-1.

We then attempted to discern whether ULK1 degradation was mediated by the ubiquitin-proteasome system (UPS), however, responses across a dose course of ALP-1 with and without co-treatment of proteasome inhibitor MG132 overlapped, with no clear rescue observed in ULK1-HiBiT KI HEK293 cells (**Fig. 4B**). In line with this observation, when performing IP-MS analysis in SH-SY5Y cells stably expressing Flag-ULK1 to identify ALP-1 induced protein-protein interactions, no E3 ligases were enriched (**Fig. S4A-F**). We also demonstrated that ULK1 degradation could not be reversed using pan-caspase inhibitor Z-VAD-FMK, showing disappearance of ULK1 was independent of caspase activity and apoptosis (**Fig. S4G**). We looked at the possibility of degradation via the lysosomal pathway through co-treatment of ULK1-HiBiT KI HEK293 cells with Bafilomycin A1 (BafA1). This revealed a clear partial rescue of degradation at 1 and 3 µM concentrations (**Fig. 4C**). Likewise, co-treatment of ALP-1 with TAK-243, a UBA1 inhibitor, demonstrated rescue across a dose course (**Fig. 4D**), as did simultaneous co-treatment with both BafA1 and TAK-243 (**Fig. 4E, Fig. S4H, I**). Taken together these data support no obvious dependence on proteasome activity, however, both ubiquitination and lysosomal activity are necessary for ALP-1 mediated degradation of ULK1.

Aware that it has previously been shown that electrophilic compounds may cause non-specific aggregation of proteins, leading to apparent degradation in detergent soluble cellular fractions^38^, we assessed the detergent-insoluble fraction for accumulation of ULK1 and ATG13 (**Fig. 4F**). No accumulation of either protein was observed upon ALP-1 treatment. Nor was any greater accumulation observed when ALP-1 was co-treated with BafA1 and/or TAK-243, than when these inhibitors were treated alone (**Fig. S4J**). Degrasyn was used as a positive control in this experiment as it has been demonstrated previously to cause ULK1 aggregation^39^.

To further evaluate the effects of ALP-1 on ULK1, we visualised ULK1 localisation using confocal microscopy (**Fig. 4G**). When treating these cells with 1 µM ALP-1 for 4 hours we observed ULK1 puncta formation. Further accumulation of these puncta was observed upon co-treatment with BafA1 and TAK-243, confirming that concomitant inhibition of UBA1 and autophagy impedes ALP-1 induced ULK1 clearance. This supports a model in which ALP-1 induces the formation of soluble ULK1 puncta whose clearance relies on E1 and lysosome activity.

As a final control experiment, we tested whether ULK1 degradation could be abolished by competition with parent inhibitor **3c**. Surprisingly, we found that degradation caused by concentrations of ALP-1 up to 10 µM could not be off-competed with a 2 h pre-treatment of 1 or 10 µM of **3c** (**Fig. 4H**). Further probing of this using increased excess of **3c** was prohibited by both solubility and viability limitations.

Taken together the data support that ALP-1 is a monovalent degrader of ULK1 that induces the formation of soluble ULK1 puncta which are cleared via ubiquitin dependent lysosomal degradation.

### ALP-1 engages the ULK1 Kinase Domain via Chloroacetamide Mediated Labelling of Asp102

Intrigued by the control experiment that showed the non-electrophilic ALP-1 control molecule **3c** could not out-compete ALP-1 at equivalent concentrations in cells, we looked to further characterise ALP-1:ULK1 engagement. To begin, we measured the interaction between ALP-1 and its controls with full-length ULK1 in a live cell nanoBRET target engagement assay. In this assay a fluorescent kinase-binding tracer is displaced by test compounds, leading to a decrease in BRET between the tracer and NanoLuc upon compound binding, and therefore a loss of signal. To our surprise, ALP-1 was shown to be >30-fold more potent than **3c**. As expected, N-methylated **3a** showed no engagement of ULK1 below 20 µM (**Fig. 5A, Fig S5A**).

**Figure 5.**
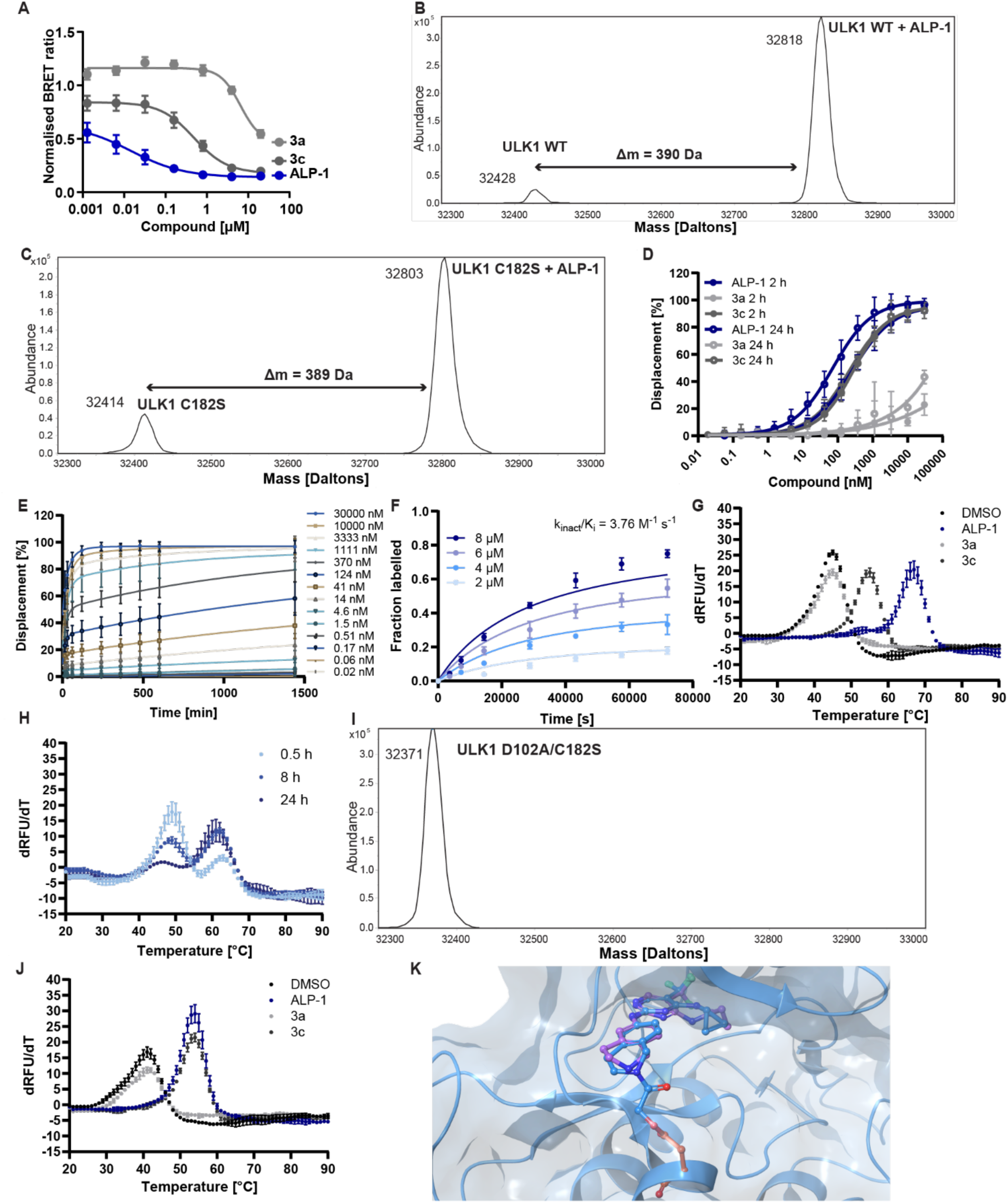
Characterising covalent engagement of ULK1 by ALP-1. (**A**) NanoBRET based ALP-1 target engagement profile in live cell mode in NanoLuc-ULK1 overexpressed HEK293 cells. Inhibition curves were generated with non-linear 4 parameter fit correction (Inhibition curves are means of n=3 biological replicates, ± SD). (**B**) Covalent modification of wild-type (WT) ULK1 kinase domain and (**C**) C182S ULK1 kinase domain by ALP-1. n=3 biological experiments performed, respective panels are representatives of one experiment. (**D**) Displacement of fluorescent tracer bound to ULK1 kinase domain in a fluorescence polarisation assay after incubation with ALP-1, **3a**, or **3c** after 2 h and 24 h incubation (the curves represent mean values of n=4 biological replicates ± SD). (**E)** Kinetic measurement of displacement of fluorescent tracer bound to the ULK1 kinase domain by ALP-1 measured by fluorescence polarisation (the curves represent mean values of n=4 biological replicates ± SD). (**F**) Kinetics of ALP-1 binding to ULK1 kinase domain measured by intact mass spectrometry (the curves represent mean values of n=3 biological replicates ± SD). (**G**) The effect of ALP-1, **3a**, and **3c** on thermal stability of ULK1 kinase domain measured by differential scanning fluorimetry (n=3 biological experiments performed, the panel is a representative of one experiment). (**H**) ALP-1-induced changes of thermal stability of ULK1 kinase domain measured by differential scanning fluorimetry at different incubation times (n=3 biological experiments performed, the panel is a representative of one experiment). (**I**) Intact mass spectrometry of ULK1 D102A/C182S with ALP-1 showing lack of labelled adduct (n=3 biological experiments performed, the panel is a representative of one experiment). (**J**) The effect of ALP-1, **3a**, and **3c** on thermal stability of D102A/C182S ULK1 kinase domain measured by differential scanning fluorimetry (n=3 biological experiments performed, the panel is a representative of one experiment). (**K**) Overlay of the unconjugated crystal structure of ALP-1 (purple) with ULK1 and ALP-1 (blue) covalently docked to D102 (orange) of ULK1.

The enhanced engagement affinity and the dependency of this effect on the presence of the electrophilic warhead led us to question whether ALP-1 covalently labels ULK1 itself in *cis*. To address this, we incubated recombinant ULK1 kinase domain with ALP-1 and **3a**, then measured the mass shift by intact protein mass spectrometry. We observed that ALP-1 covalently labels wild-type ULK1 (**Fig. 5B**) but also observed partial labelling with **3a** (**Fig. S5B, C**). It has previously been reported that C182 on the ULK1 activation loop engages cysteine reactive fragments^40^ and so we queried if this cysteine residue was to some extent hyperactive *in vitro*, potentially masking other binding events. We therefore re-tested both ALP-1 and **3a** with a C182S ULK1 mutant. In this case negative control **3a** showed no conjugation to ULK1 **(Fig. S5D, E)**, but ALP-1 retained its covalent engagement (**Fig. 5C**). This confirmed that ALP-1 covalently engages ULK1, and that both electrophile and hinge-binding motifs are required for this interaction. This also excluded C182 as the residue targeted by ALP-1.

We next sought to further characterise the covalent interaction. We established a fluorescence polarisation (FP) assay and observed that the IC50 of ALP-1 decreased over time from 250 nM after a 2 h incubation to 72 nM after 24 h, while IC50 values of **3c** stayed approximately the same (210 nM at 2 h vs. 280 nM at 24 h). We did not detect significant binding of **3a** at concentrations used in the assay (**Fig. 5D**). When analysing tracer displacement over time, we observed a biphasic curve characteristic of time dependent covalent interactions (**Fig. 5E**), which suggested that ALP-1 occupies the same ATP binding pocket as the tracer. However, due to slow tracer displacement by both ALP-1 and **3c** (**Fig. S6A**), we found it challenging to deconvolute the kinact/Ki values from the FP results. Instead, we measured the labelling kinetics via intact mass spectrometry^41^ (**Fig. 5F**). We observed slow engagement of ULK1 by ALP-1 with kinact/Ki = 3.76 M^-1^ s^-1^. We also measured the effect of ALP-1 binding on thermal stability of ULK1 kinase domain using differential scanning fluorimetry (DSF). The reversibly bound control **3c** produced a single transition with 9 °C shift in kinase domain stability. With ALP-1, however, we observed two phase transitions: one at the same temperature as **3c** and then a second shift 22 °C higher than the Tm of DMSO or **3a** treated ULK1 (**Fig. 5G**). The proportion of the observed peaks corresponding to the phase transitions changed over time, with the increase in the peak corresponding to the second phase transition, suggesting that it corresponded to the covalent engagement of ULK1 (**Fig. 5H**).

These observations led us to two alternative theories: either ALP-1 binds to a secondary binding site outside of the ATP binding pocket, or it engages a non-cysteine residue. Introducing C47A/C182S mutations to the ULK1 kinase domain did not abolish covalent ALP-1 binding (**Fig. S6B-D**). Since these are the only solvent-exposed cysteine residues in the kinase domain, we attempted to identify the covalently bound residue of ULK1 by X-ray crystallography and unbiased mass spectrometry. We obtained a co-crystal structure of ULK1 kinase domain with ALP-1, showing an expected binding mode in the ATP site, however, with no continuous electron density around the electrophilic warhead. We did note that the THIQ pointed towards an aspartate residue, D102, though were not aware of any precedence for chloroacetamides being reactive with acidic residues (**Fig. S6E, F**). In parallel, we attempted identifying the covalently labelled residue via unbiased mass spectrometry. Remarkably, the analysis of the peptides resulting from both trypsin and AspN digestion supported labelling of the D102 residue (**Fig. S7, S8**). To validate this observation, we expressed and purified an ULK1 kinase domain construct with a dual D102A/C182S mutation. This point mutant was not covalently labelled when tested by intact mass spectrometry (**Fig. 5I**). Similarly, DSF measurement showed that ALP-1 has the same effect on protein stability as non-electrophilic control **3c** with this point mutant. This further supported covalent D102 engagement and that this covalent engagement is responsible for the additional thermal stabilisation (**Fig. 5J**). Taken together, the data support that the carboxylate anion of D102 is able to attack the chloroacetamide warhead, displacing the chloride and forming an ester conjugate of D102 (**Fig. 5K**). The slow rate of labelling suggests this is not an intrinsically reactive combination, as may be expected, with the optimal geometric positioning of the warhead proximal to D102 likely crucial.

## Discussion

In this study we developed electrophilic tool compounds, we termed scout molecules, to identify kinases that may be degraded in cells as starting points for targeted probe discovery. We minimally decorated a promiscuous kinase ATP-competitive inhibitor with reactive warheads for a range of nucleophilic amino acids. Our initial exploration of a focussed set of kinases via western blot. Alongside this, unbiased proteome-wide mass spectrometry, nominated a small set of hit kinases, whose abundance was lowered upon scout molecule treatment and which provide potential rational starting points for multiple future kinase degradation programmes. This led us to the key finding that a chloroacetamide warhead, a commonly employed cysteine electrophile, can in fact label a solvent exposed, acidic amino acid residue in addition to providing a lead molecule for monovalent ULK1 degradation, ALP1.

Profiling of our scout molecules showed the overall rate for finding degrader hit kinases was relatively low, even across an extended panel of probes. Nevertheless, when accounting for observations in our cycloheximide control experiment, where possible, we were able to identify RIOK2, LZK and ULK1 as promising degrader starting points. This aligned with recent results by Mozes et al., who disclosed their own learnings from use of electrophilic kinase probes^42^, including the same chloroacetamide and acrylamide analogues (herein named as KinoGlu1 and 2) that we also profiled here. They identifed AURKA as a stabilisation hit, before demonstrating that introducing a linker between the ligand and the electrophile flipped this to become an AURKA degrader. Reinspection of our own data showed that a trend towards AURKA stabilisation with KinoGlu 1 can be seen, though it did not meet our cut-offs for hit nomination. It should be noted that cut-offs, doses and treatment times were not always identical between the studies.

Due to the complex and critical role played by ULK1 in autophagy initiation, ULK1 inhibitors have garnered significant attention and recently the first clinical phase inhibitors were disclosed and are being profiled against a range of solid tumour types^21^. Covalent fragments and inhibitors of ULK1, targeting activation loop C182, have been reported^40, 43^, though are of significantly lower affinity than known reversible inhibitors, such as the compounds progressed to the clinic. Additionally, methods to chemically control autophagy initiation are of interest to study a broad range of disease settings and pathways, for example in the context of neurodegenerative diseases where deficiencies in autophagic processes are well documented^44, 45^. Chemical tools with novel mechanisms of action to probe autophagy regulators are therefore highly sought and we were interested to test if KinoGlu 1, which had revealed ULK1 as a possible degrader hit, could be modified to generate a chemical tool with a differentiated mechanism of action versus classical reversible inhibitors. Changing the initial multi-kinase scaffold of TL13-87 for a scaffold based on ULK1 inhibitor SBP-7455 yielded ALP-1, which degrades ULK1 with moderate potency across multiple cell lines. Recently, a study focussed on optimising the SBP-7455 core, identified the more potent ULK1 inhibitor, SBP-5147^23^. This molecule contains an unsubstituted THIQ analogous to the design of our own compounds, though in our case we include further substitution of the amino group, rendering this non-basic. Consistent with SAR from that study, this difference of amide for amine may reduce potency as an inhibitor in the case of our control compound **3c**, by preventing reversible hydrogen-bonding interactions with D102 on the lip of the ATP-site. However, in the case of ALP-1, the chloroacetamide switches the function of the compound to be a degrader as well as a more potent ULK1 engager. Based on our proteomics data, the chloroacetamide warhead itself would likely require optimisation to convert this promising chemical tool into a selective ULK1 degrader chemical probe, highlighting the opportunities of this warhead for scouting labelling events, but also the limitations for discovery of highly specific probes. Nevertheless, consistent with its ability to potently engage ULK1 and degrade it, ALP-1 shows a marked improvement in shutdown of ULK1 specific ATG14 phosphorylation and autophagy induction, as compared with its non-electrophilic molecular matched pair, underlining the potential of this approach.

Mechanism of action studies highlighted both lysosome and E1, but not proteasome, dependency. Whilst IP-MS did not highlight any clear candidates for further follow-up, it is also plausible that unoptimised compounds such as ALP-1 may act via multiple mechanisms, clouding the impact of individual mechanistic contributions, and these profiles may shift during optimisation^26, 46^. Given that methylation of the hinge binder (**3a**) and removal of the electrophile (**3c**) shut down most activity in both degradation and target engagement assays, we were intrigued by the observation that competition with the non-electrophilic control at equivalent concentrations could not appreciably rescue degradation of ULK1. This control experiment is rarely observed in monovalent degrader discovery studies but is of significant value. In our case, combined with our observation of vastly improved ULK1 cellular engagement, this prompted us to question if we were labelling ULK1 itself in *cis*, rather than labelling a secondary effector protein in *trans.* This was in the context of considering previous work using electrophilic kinase inhibitors and tools to label residues outside of the canonical binding site^18, 19^.

Whilst efforts to target nucleophilic amino acids beyond cysteine have advanced, carboxylic acid side chains have remained a significant challenge given their weak nucleophilicity and tendency towards solvent exposure and/or involvement in molecular interactions^47^. The tractability of aspartate as a covalent target has gained renewed attention, predominantly through efforts to target the KRAS G12D mutation, prompting the development of specialised warheads^48, 49^ and template assisted covalent inhibition approaches using chaperone proteins^50^. Our data supports that the chloroacetamide of ALP-1 can label the ULK1 kinase domain, engaging D102 of the D-helix, representing a rare case of covalently targeting an aspartic acid residue. The slow rate of ULK1 labelling shows that the chloroacetamide of ALP-1, although able to form an adduct, is likely a sub-optimal electrophile for D102 engagement. It also, however, raises an important consideration as chloroacetamide is so commonly employed as a cysteine targeted warhead, yet there appears to be unappreciated reactivity beyond cysteine. We believe this represents a valuable starting point for further development of covalent kinase targeted molecules and one which may be broadly applicable given the preponderance of acidic residues at the lip of ATP binding sites^51^.

## Conclusion

In summary, we present an approach to systematically identify chemical leads for modulating target proteins beyond classical reversible inhibition. This unbiased approach has led us to discoveries that would not have been prospectively and rationally envisaged, most notably the observation that a chloroacetamide warhead can conjugate an acidic amino acid residue and that the high-profile drug target ULK1 is targetable for monovalent degrader approaches. Mechanistic studies will continue to discern the relationship between covalent engagement and degradation of ULK1 to inform on future optimisation efforts. However, this study highlights a potentially underexplored opportunity to utilise surface exposed acidic residues that are proximal to established atp-binding pockets, as a means for covalent kinase targeting.

## Acknowledgements

This project received funding from UK Engineering and Physical Sciences Research Council (EPSRC) grant number EP/X020088/1 (W.F., N.M.K.) and by the Wellcome Trust Award 226943/Z/23/Z (W.F., M.I.). A.L.P. is supported by an EN & MN Lindsay Scholarship. Q.A. is supported by the Wellcome Trust. I.G.G. was funded by the Medical Research Council, UK (MC_UU_00038/2). We are thankful to Dr Conner Craigon for guide

RNA design and the University of Dundee FACs facility for single cell sorting during HiBiT Knock-In cell line generation; the University of Dundee Proteomics facility for assisting with MS-proteomics data analysis, MRC Reagents and Services including support with generating constructs used in this study and amplicon sequencing. We acknowledge Diamond Light Source for time on beamline i24 under proposal MX41060, beamline session MX41060-27. We thank Dr Vesna Vetma, Dr Luke Simpson and Dr Ilaria Puoti for support and training with early chemoproteomics studies and western blotting assays. We are thankful to Dr Michael Krummhaar for his help with data acquisition for crystallographic experiment.

## Author Contributions

A.L.P. and N.M.K. contributed equally and will be putting their name first on the citation in their curriculum vitae. W.F. conceived the idea, acquired research funds and directed the project. I.G. and A.R. provided input to experimental design. W.F. and A.L.P. designed compounds. A.L.P. and W.F. performed synthesis of compounds.

N.M.K. and M.I. generated and validated cell lines. A.L.P., N.M.K. and Q.A. performed cellular profiling experiments. M.I. purified proteins and designed and performed intact MS, DSF, and crystallographic experiments. M.I. and N.M.K. performed FP experiments. L.W. performed autophagy assays. A.R. performed microscopy experiments. N.M.K. and G.S. designed chemoproteomics studies, performed sample preparation and mass spectrometry experiments, including related data processing and analysis. A.L.P., W.F., N.M.K. and M.I. wrote the manuscript with input from all other authors.

## Lead Contact

Further information, questions and requests for materials and reagents should be directed to Dr William Farnaby. ORCID ID: 0000-0001-8610-932X.

## Supplementary Information

### Supplementary Figures and Data

**Figure S1.**
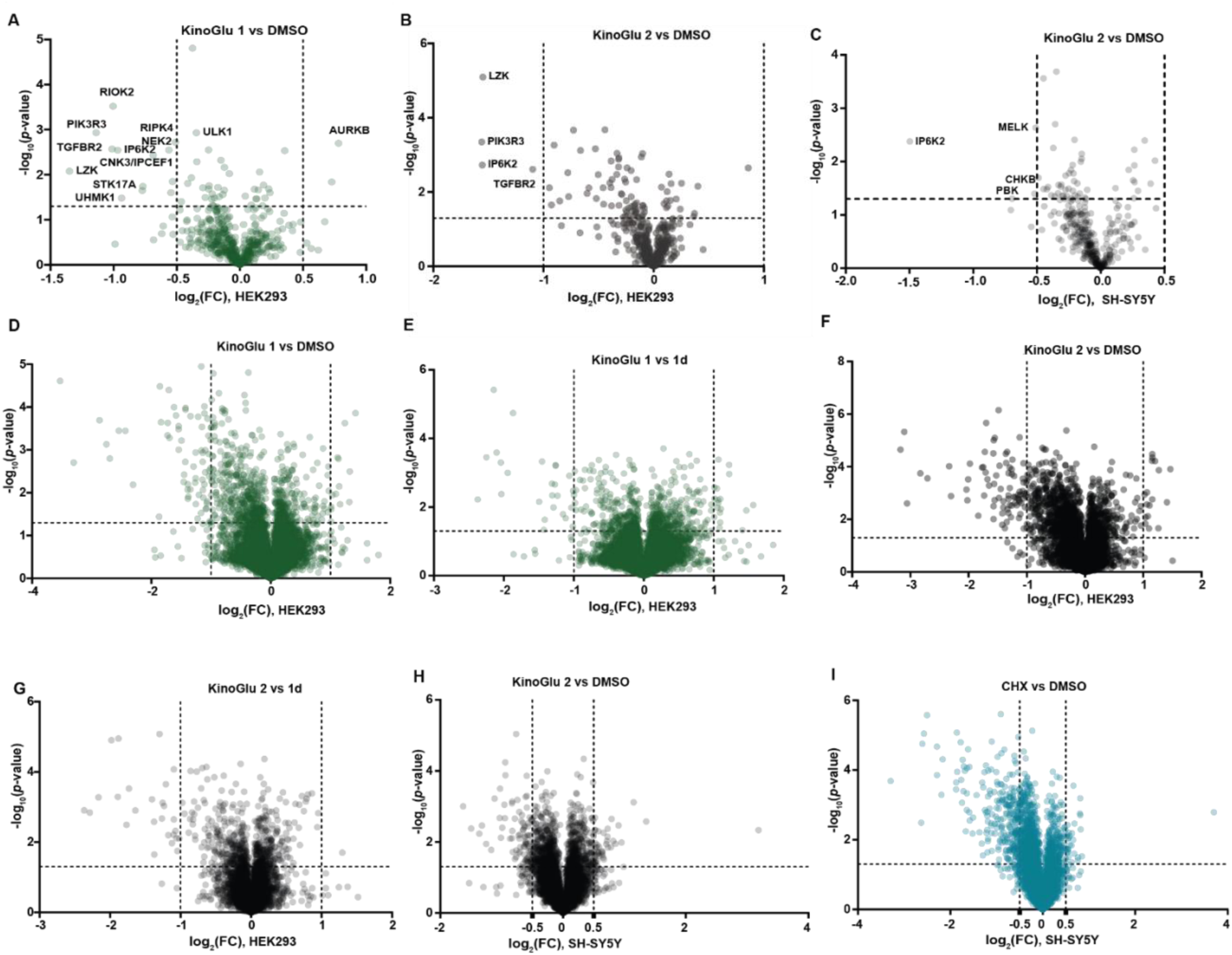
Proteomics profiling of KinoGlu 1 and KinoGlu 2. Kinome-wide abundance profiling using (**A**) KinoGlu 1 (1 µM, 4 h) and (**B**) KinoGlu 2 (1 µM, 4 h) normalised to DMSO in HEK293 cells and (**C**) KinoGlu 2 (1 µM, 4 h) in SH-SY5Y cells (at least n=3 biological replicates). Changes in total proteome in response to KinoGlu 1 treatment (1 µM, 4 h) compared to negative controls (**D**) DMSO and (**E**) **1d** in HEK293 cells. Changes in total proteome in response to KinoGlu 2 treatment (1 µM, 4 h) compared to negative controls (**F**) DMSO and (**G**) **1d** in HEK293 cells (n=3 biological replicates). Changes in total proteome in response to (**H**) KinoGlu 2 treatment (1 µM, 4 h) and (**I**) cycloheximide (CHX, 100 ug/ml, 4 h) compared to negative control DMSO in SH-SY5Y cells (at least n=3 biological replicates).

**Figure S2.**
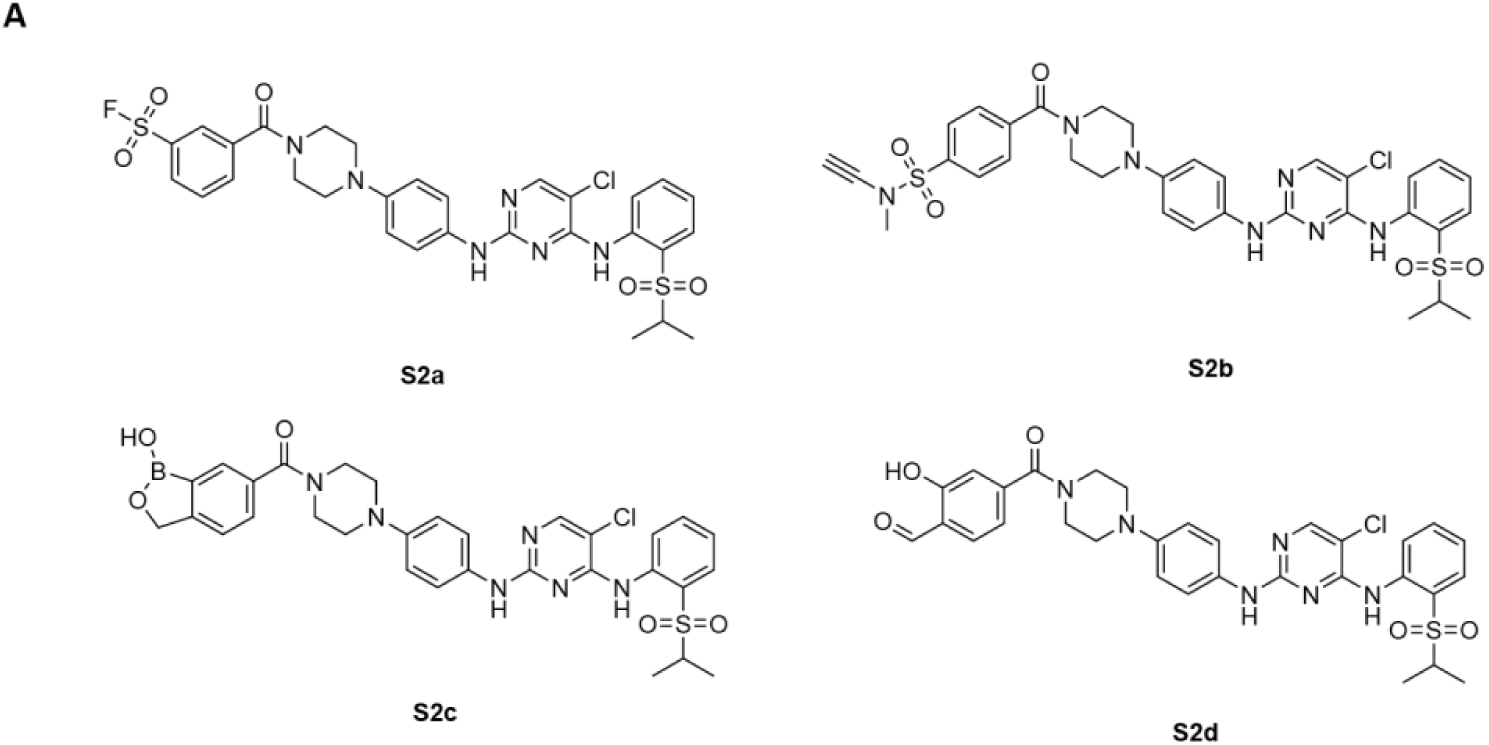
Chemical structures of beyond-cysteine targeting KinoGlus. **(A) S2a (sulfonyl fluoride), S2b** (ynamide), **S2c** (benzoxaborole) and **S2d** (salicaldehyde).

**Figure S3.**
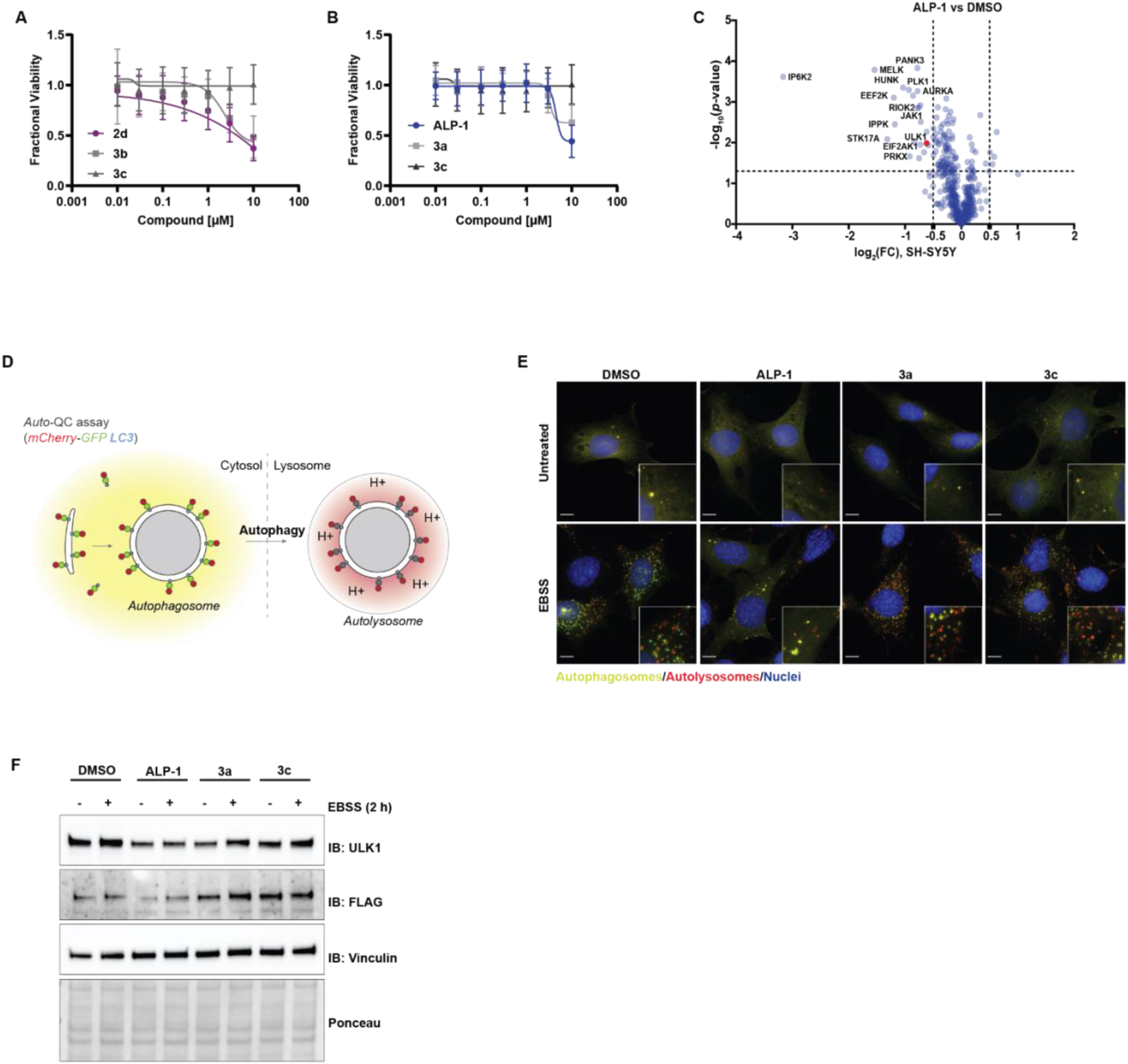
Effect of ULK1-targeted degraders on viability, proteome and downstream signalling. Cell viability of ULK1-HiBiT KI HEK293 cells upon dose-dependent (**A**) **2d**, **3b** and **3c** treatment (8 h) (**B**) ALP-1, **3a** and **3c** treatment (8 h). Viability curves were generated with non-linear 4 parameter fit correction (Curves are means of n=3 biological replicates, ± SD). (**C**) Proteome-wide activity profiling of ALP-1 normalised to DMSO on kinase level in SH-SY5Y cells (n=3 biological replicates, 3 µM, 8 h). (**D**) Schematic of the Auto-QC assay. (**E**-**F**) ULK1/2 double-knockout (DKO) MEFs stably expressing FLAG-tagged ULK1 and the Auto-QC reporter were pretreated with ALP-1, **3a** or **3c** (1 µM, 4 h), followed by 2 h of combined compound and EBSS treatment. (**E**) Representative wide-field images and (**F**) representative immunoblots of the indicated proteins.

**Figure S4.**
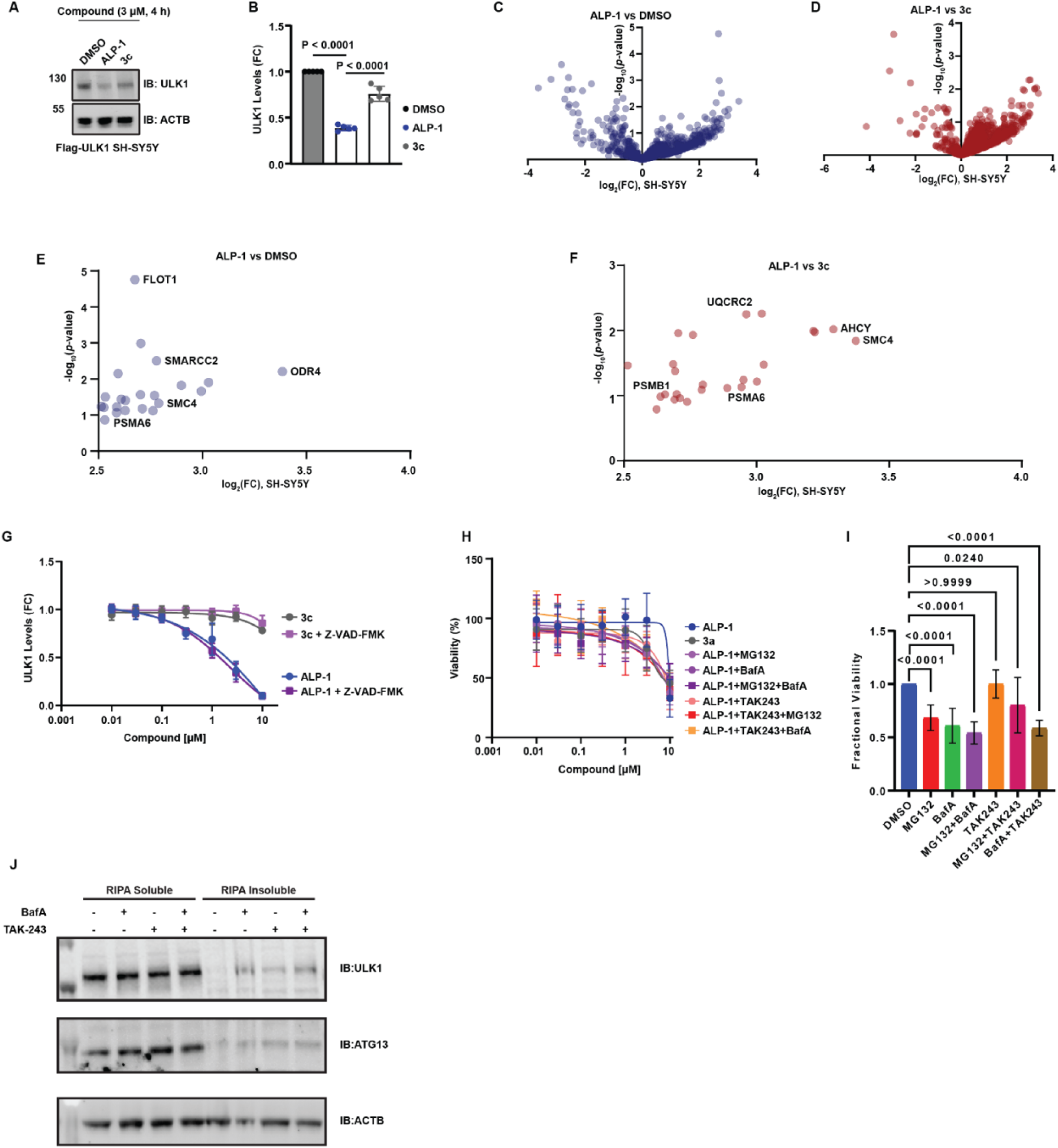
Mode-of-action studies of ALP-1. (**A**) Immunoblotting based confirmation of the effect of ALP-1 and **3c** (3 µM, 4 h) in stably Flag-ULK1 expressing SH-SY5Y cells used for IP-MS experiments. Representative blots of n=5 biological replicates. (**B**) Relative quantification of ULK1 levels normalised to internal controls in stably Flag-ULK1 expressing SH-SY5Y cells following treatments with ALP-1 and **3c** (3 µM, 4 h) (graph represents n=5 biological replicates, mean, ±SD). p values were calculated as indicated in the panel by using unpaired t-test (GraphPad Prism, v10.6.0(890)). Changes in ULK1 interactome upon ALP-1 (3 µM, 4 h) treatment compared to (**C**) DMSO and (**D**) **3c**. ALP-1 mediated enhanced protein-protein interactions with ULK1 compared to (**E**) DMSO and (**F**) **3c** (n=5 biological replicates). (**G**) ALP-1 and **3c** (10, 3, 1, 0.3 and 0.1 µM) in combination with Z-VAD-FMK (40 µM) in ULK1-HiBiT KI HEK293 cells. Dose-dependent degradation curves were generated with non-linear 4 parameter fit correction and representative graphs are combined results of n=3 biological replicates. (Dose response curves are means of n=3 biological replicates, ± SD). (**H**) The effect of ALP-1 (10, 3, 1, 0.3 and 0.1 µM) in combination with MG132 (1 µM), Bafilomycin A (BafA1, 100 nM) and TAK-243 (1 µM) alone or co-treatments with indicated concentrations; MG132 and BafA1, TAK-243 and MG132, TAK-243 and BafA. ULK1-HiBiT KI HEK293 cells pre-treated with aforementioned inhibitors for 2 h and then co-treated with ALP-1 for further 8 h. Viability curves were generated with non-linear 4 parameter fit correction and representative graphs are combined results of n=3 biological replicates, ± SD. (**I**) The effect of inhibitors alone on cell viability in ULK1-HiBiT KI HEK293 cells. Graph is representative of n=3 biological replicates, p values are calculated as indicated in the panel compared to DMSO using unpaired t-test (GraphPad Prism). (**J**) Western blot of ULK1 and ATG13 levels in the RIPA soluble and insoluble fractions of SHSY-5Y cells upon 8 h treatment with BafA1 (100 nM), TAK-243 (1 µM), or a combination thereof. Blots representative of n=3 biological replicates.

**Figure S5.**
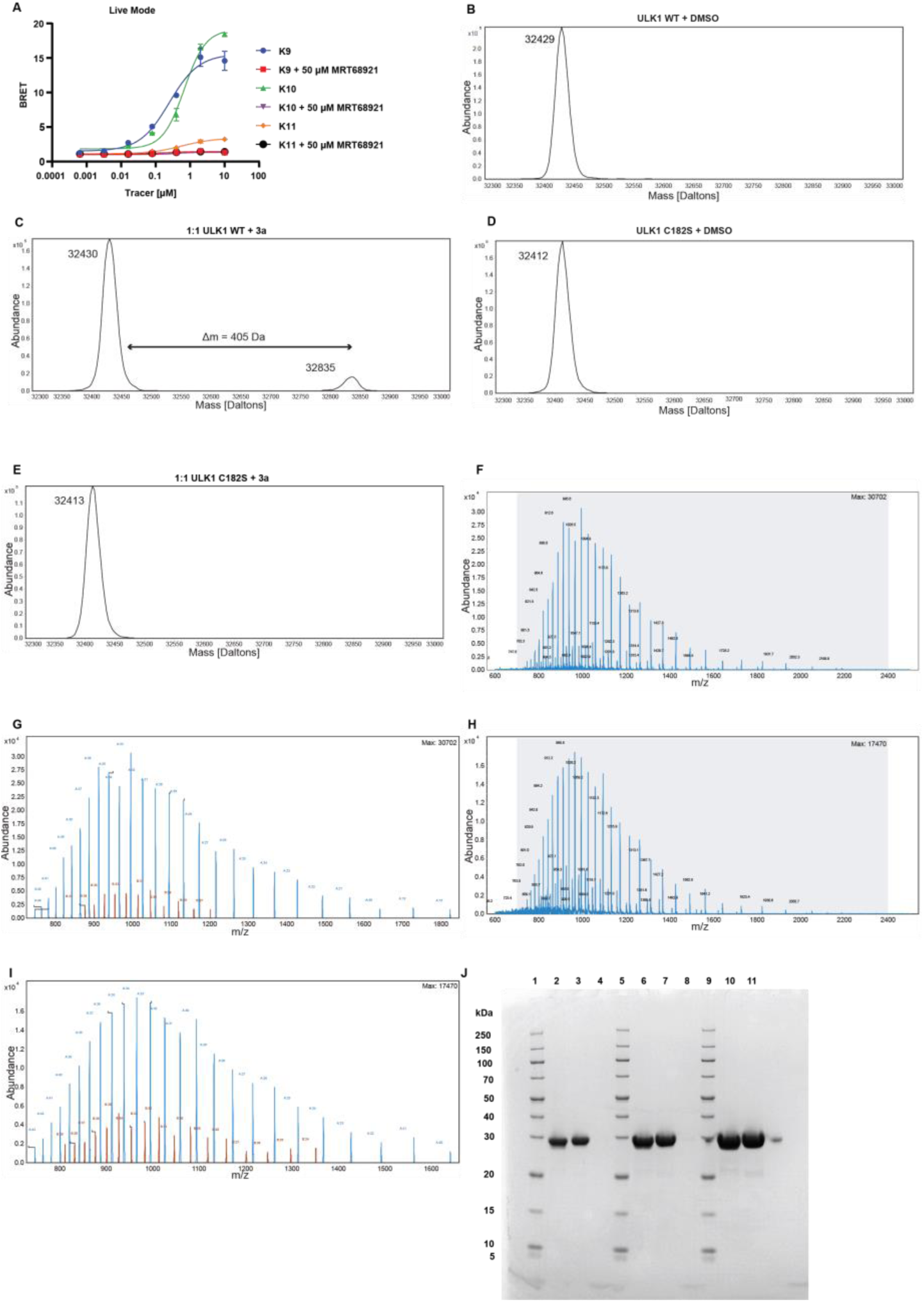
Intact MS labelling experiments using wild type (WT) and C182S ULK1 kinase domain. (**A**) Tracer optimisation for NanoLuc-ULK1 in live cells. NanoLuc-ULK1 overexpressed HEK293 cells incubated with tracers (K9, K10 and K11, Promega) alone or with excess amount of ULK1 inhibitor (MRT68921, 50 µM) for 2 h, donor and acceptor signals measured to calculate BRET ratios. Inhibition curves were generated with non-linear 4 parameter fit correction (Curves are representative of one experiment from 3 biological replicates). (**B**-**E**) The plots represent deconvoluted mass of protein incubated at 10 µM concentration at room temperature (**B**, **C**) DMSO (2%, 18 h) or (**D**, **E**) **3a** (10 µM, 18 h). m/z envelopes used for deconvolution of protein masses in Figure 4H and I. (**F**) Recorded m/z envelope of wild type ULK1 kinase domain labelled with ALP-1 (**G**) m/z peaks used for deconvolution of ALP-1-labelled (blue) and unlabelled (red) wild type ULK1; (**H**) Recorded m/z envelope of ULK1 C182S kinase domain labelled with ALP-1 (**I**) m/z peaks used for deconvolution of ALP-1-labelled (blue) and unlabelled (red) ULK1 C182S. (**J**) SDS PAGE gel with recombinant ULK1 kinase domain samples stained with InstantBlue Coomassie Protein Stain (Abcam, #ab119211). 1. PageRuler™ Unstained Broad Range Protein Ladder (Thermo Fisher Scientific, #26630); 2. 1 µg ULK1(1-283) WT; 3. 1 µg ULK1(1-283) C182S; 4. empty lane; 5. PageRuler™ Unstained Broad Range Protein Ladder; 6. 2 µg ULK1(1-283) WT; 7. 2 µg ULK1(1-283) C182S; 8. Empty lane; 9. PageRuler™ Unstained Broad Range Protein Ladder; 10. 4 µg ULK1(1-283) WT; 11. 4 µg ULK1(1-283) C182S.

**Figure S6.**
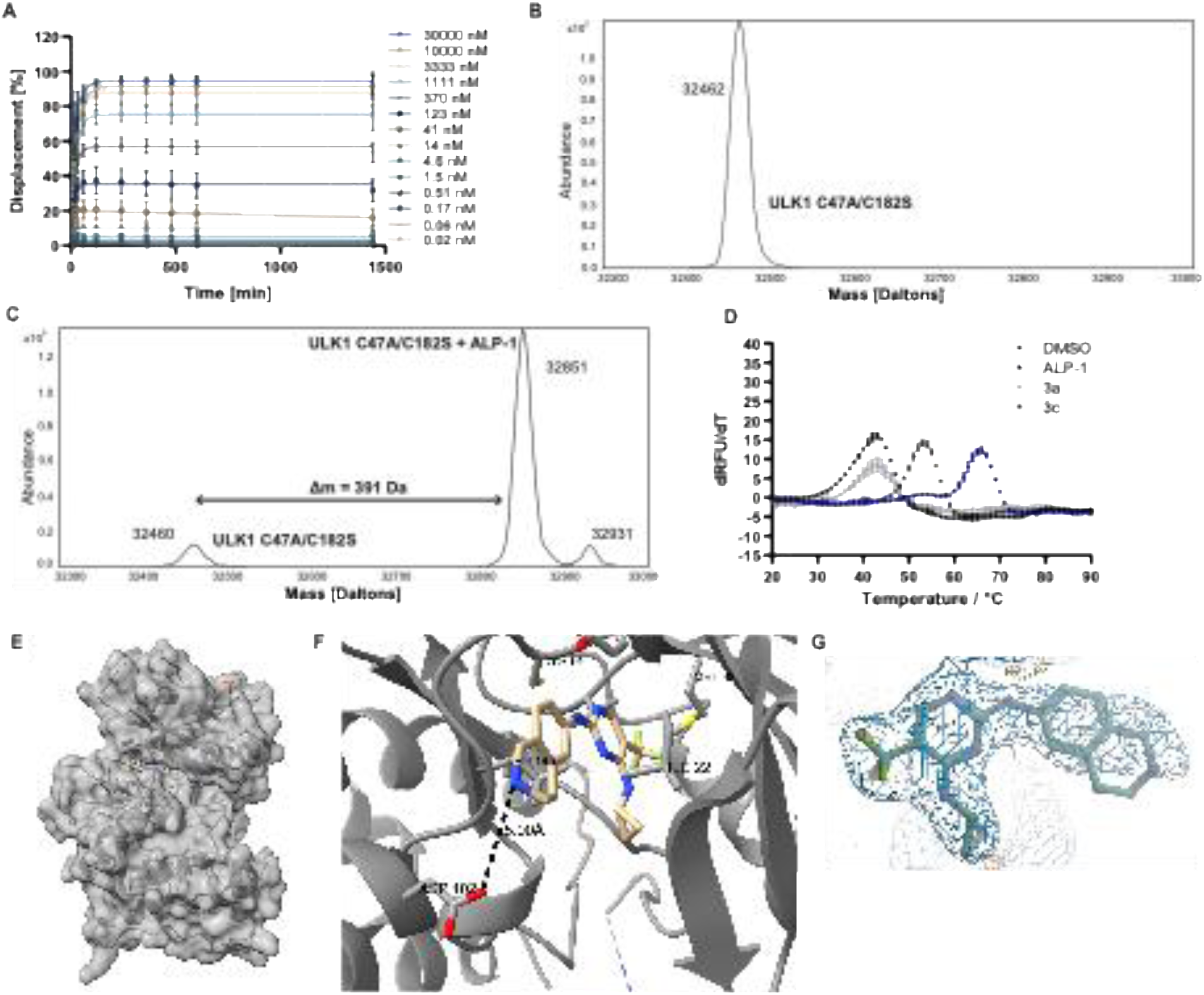
Validation of ALP-1 labelled residue. (**A**) Kinetic measurement of displacement of fluorescent tracer bound to the ULK1 kinase domain by **3c** measured by fluorescence polarisation (the curves represent mean values of n=4 biological replicates ± SD). (**B**) Intact mass spectrometry of ULK1 C47A/C182S after 18 h incubation with 2% DMSO (n=3 biological experiments performed, the panel is a representative of one experiment). (**C**) Intact mass spectrometry of ULK1 C47A/C182S after 18 h incubation with ALP-1 showing formation of labelled adduct (n=3 biological experiments performed, the panel is a representative of one experiment). (**D**) The effect of ALP-1, **3a**, and **3c** on thermal stability of C47A/C182S ULK1 kinase domain measured by differential scanning fluorimetry after 18 h of incubation with the compounds (n=3 biological experiments performed, the panel is a representative of one experiment). (**E**) An overview of the crystal structure of ULK1 C182S kinase domain in complex with ALP-1. (**F**) A close-up view of ALP-1 bound to ATP binding pocket of ULK1 KD C182S. The ALP-1 structure was truncated to match the visible electron density. (**G**) A close-up view of the ALP-1 electron density in the co-crystal structure of ULK1 complex with ALP-1.

**Figure S7.**
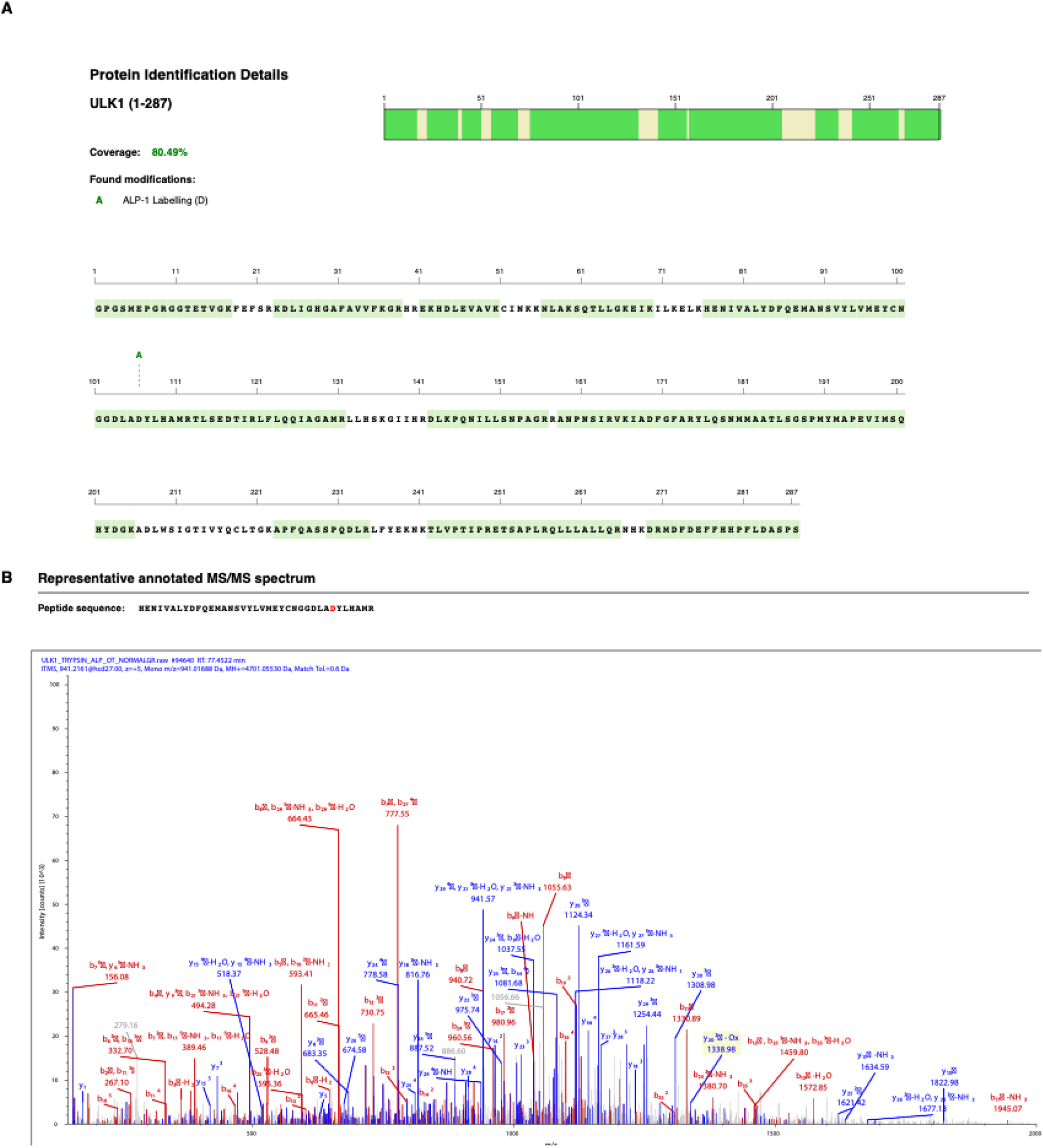
LC–MS/MS identification of ALP-1-labelled ULK1 sites following trypsin digestion. (**A**) ULK1 sequence coverage and identified post-translational modifications. ALP-1 labelling was detected at Asp102, it appears as Asp106 in the figure because of the addition of four amino acids (GPGS) at the N-terminus. Overall protein sequence coverage is 80.49%. (**B**) Representative annotated MS/MS spectrum of the ALP-1-labelled peptide, HENIVALYDFQEMANSVYLVMEYCNGGDLADYLHAMR. The ALP-1-labelled Asp102 residue is highlighted in red.

**Figure S8.**
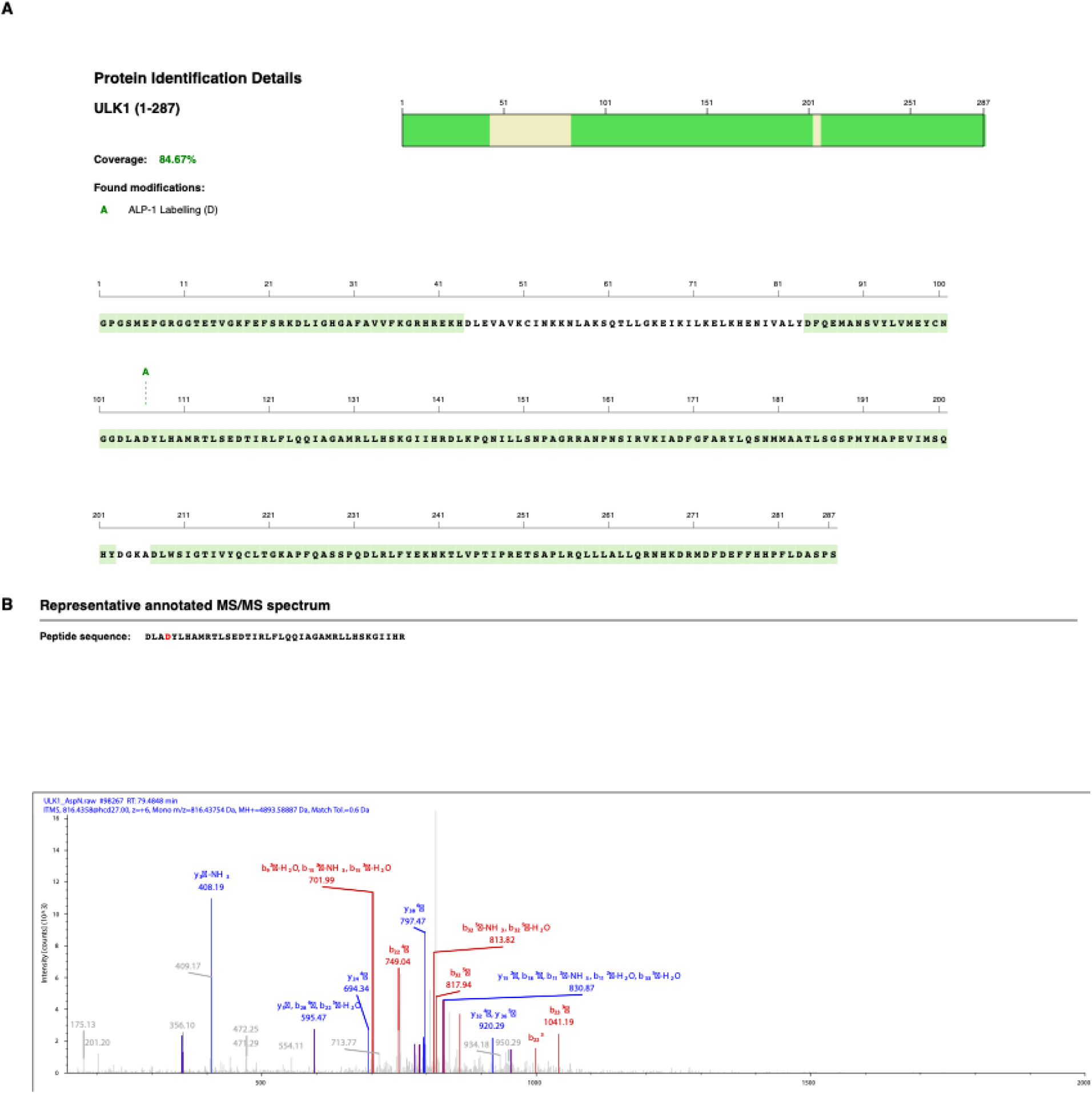
LC–MS/MS identification of ALP-1-labelled ULK1 sites following AspN digestion. (**A**) ULK1 sequence coverage and identified post-translational modifications. ALP-1 labelling was detected at Asp102; however, it appears as Asp106 in the figure because of the addition of four amino acids (GPGS) at the N-terminus. The overall protein sequence coverage was 84.67%. (**B**) Representative annotated MS/MS spectrum of the ALP-1-labelled peptide, DLADYLHAMRTLSEDTIRLFLQQIAGAMRLLHSKGIIHR. The ALP-1-labelled Asp102 residue is highlighted in red.

**Table S1.** Statistic parameters for crystallographic data collection and refinement.

| **Data collection** |  |
|---|---|
| Wavelength | 0.6199 |
| Resolution range | 53.02 - 2.288 (2.39 - 2.29) |
| Space group | P 1 21 1 |
| Unit cell dimensions |  |
| a, b, c (Å) | 47.19, 64.62, 93.88 |
| α, β, γ (°) | 90.00, 98.93, 90.00 |
| Total reflections | 159622 (21106) |
| Unique reflections | 24165 (3129) |
| Multiplicity | 6.6 (6.7) |
| Completeness (%) | 93.23 (85.29) |
| Mean I/sigma(I) | 8.14 (0.78) |
| Wilson B-factor | 63.33 |
| R-merge | 0.08395 (2.431) |
| R-meas | 0.09135 (2.635) |
| R-pim | 0.0356 (1.01) |
| CC1/2 | 0.998 (0.449) |
| CC* | 1 (0.787) |
| **Refinement** |  |
| Reflections used in refinement | 23655 (2668) |
| Reflections used for R-free | 1173 (149) |
| R-work | 0.226 (0.399) |
| R-free | 0.283 (0.461) |
| Number of non-hydrogen atoms | 4264 |
| macromolecules | 4212 |
| ligands | 50 |
| solvent | 2 |
| Protein residues | 517 |
| RMS Deviations |  |
| Bonds (Å) | 0.008 |
| Angles (°) | 0.88 |
| Ramachandran favored (%) | 96.79 |
| Ramachandran allowed (%) | 3.21 |
| Ramachandran outliers (%) | 0 |
| Rotamer outliers (%) | 1.77 |
| Clashscore | 7.01 |
| Average B-factor | 87.55 |
| macromolecules | 87.59 |
| ligands | 85.68 |
| solvent | 58.09 |

**Table S2.**
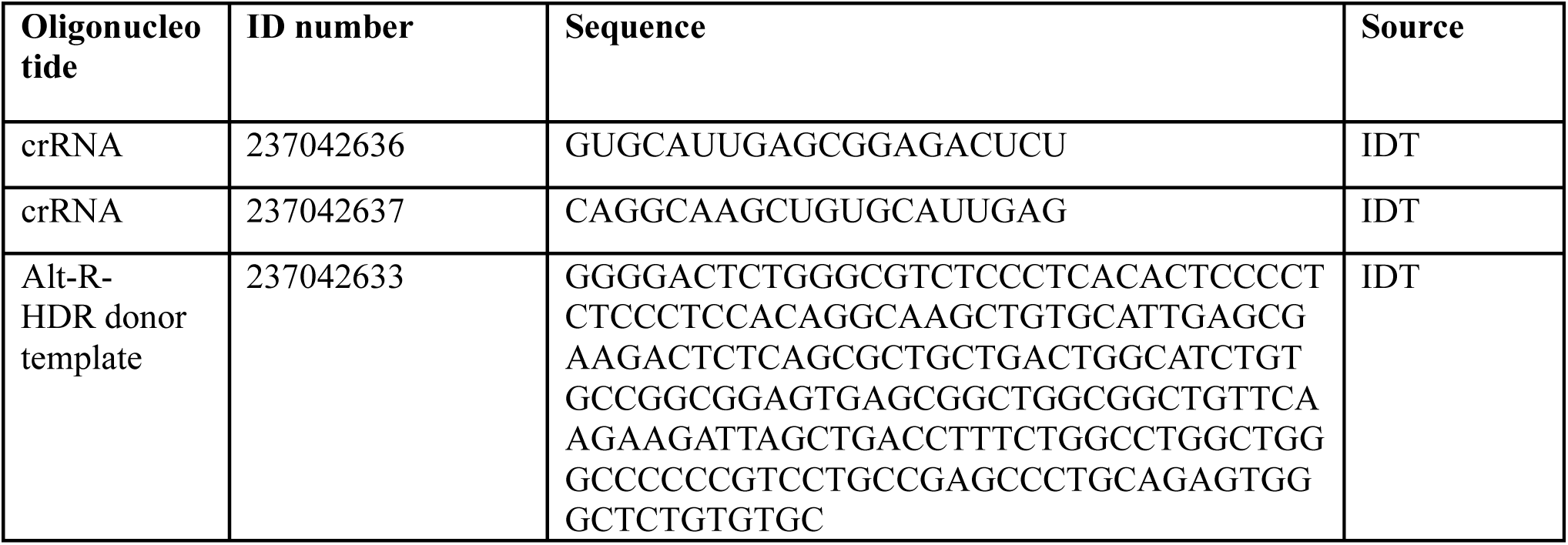
Oligonucleotides used for generation of CRISPR HiBiT Knock-In cell line.

**Table S3.** The list of the plasmids used for recombinant protein expression and their open reading frames encoding ULK1 kinase domain.

| Plasmid | Protein sequence | Source |
|---|---|---|
| pET28a-6xHis-TEV-T7 tag-AviTag-PreScission-ULK1(1-283) | MGSSHHHHHHSSGENLYFQGHMASMTGGQQMGRGSGLNDIFEAQKIEWHEGSGLEVLFQGPGSMEPGRGGTETVGKFEFSRKDLIGHGAFAVVFKGRHREKHDLEVAVKCINKKNLAKSQTLLGKEIKILKELKHENIVALYDFQEMANSVYLVMEYCNGGDLADYLHAMRTLSEDTIRLFLQQIAGAMRLLHSKGIIHRDLKPQNILLSNPAGRRANPNSIRVKIADFGFARYLQSNMMAATLCGSPMYMAPEVIMSQHYDGKADLWSIGTIVYQCLTGKAPFQASSPQDLRLFYEKNKTLVPTIPRETSAPLRQLLLALLQRNHKDRMDFDEFFHHPFLDASPS | GeneScript |
| pET28a-6xHis-TEV-T7 tag-AviTag-PreScission-ULK1(1-283) C182S | MGSSHHHHHHSSGENLYFQGHMASMTGGQQMGRGSGLNDIFEAQKIEWHEGSGLEVLFQGPGSMEPGRGGTETVGKFEFSRKDLIGHGAFAVVFKGRHREKHDLEVAVKCINKKNLAKSQTLLGKEIKILKELKHENIVALYDFQEMANSVYLVMEYCNGGDLADYLHAMRTLSEDTIRLFLQQIAGAMRLLHSKGIIHRDLKPQNILLSNPAGRRANPNSIRVKIADFGFARYLQSNMMAATL**S**GSPMYMAPEVIMSQHYDGKADLWSIGTIVYQCLTGKAPFQASSPQDLRLFYEKNKTLVPTIPRETSAPLRQLLLALLQRNHKDRMDFDEFFHHPFLDASPS | This study |
| pET28a-6xHis-TEV-T7 tag-AviTag-PreScission-ULK1(1-283) C47A/C182S | MGSSHHHHHHSSGENLYFQGHMASMTGGQQMGRGSGLNDIFEAQKIEWHEGSGLEVLFQGPGSMEPGRGGTETVGKFEFSRKDLIGHGAFAVVFKGRHREKHDLEVAVK**A**INKKNLAKSQTLLGKEIKILKELKHENIVALYDFQEMANSVYLVMEYCNGGDLADYLHAMRTLSEDTIRLFLQQIAGAMRLLHSKGIIHRDLKPQNILLSNPAGRRANPNSIRVKIADFGFARYLQSNMMAATL**S**GSPMYMAPEVIMSQHYDGKADLWSIGTIVYQCLTGKAPFQASSPQDLRLFYEKNKTLVPTIPRETSAPLRQLLLALLQRNHKDRMDFDEFFHHPFLDASPS | This study |
| pET28a-6xHis-TEV-T7 tag-AviTag-PreScission-ULK1(1-283) D102A/C182S | MGSSHHHHHHSSGENLYFQGHMASMTGGQQMGRGSGLNDIFEAQKIEWHEGSGLEVLFQGPGSMEPGRGGTETVGKFEFSRKDLIGHGAFAVVFKGRHREKHDLEVAVKCINKKNLAKSQTLLGKEIKILKELKHENIVALYDFQEMANSVYLVMEYCNGGDLA**A**YLHAMRTLSEDTIRLFLQQIAGAMRLLHSKGIIHRDLKPQNILLSNPAGRRANPNSIRVKIADFGFARYLQSNMMAATL**S**GSPMYMAPEVIMSQHYDGKADLWSIGTIVYQCLTGKAPFQASSPQDLRLFYEKNKTLVPTIPRETSAPLRQLLLALLQRNHKDRMDFDEFFHHPFLDASPS | This study |
| pRSF-DUET-12xHis-TEV-AviTag-PreScission-ULK1(1-283) C47A/C182S | MGSSHHHHHHHHHHHHGSSMKIEEGKLVIWINGDKGYNGLAEVGKKFEKDTGIKVTVEHPDKLEEKFPQVAATGDGPDIIFWAHDRFGGYAQSGLLAEITPDKAFQDKLYPFTWDAVRYNGKLIAYPIAVEALSLIYNKDLLPNPPKTWEEIPALDKELKAKGKSALMFNLQEPYFTWPLIAADGGYAFKYENGKYDIKDVGVDNAGAKAGLTFLVDLIKNKHMNADTDYSIAEAAFNKGETAMTINGPWAWSNIDTSKVNYGVTVLPTFKGQPSKPFVGVLSAGINAASPNKELAKEFLENYLLTDEGLEAVNKDKPLGAVALKSYEEELAKDPRIAATMENAQKGEIMPNIPQMSAFWYAVRTAVINAASGRQTVDEALKDAQTNSSSNNNNNNNNNNLGIEENLYFQGSGLNDIFEAQKIEWHEGSGLEVLFQGPGSMEPGRGGTETVGKFEFSRKDLIGHGAFAVVFKGRHREKHDLEVAVK**A**INKKNLAKSQTLLGKEIKILKELKHENIVALYDFQEMANSVYLVMEYCNGGDLADYLHAMRTLSEDTIRLFLQQIAGAMRLLHSKGIIHRDLKPQNILLSNPAGRRANPNSIRVKIADFGFARYLQSNMMAATL**S**GSPMYMAPEVIMSQHYDGKADLWSIGTIVYQCLTGKAPFQASSPQDLRLFYEKNKTLVPTIPRETSAPLRQLLLALLQRNHKDRMDFDEFFHHPFLDASPS | This study. |

### Materials and Methods

#### Cell culture

SH-SY5Y (Human neuroblastoma, ATCC, CRL-2266) cells and HEK293 (Human embryonic kidney) cells were purchased from ATCC. HEK293FT cells were a generous gift from the MRC PPU. SH-SY5Y, HEK293 and HEK293FT cells were maintained in DMEM GlutaMAX (High glucose; Gibco) supplemented with 10% fetal bovine serum (FBS; Thermo Fisher), 100 U/mL Penicillin-Streptomycin (P/S, Thermo Fisher). ULK1/2 DKO MEF (Mouse embryonic fibroblast cells, a kind gift from Prof. Craig Thompson, Memorial Sloan-Kettering Cancer Center) cells were maintained in DMEM media (Thermo Scientific) supplemented with 10% (v/v) FBS, 2 mM L-glutamine, 100 U/mL penicillin and 0.1 mg/mL streptomycin. For autophagy assays, cells were treated with amino acid–free Earle’s Balanced Salt Solution (EBSS, Fischer) for the indicated duration. All cell lines were grown in a humidified incubator at 37°C, 5% CO_2_ and routinely tested for mycoplasma contamination. All cell culture procedures were performed under sterile conditions in line with biological safety requirements.

#### CRISPR/Cas9 mediated HiBiT Knock-In cell line generation

ULK1-HiBiT KI HEK293 cells were generated via the RNP transfection method of delivering single-stranded oligonucleotides (IDT) as the Alt-R-HDR donor template (IDT, 237042633), spCas9 (Sigma-Aldrich) and target specific Alt-R CRISPR-Cas9 crRNAs (IDT, 237042636, 237042637) hybridised with Alt-R CRISPR-Cas9 tracrRNA (IDT, #1073189) as described here. The sequences or crRNAs and HDR donor template are provided in Table **S1.**

Briefly, two parts of the guides (IDT, trRNA and crRNA) were mixed according to manufacturer’s instructions and annealed at 95°C for 5 min to form gRNA, after gRNAs were cooled down Cas9 was added to the mixture to form an RNP complex. HEK293 cells were resuspended in R buffer and RNP complex was added into the suspension alongside donor template. Cells were electroporated using a Neon Electroporation system (Thermo Fisher). After electroporation, cells were plated in 6-well plates in pre-warmed culture media. 48-72 h post electroporation, knock-in efficiency of the pool cells was tested by HiBiT lytic detection (Promega) using a PheraStar plate reader (BMG). After successful detection of HiBiT signal, the HiBiT KI pool population underwent single cell sorting (Sony Biotechnology, SH800). After 2-3 weeks of recovery in conditioned media and expansion, validation of HiBiT insertion of single cell clones was conducted via HiBiT lytic assay (Promega), western blotting and finally amplicon sequencing (MiSeq, Illumina).

#### Amplicon Sequencing

Genomic DNAs from cell pellets were extracted using QuickExtract DNA extraction solution (Lucigen; Catalog no: #QE09050) according to manufacturer’s’ instructions. Amplicon libraries were generated using a two-step PCR protocol. In the first PCR, target-specific primers with Illumina overhang adapter sequences (Forward: TCGTCGGCAGCGTCAGATGTGTATAAGAGACAGACCAAGTGTGAGTGCCCG, Reverse: GTCTCGTGGGCTCGGAGATGTGTATAAGAGACAGCAGCACACAGAGCCCACTC) were used to amplify the region of interest. PCR reactions were performed in 50 µL volumes using Platinum™ SuperFi II PCR Master Mix (Thermo Fisher Scientific; Catalog no: #12368010) with 2× buffer, 0.5 µM each primer, and 5 µL of template DNA (genomic DNAs from QuickExtraction). Thermal cycling conditions were as follows: initial denaturation at 98°C for 30 s; followed by 30 cycles of 98°C for 10 s, 60°C for 10 s, 72°C for 15 s; and a final extension at 72°C for 5 min. Amplicons were purified using AMPure XP beads (Beckman Coulter) at a 1:1 ratio and eluted in 30 µL of nuclease-free water. For the indexing PCR, dual-index barcodes were added using Illumina indexing primers (Illumina DNA/RNA UD Indexes Set A, Tagmentation (20091654). The Indexing PCR was performed with 8 cycles of amplification using KAPA HiFi HotStart ReadyMix (Roche), followed by another round of AMPure XP bead purification (1:1) and elution in 20 µL 10 mM TRIS pH 8.5. Purified libraries were quantified using the Qubit dsDNA HS Assay Kit (Thermo Fisher Scientific) and library size was verified by capillary electrophoresis on a Qiagen QIAxcel. Equimolar pooling of libraries was performed to generate a final library pool at 4 nM. The pooled library was denatured and diluted according to Illumina protocols and loaded at 8 pM with 10% PhiX control (Illumina) for sequencing. Sequencing was performed on an Illumina MiSeq platform using a MiSeq Nano v2 reagent kit (2 × 150 bp paired-end reads). Demultiplexing and generation of FASTQ files were carried out using Illumina’s MiSeq Reporter software. FASTQ files further analysed on CRISPResso2 software for successful insertion^1^.

#### Stable Cell line generation

Briefly, Flag-ULK1 expressing stable SH-SY5Y cell line was generated as follows: HEK293FT cells were used to produce retroviruses and co-transfected with 3.8 µg pCMV-gag-pol (Cell Biolabs; cat no: #RV-111), 2.2 µg pCMV-VSV-G (Cell Biolabs; Cat no: #RV-110) and 6 µg of pBabeD-Flag-ULK1 WT (MRC Reagents and Services; DU83134). Plasmid was added into 600 µl Opti-MEM (Gibco) and 24 µl of PEI (1 mg/ml, Polysciences, #24765) dissolved in 25 mM HEPES buffer (pH 7.5) and vortexed for 15 s and transfection mixture incubated for 20 min at room temperature and added dropwise onto cells that were plated in 10 cm plates 16 h prior to transfection. Cell media was refreshed 24 h post transfection and viruses were harvested 48 h and 72 h post transfection; harvests were combined and filtered through 0.45 µM syringe filters. The supernatant was then used to transduce target cells in a 1:2 dilution with full growth media supplemented with 8 µg/ml Polybrene (Sigma-Aldrich #TR-1003-G). Cell media was changed for fresh media containing 2 µg/mL Puromycin 24 h post transduction and Puromycin was then removed from culture media after 48 h.

ULK1/2 double-knockout (DKO) stably expressing FLAG-tagged ULK1 and the *Auto-QC* reporter MEF cell lines were generated by retroviral infections. HEK293FT cells co-transfected directly in the growth media with polyethyleneimine/PEI (3:1 ratio) using pBABE-puro-mCherry-GFP-LC3 (*Auto*-QC, DU55696) and pBabeD-hygro-FLAG-ULK1 (DU45617) constructs generated by the MRCPPU Reagents, University of Dundee and are available on the website (https://mrcppureagents.dundee.ac.uk/reagents) with pCMV-gag-pol (Cell Biolabs; cat no: #RV-111) and pCMV-VSV-G (Cell Biolabs; Cat no: #RV-110) to produce retroviral particles. Virus was harvested 48 h post transfection and applied to cells in the presence of 10 µg/ml polybrene. Cells were selected with 500 µg/ml hygromycin (Source Bioscience) or 2 µg/ml puromycin (Sigma) 48 h post transduction. Stable cell pools were used for experiments.

#### Western Blotting

SH-SY5Y and HEK293 were plated in either 6 cm plates or 10 cm plates at varying densities (0.5-1 ×10^6^ cells per plate) depending on experimental set up in respective culturing conditions as described above. Stock solutions of compounds freshly prepared as 10 mM in DMSO (Sigma). Cells were washed twice with ice-cold PBS (Gibco) and pelleted. Cells were resuspended in RIPA buffer, supplemented with cOmplete EDTA-free protease inhibitor cocktail (Roche,11873580001, 1/10mL) and phosSTOP (Roche, 04906845001, 1 tablet/10mL), lysed for 20 min on ice then cleared via centrifugation at 16000×g, 4 °C for 20 min. Following clearance of lysates, protein concentrations were determined by the BCA assay (Fisher Scientific, 23225). 10-30 µg of proteins from lysates were prepared in 4×LDS buffer (Thermofisher) and 50 mM dithiothreitol (DTT), then denatured at 95 °C for 5 min. For western blotting of the RIPA insoluble fraction, the pellet remaining after clearance of lysate was washed with RIPA x3, then resuspended in 0.1 µL of 7 M urea and 1% SDS in RIPA, supplemented as above, for every 1 µg of proteins in the soluble fraction. The resulting suspension was heated on a thermoshaker to 37 °C at 1000 rpm for 2 h then cleared via centrifugation at 16000×g, 4 °C for 20 min. Samples were prepared in 4×LDS buffer (Thermofisher) and 50 mM dithiothreitol (DTT), then denatured at 70 °C for 5 min. Proteins were separated on NuPAGE 4-12% Bis-Tris gels (Invitrogen) in MOPS buffer (Invitrogen) by SDS-PAGE and transferred onto nitrocellulose membranes using an iBlot3 system (Thermo Fisher Scientific) or SureLock™ Tandem Midi Blot Module (Invitrogen). Membranes were blocked with 5% fat-free milk (Marvel) in TBS-T (20 mM Tris (pH 7.5), 150 mM NaCl, 0.1% (v/v) Tween-20 (Sigma) at room temperature (rt) for 30 minutes. MEF cell lines were lysed with RIPA lysis buffer (Tris-HCl pH 8 50 mM, NaCl 150 mM, EDTA 5 mM, Triton 1%, SDS 0.1%, Deoxycholate-Na 0.5%) supplemented with cOmplete, EDTA-free-Protease Inhibitor Cocktail (Roche) and phosphatase inhibitor cocktail (115 mM sodium molybdate, 400 mM sodium tartrate dihydrate, 1 M β-glycerophosphoric acid disodium salt, pentahydrate, 100 mM sodium fluoride, 100 mM activated sodium orthovanadate). Equal amounts of protein (around 10–15 μg) were resolved by 6–14% Bis Tris gels, transferred to nitrocellulose membrane (Amersham Protan 0.45 μm nitrocellulose membrane), blocked in 2% milk in TBST. Membranes were probed with primary antibodies overnight at 4 °C on a rocker. The following primary antibodies were used as indicated dilutions in 5% (w/v) BSA (Sigma) in TBS-T: anti-ULK1 (D8H5) (Cell Signalling Technologies, #8054S, 1;1000), anti-Aurk B (Abcam, #ab2254, 1:1000), anti-IP6K2 (Abcam, #ab252414, 1:1000), anti-LIMK2 (Proteintech, #12350-1-AP, 1:1000), anti-ATG14 (Cell Signalling Technologies, # 96752, 1:250), anti-phospho Ser29 ATG14 (Cell Signalling Technologies, #92340S, 1:1000), anti-ATG13 (Cell Signalling Technologies, #13468T, 1:1000), anti-GSK3 (Cell Signalling Technologies, 5676, lot 7, 1:1000), anti-FLAG HRP (Sigma, A8592, 1:2,000).

Membranes were washed 3 times for 5 minutes with 1x TBS-T on a shaker and incubated with fluorescently labelled or HRP-linked secondary antibodies for 1 hr at RT: anti-mouse IRDye 680RD (LiCOR, 926-68070, 1:5,000), anti-rabbit IRDye 800CW (LiCOR, 926-32211, 1:5,000), anti-mouse StarBright blue 520 (Bio-Rad, 12005866, Bio-Rad, 1:5000), anti-rabbit (LiCOR, 926-32211, 1:10000), anti-mouse (Invitrogen, A21057, 1:10000), goat anti-Rabbit IgG (H+L) HRP conjugate (Thermo Scientific, #31460, 1:5000) and goat anti-mouse IgG (H+L) HRP conjugate (Thermo Scientific, #31430, 1:5,000).. For loading control, hFAB™ Rhodamine Anti-GAPDH Primary Antibody (Bio-Rad, 12004167, 1:10000), hFAB Rhodamine Anti-Actin Primary Antibody (Bio-Rad, 12004164, 1:10000) or anti-Vinculin (Abcam, #ab129002, 1:10000) were used. For chemiluminescence signal detection was performed using ECL (Biorad, Clarity Western ECL Substrate). Images were captured by using Chemidoc imager (Bio-Rad, fluorescence and chemiluminescence) and band intensities were quantified using Image Lab software (Bio-Rad, v6.1).

#### Multiplexing HiBiT Degradation and Cell Viability Assays

ULK1-HiBiT KI HEK293 cells were seeded onto white-opaque 96-well plates at a density of 2×10^4^ cells per well. After 16 h, cells were treated with test compounds or DMSO with indicated concentrations for 4 or 8 h. For mode-of-action studies, cells were pre-treated for 2 hr with following inhibitors: MG132 (1µM, Merck, #474790-5mg), BafA1 (100nM, Enzo Life Sciences, #BML-CM11), TAK-243 (1µM, Selleckchem, #S8341) or Z-VAD-FMK (MedChemExpress, #HY-16658B) and then co-treated with test compounds. 30 minutes before the end of treatment, 20 µl of reaction buffer containing Gly-Phe-AFC substrate (1:200 in cell culture media, Chem Cruz, #sc506265) added to the each well and cells were returned to cell culture incubator. At the end of the treatment time, HiBiT lytic reagents were prepared according to manufacturers’ instructions (Promega) and 60 µL of reagent dispensed to each well. Fluorescence (viability) and luminescence (degradation) signals were measured by GloMAX discover plate reader (Promega) or PheraStar plate reader (BMG, LABTECH Inc.). Luminescence signal was normalised to fluorescence signals and DMSO conditions using Excel and further analysed with GraphPad Prism (v10.1.2). Degradation curves were generated using 4-parameter non-linear regression fit to calculate DC_50_ and D_max_ values.

#### NanoBRET Target Engagement Assays

HEK293 cells were transiently transfected with NanoLuc-ULK1 construct (Promega, NV2211), 48 h post transfection cells resuspended in Opti-MEM (Gibco) were seeded into 96-well white-opaque nonbinding surface plates (Corning) at a density of 2×10^4^ cells per well. Immediately after the addition of test compounds, tracer K-10 (Promega, N2642) was added to a 700 nM final concentration based on our tracer optimisation tests (**Fig. S3A**). Plates were incubated in darkness at 37 °C for 2 h. NanoBRET Nano-Glo Substrate and extracellular NanoLuc Inhibitor (Promega, N1571) were prepared and added per manufacturer instructions. Acceptor and donor signals were measured using a PheraStar plate reader (BMG, LABTECH Inc.). BRET ratios were calculated and normalised to DMSO conditions using Excel and further analysed using GraphPad Prism (v10.1.2). Inhibition curves were generated using a 4-parameter non-linear regression fit to calculate IC_50_ values.

#### Quantitative Unbiased Proteomics

For unbiased profiling of compounds at the proteome level, HEK293 or SH-SY5Y cells were seeded onto 10 cm plates at a density of 2-2.5×10^6^ cells per plate. When cells were confluent, culturing media was refreshed and treated with a freshly prepared 1µM or 3µM final concentration of test compounds (KinoGlu 1, KinoGlu 2, ALP-1**, S2a**, **S2b**, **S2c**, **S2d** and CHX), negative controls (**3c** and **1d**) and DMSO for 4 or 8 hr in 3 or 4 biological replicates. After treatment, cells were washed twice with PBS (Gibco), collected in low binding tubes and lysed in RIPA buffer supplemented with Protease (Roche) and Phosphatase (Roche) inhibitors as previously described for 20 min on ice. Lysates were cleared, and protein concentrations were determined by using Pierce BCA assay (Thermo Scientific). 150 µg of proteins for each condition were aliquoted into LoBind tubes (Eppendorf) and S-Trap (Protifi) protocol has been followed according to manufacturers’ instructions with slight modifications. Briefly, protein lysate was mixed with 2× Lysis buffer (10% SDS, 100mM TEAB pH 8.5), proteins were reduced with 5 mM DTT at 60 °C for 15 min, alkylated (final concentration 20 mM IAA or CAA) at rt for 10 min in dark and acidified (27.5% Phosphoric acid). The solution was diluted with 7 times using binding/wash buffer (100 mM TEAB in 90% methanol). Proteins were then trapped into columns and cleaned with binding/wash buffer (100 mM TEAB in 90% methanol) via centrifugation at 4000×g for 30 sec. Cleaned proteins were then went through tryptic digestion (Trypsin, Thermo Scientific in 50 mM TEAB) overnight at 37°C. Digested peptides were sequentially eluted in 50 mM TEAB (Sigma-Aldrich), 0.2% formic acid (Sigma-Aldrich) and 50% acetonitrile (Sigma-Aldrich). Eluates were pooled and dried down. For TMT proteomics experiments, dried and cleaned peptides were further reconstituted in 110 µl of 100 mM TEAB. TMT-18-plex reagent (Thermo Scientific) was brought to rt and reconstituted in 40 µl of anhydrous acetonitrile. TMT reagent solution was transferred to the digested peptides and incubated on thermoshaker for at 400 rpm at rt for 1hr in dark. Peptides were quenched by the addition of 8 µl of 5% Hydroxylamine for 1hr at rt. TMT-labelled peptides pooled together and dried down in by speed-vac.

#### Immunoprecipitation-Mass Spectrometry

Flag-ULK1 expressing stable SH-SY5Y cells were seeded onto 15-cm plates at a density of 4-5×10^6^ cells per plates. When cells became confluent, they were treated with freshly prepared test compound (ALP-1) and negative controls (**3c** and DMSO) at 3 µM final concentrations for 4 h in 5 biological replicates. After treatment, cells were harvested in ice-cold PBS into LoBind tubes (Eppendorf), proteins were extracted with RIPA buffer and protein concentrations were measured by BCA assay, as previously described. 800 µg of total cell lysates for each treatment condition loaded onto 40µl of Flag-bead slurry (MRC Reagents and Services) in LoBind tubes (Eppendorf) and incubated on tube rotator overnight at 4 °C. Beads were washed three times with extraction buffer and co-immunoprecipitants were eluted with 50 µl of 10% SDS by heating for 5 min at 95°C. Elution mixes were then further processed using the STRAP (Protifi) protocol as previously described for tryptic digestion (Thermo Scientific, 37°C, overnight) of co-immunoprecipitated proteins. Digested peptides were sequentially eluted in 50 mM TEAB (Sigma-Aldrich), 0.2% formic acid (Sigma-Aldrich) and 50% acetonitrile (Sigma-Aldrich). Elutions were pooled and dried down. Dried and cleaned peptides were further reconstituted in 110 µl of 100 mM TEAB for TMT labelling. TMT-18-plex reagent (Thermo Scientific) was brought to rt and reconstituted in 40 µl of anhydrous acetonitrile. TMT reagent solution was transferred to the digested peptides and incubated on thermo-shaker for at 400 rpm at rt for 1hr in dark. Peptides were quenched by the addition of 8 µl of 5% Hydroxylamine for 1 h at rt. TMT-labelled peptides pooled together and dried down by speed-vac.

Pooled peptides were separated by basic reverse phase chromatography fractionation on a C_18_, column with flow rate at 200 μL/min with two buffers: buffer A (10 mM ammonium formate, pH 10) and buffer B (80% ACN, 10 mM ammonium formate, pH 10). Peptides were resuspended in 100 μL of buffer A (10 mM ammonium formate, pH10) and resolved on a C_18_ reverse phase column by applying a non-linear gradient of 7–40%. A total of 80 fractions were collected and concentrated into 30 fractions.

The dried peptides were reconstituted in 0.1% formic acid and analysed on an Orbitrap Ascend mass spectrometer coupled to a Thermo Fisher Scientific Vanquish Neo UHPLC. The peptides were separated on an analytical column (Acclaim PepMap RSLC C18, 75 μm × 50 cm, 2 μm, 100 Å) at a flow rate of 300 nL/min, using a step gradient of 2–7% solvent B (90% ACN/0.1% FA) for the first 6 min, followed by 7–18% up to 89 min, 18–27% up to 89–114 min and 27-35% to 114-134 min. The total run time was set to 155 min. The mass spectrometer was operated in a data-dependent acquisition mode in SPS MS3 (FT-IT-HCD-FT-HCD) method. A survey full scan MS (from m/z 350–1500) was acquired in the Orbitrap at a resolution of 120,000 at 200 m/z. The AGC target for MS1 was set as 4 × 105 and ion filling time as 50 ms. The precursor ions for MS2 were isolated using a Quadrupole mass filter at a 0.7 Da isolation width, fragmented using a normalized 30% HCD of ion routing multipole and analysed using ion trap. The top 10 MS2 fragment ions in a subsequent scan were isolated and fragmented using HCD at a 55% normalized collision energy and analysed using an Orbitrap mass analyser at a 60,000 resolution, in the scan range of 100–500 m/z.

The proteomics raw data were searched using SEQUEST HT search engines with Proteome Discoverer 3.0 (Thermo Fisher Scientific). The following parameters were used for searches: Precursor mass tolerance 10 ppm, Fragment mass tolerance 0.1, Enzyme: trypsin, Mis-cleavage: −2, Fixed modification: carbamidomethylation of cysteine residues and TMT of lysine and N-terminal, Dynamic modification: oxidation of methionine. The data were filtered for 1% PSM, peptide and protein level FDR. Only unique peptides were selected for the quantification.

#### ULK1 kinase domain digestion and mass spectrometry analysis

In-solution digestion of purified ULK1 C182S mutant was performed by dissolving 5 µg of ALP-1-labelled protein in 50 mM triethylammonium bicarbonate (TEAB). The protein was digested with 500 ng of Asp-N or trypsin for 12–14 h at 37 °C. Following digestion, the samples were acidified with trifluoroacetic acid (TFA) to a final concentration of 1% and centrifuged at high speed to remove any insoluble or undigested material. The peptides were desalted using C_18_ material on zip tip and dried using speedvac. The dried peptides were reconstituted in 1% formic acid and analysed on an Orbitrap Ascend Tribrid mass spectrometer coupled with Thermo Fisher Scientific Vanquish Neo UHPLC. The peptides were enriched on a trap column and separated on an analytical column (Easy-Spray PepMap Neo C18, 2 μm, 75 μm × 150 mm) at 300 nl/min. Chromatographic separation was performed using a gradient elution: 5% BufferA (0,1% formic acid) and 35% buffer B (90% ACN, 0,1% Formic Acid) at 70 min. The total run time was 90 min. The mass spectrometer was operated in a data-dependent acquisition mode. A survey full scan MS (from *m*/*z* 300 to 1600) was acquired in the Orbitrap at a resolution of 120,000 at 200*m*/*z*. The AGC target for MS1 was set as 4 x10^5^ and the ion filling time as 251 ms. The most intense ions with charge state ≥2 were isolated and fragmented using higher collision dissociation (HCD) fragmentation, with 27% normalized collision energy, and detected at a mass resolution of 30,000 at 200*m*/*z*. For MS2 acquisition, precursor ions were isolated using an isolation window of 1.2 m/z. Fragment ions were detected in the ion trap, with an AGC target of 1 × 10⁴ and a maximum injection time of 35 ms. The proteomics raw data were searched using SEQUEST HT search engines with Proteome Discoverer 3.0 (Thermo Fisher Scientific) and database of E. coli with addition of ULK1 sequence. The following parameters were used for searches: Precursor mass tolerance 10 ppm, Fragment mass tolerance 0.1, Enzyme: AspN for Asp-N and trypsin for trypsin digestion, Mis-cleavage: −1, Dynamic modification: oxidation of methionine and ALP modification on Aspartate. The data were filtered for 1% PSM, peptide.

#### Microscopy

Cells stably expressing *Auto*-QC reporter system (mCherry-GFP-LC3) were seeded onto sterile glass coverslips in 24-well dishes. After treatment, coverslips were washed twice with PBS, fixed with 3.7% (w/v) formaldehyde, 200 mM HEPES pH 7.0 for 10 min and washed twice with PBS. After nuclei counterstaining with 1 μg/mL Hoechst-33258 dye, slides were washed and mounted in ProLong Gold (Invitrogen).

#### Flow cytometry analysis

A total of 2.5 × 10^5^ stably expressing *Auto*-QC reporter cells were seeded in a 6 cm dish. Following treatment, cells were washed with PBS, trypsinised for 5 min, neutralized with PBS containing 2% FBS, and centrifuged at 1,200 rpm for 4 min. The pellet of cells was resuspended in 100 μL of PBS and 0.5 mL of 3.7% (w/v) formaldehyde, 200 mM HEPES pH 7.0 were added. After 30 min at rt, 2 ml of PBS + 2% FBS was added before centrifugation 5 min at 1200 rpm. Finally, the pellet of cells was resuspended in 2% FCS in PBS and analysed by flow cytometry. For each independent experiment, at least 50000 events were acquired on LSRFortessa cell analyzer. Based on FCS and SSC profiles, living cells were gated. As negative control, cells expressing any autophagy reporter were used. To quantify the percentage of cells undergoing autophagy, the ratio GFP/mCherry was analysed. The gate used for the nontreated condition or control cells was applied to all the other conditions.

Representative results of at least three independent experiments (biological replicates) are shown in all panels. Data are presented as mean and standard deviations (s.d.). GraphPad Prism software was used for all statistical analyses. Statistical significance was determined using two-way ANOVA with Tukey’s multiple comparisons test. *P*-values are indicated as \**P* < 0.05, \*\**P* < 0.01, \*\*\**P* < 0.001 and \*\*\*\**P* < 0.0001. ns: *P* > 0.05.

#### Confocal microscopy

APRE19 cells stably expressing GFP-ULK1 were seeded onto sterile glass coverslips in 12-well dishes. After treatment, coverslips were washed twice with PBS, fixed with 3.7% (w/v) paraformaldehyde, 200 mM HEPES pH 7.0 for 15 min and washed twice with PBS. Then slides were washed and mounted in ProLong Glass antifade (Thermo Fisher, P36980) on a SuperFrost microscope slide (VWR; 631-0847) and cured in the dark overnight before imaging on a Zeiss 880 Airyscan microscope (Alpha Plan-APO 100×/1.46 oil DIC VIS objective (Zeiss) and Alpha Plan-APO 63×/1.4 oil objective (Zeiss)).

#### Purification of recombinant ULK1 kinase domain

The DNA construct encoding kinase domain of ULK1 (residues 1-283) was synthesised by Genescript. The codon optimised sequence encoding ULK1 kinase domain was cloned into pET28a vector with N-terminal 6x-His tag followed by TEV protease cleavage site, T7 tag, AviTag, and a cleavage site for PreScission protease. The C182S mutation was introduced by site-directed mutagenesis. Briefly, the plasmid encoding wild type ULK1 was amplified by PCR using Q5® High-Fidelity 2X Master Mix (New England Biolabs, #M0492S) and a set of primers with forward primer introducing C182S mutation (Forward: CAATATGATGGCTGCCACCCTT**TCC**GGTAG; Reverse: GACTGCAGGTAACGCGCAAAACCGAAG).

The PCR product was treated with DpnI enzyme (New England Biolabs, #R0176S) to remove template DNA and was purified using NucleoSpin™ Gel and PCR Clean-up kit (Marcherey Nagel, #740609). The purified DNA was circularised by treating with T4 polynucleotide kinase (Thermo Scientific, #EK0031) and T4 DNA ligase (Thermo Scientific, #EL0014). The resulting construct was transformed in *E. coli* DH5-alpha and isolated using Monarch Plasmid Miniprep Kit (New England Biolabs, #T1110L). The plasmid sequence was confirmed by whole plasmid sequencing. To obtain constructs with double mutations, the procedure was repeated with pET28a-His-TEV-AviTag-3C-ULK1 KD C182S construct as template and primers introducing point mutations C47A (Forward: AG**GCG**ATTAACAAGAAAAACCTGGCAAAAAGCC; Reverse: TAACCGCAACTTCCAGGTCGTG) or D102A (Forward: TGGCT**GCG**TATCTCCACGCTATGCGCACC;

Reverse: AATCACCACCATTGCAGTACTCCATC). The open reading frames of the plasmids are available in Table **S2.** The ORF sequence of ULK1 KD C47A/C182S was recloned in pRSF-DUET-MBP plasmid using Gibson assembly method. Briefly, the pRSF-DUET-MBP plasmid backbone was digested with BamHI and XhoI restriction enzymes (New England Biolabs #R0136S and #R0146S) and purified from agarose gel using NucleoSpin™ Gel and PCR Clean-up kit (Marcherey Nagel, #740609). The ULK1 insert was amplified by PCR using primers with overhands complementary to the plasmid backbone (Forward: AG**GCG**ATTAACAAGAAAAACCTGGCAAAAAGCC; Reverse: TAACCGCAACTTCCAGGTCGTG) or D102A (Forward: **GGAAAACCTGTATTTTCAGG**GATCCGGACTAAATGATATATTTG; Reverse: CTTTACCAGACTCGAGTGCTTAA**GACGGGCTTGCGTCCAG**). The PCR product was treated with DpnI enzyme and purified as described above. Then, 50 pmol of digested plasmid and 50 pmol of the insert were added to 5 µL of NEBuilder® HiFi DNA Assembly Master Mix (new England Biolabs #E2621L) and incubated at 40 °C for 10 min, then at 60 °C for 60 min. The mixture was then transformed in *E. coli* DH5-alpha. The plasmid was then isolated and sequenced as described above.

ULK1-encoding constructs were co-transformed with a plasmid encoding lambda protein phosphatase in *E. coli* Lemo21 (DE3) strain (New England Biolabs, #C2528J) and grown at 30 °C with selection antibiotics (50 µg/mL kanamycin, 100 µg/mL ampicillin, 30 µg/mL chloramphenicol) in presence of 50 µM rhamnose to repress protein expression in absence of IPTG. The C47A/C182S and D102A/C182S mutants were expressed without lambda phosphatase, and they were cultured in the same conditions, but without the addition of ampicillin in the growth media. Overnight cultures of transformed bacteria were diluted 50 times in TB media and grown at 30 °C until the cultures reached OD600 = 1.6-1.8. The cultures were then cooled down to 18 °C for 1 h before the protein expression was induced with 0.4 mM IPTG and grown overnight.

The bacteria were then collected by centrifugation and resuspended in Ni buffer A (25 mM Tris pH 7.5, 300 mM NaCl, 10 mM imidazole, 0.5 mM TCEP, 10% glycerol) with addition of cOmplete EDTA-free Protease Inhibitor Cocktail (Roche, #11873580001) and benzonase. The bacteria were lysed with Cell Disruptor homogeniser (Constant Systems), the clarified lysate was loaded on 5 mL HisTrap HP column (Cytiva) and the proteins were eluted in a gradient of Ni buffer B (25 mM Tris pH 7.5, 300 mM NaCl, 500 mM imidazole, 0.5 mM TCEP, 10% glycerol). The protein-containing fractions were pooled and loaded in SnakeSkin dialysis tubing 10,000 MWCO (Thermo Scientific, #88243) with PreScission protease and dialysed overnight at 4 °C against dialysis buffer (20 mM Tris pH 7.0, 100 mM NaCl, 1 mM TCEP, 10% glycerol). The following day, the protein was diluted to NaCl concentration 45 mM with dilution buffer (20 mM Tris pH 7.0, 0.5 mM TCEP, 10% glycerol) and loaded on 5 mL HiTrap SP HP column (Cytiva). The column was washed with 3 column volumes of S buffer A (20 mM Tris pH 7.0, 45 mM NaCl, 0.5 mM TCEP, 10% glycerol) and eluted in 0-30% gradient of S buffer B (20 mM Tris pH 7.0, 1 M NaCl, 0.5 mM TCEP, 10% glycerol). The protein eluted in multiple peaks corresponding to different phosphorylation forms of ULK1. The fractions corresponding to monophosphorylated (wild type, C182S, and D102A/C182S mutants) or double-phosphorylated protein (C47A/C182S mutant) were pooled, concentrated in 10,000 MWCO centrifugal filter (Amicon) and further purified on Superdex75 10/300 GL column (Cytiva) in SEC buffer (25 mM Tris pH 7.5, 150 mM NaCl, 10% glycerol). The protein was concentrated, aliquoted, flash frozen in liquid nitrogen, and stored at -80 °C.

#### Analysis of intact ULK1 modification by LC-MS

ULK1 kinase domain proteins were incubated at 10 µM with 10 µM of ALP-1, **3a**, or with 2% DMSO. The incubation was carried out in a buffer containing 25 mM Tris pH 7.5 and 150 mM NaCl for 18 h at room temperature. The samples were then transferred to 4 °C, separated on a ZORBAX 300SB-C3 column in 10-95% gradient of acetonitrile with 0.05% TFA for 12 minutes, and analysed using Agilent 6135C single quadrupole InfinityLab Pro iQ Series Mass Detect. Fragmentor voltage was 220 V, and the capillary voltage was 4000 V. The mass spectrometer acquired full scan spectra at m/z 600-2500 in positive scan mode. Protein spectra were deconvoluted using Agilent LC/MSD OpenLab software.

#### Kinetic measurement of ULK1 covalent modification by LC-MS

ULK1 WT kinase domain was incubated at 10 µM with ALP-1 at concentrations of 2, 4, 6 or 8 µM or with 2% DMSO in a buffer containing 25 mM Tris pH 7.5 and 150 mM NaCl. The protein and ALP-1 mixture was incubated at room temperature, and 5 µL aliquots were injected on ZORBAX 300SB-C3 after 1, 2, 4, 8, 12, 16, and 20 h incubation. The control DMSO-treated sample was measured after 0 h incubation. The samples were measured as described above. The ratio of labelled protein was calculated by dividing relative abundance of labelled protein by a sum of relative abundance of both labelled and unlabelled protein species. The values were plotted using GraphPad Prism and kinact/Ki parameter was calculated by fitting the data into a bimolecular reaction model as previously described^2^.

#### Fluorescence polarisation assay

Fluorescence polarisation assay was performed using a fluorescein-labelled tracer based on DCC-3116 ULK1 inhibitor. To obtain binding curve of the tracer, a serial 2-fold dilutions of ULK1 KD WT ranging from 500 to 0.06 nM were titrated against 10 nM of the tracer at 5% DMSO concentration. The assay buffer contained 25 mM Tris pH 7.5, 150 mM NaCl, 0.1% Tween-20 and 1 mg/mL BSA. The measurement was performed in a black 384-well plate. After preparing the reactions the plate was sealed, centrifuged for 1 min at 100 xg and incubated for 15 min at room temperature. The plate was then measured using PheraStar plate reader (BMG, LABTECH Inc.) with excitation at λ = 485 nm and emission at λ = 520 nm. Gain adjustment was performed using wells with free tracer and set for 35 mP. The binding curves were obtained by plotting polarisation against logarithm of ULK1 concentration and fitting four parameters variable slope curve using GraphPad Prism. ULK1 concentration used for displacement assays was 15 nM and corresponded to IC_80_ of the tracer. For displacement assay, a mix of 15 nM ULK1 and 10 nM tracer was mixed with 3-fold dilutions of ALP-1, **3a** or **3c** at concentrations ranging from 30 µM to 0.2 nM or with 5% DMSO. The plate was then sealed, centrifuged and measured as described above after 15, 30, 60, 120, 240, 360, 480, 600, and 1440 min incubation. The normalised displacement was calculated by subtracting background polarisation of free tracer and dividing the value by polarisation of DMSO-treated samples. The values were then multiplied by a factor of 100 to obtain a percentage of bound tracer and then transformed to percentage values of displaced tracer. To obtain IC_50_ values, the normalised displacement was plotted against logarithm of compound concentration and fitted into four parameters variable slope curve. To visualise displacement kinetics, the relative displacement values for each compound concentration were plotted against time.

#### Differential scanning fluorimetry

2 µM of ULK1 WT KD was incubated with ALP-1, **3a** or **3c** at 10 µM or with 4% DMSO in a buffer containing 25 mM Tris pH 7.5 and 150 mM NaCl. The protein was incubated with the compounds for 0.5, 2, 4, 8, 12, and 24 h before the measurement. For endpoint measurements ULK1 WT or mutants (C47A/C182S and D102A/C182S) were incubated in identical conditions with incubation time of 18 h. After incubation, 24 µL aliquots of the reactions were transferred to a 96-well PCR plate in triplicates. Then 1 µL of SYPRO Orange stain (Invitrogen #S6650) was added to the samples to the final concentration of x5. The plate was sealed, centrifuged for 1 min at 100 xg, and transferred to the BioRad C1000 thermal cycler with CFX96 Touch Real-Time PCR Detection System module. The plate was incubated at 20 °C for 3:30 min and then incubated in temperature increasing from 20 to 95 °C with the increase rate of 1 °C/min. The fluorescence was read after each temperature increase. The raw data was inspected by plotting relative fluorescence (RFU) against temperature. The curves showing abnormal profile with steep fluorescence increase or decrease at the beginning of the temperature gradient were excluded from the analysis. For each measured sample at least two technical replicates were analysed, and the experiment was repeated to obtain 3 biological repeats. The data was further analysed by calculating the first derivative of relative sample fluorescence (dRFU/dT) and plotting against temperature. To calculate melting temperature (Tm) the datapoints were fitted into Gaussian model using GraphPad Prism. The curves showing two distinct peaks were fitted using sum of two Gaussians model.

#### Crystallisation of ULK1 kinase domain in complex with ALP-1

The ULK1 KD C182S co-crystals were obtained by incubating purified ULK1 C182S at 2 mg/mL with equimolar amount of ALP-1 at final DMSO concentration at 2% in protein storage buffer containing 25 mM Tris pH 7.5, 150 mM NaCl, and 10% glycerol. After overnight incubation at RT, the labelling was confirmed by intact mass spectrometry. The labelled protein was concentrated to 18 mg/mL and used for crystallisation screening in sitting drop with 200 nL of protein and 200 nL reservoir solution. The crystallisation plates were incubated at 20 °C in a crystal hotel. We observed formation of microcrystals in the conditions containing 200 mM NaCl, 100 mM Bis-Tris pH 6.5, and 25% PEG 3350. The microcrystals were resupended in reservoir buffer diluted 1:1 with 100 mM Bis-Tris pH 7.5 and milled into fined fragments. The resulting seed stock was further diluted 100 times in the reservoir buffer and used in the second round of screening. For screening with seeding, the ULK1 C182S was labelled as described above and concentrated to 15 mg/mL. The screening plates were set up in sitting drop using 200 nL of protein, 150 nL of reservoir buffer, and 50 nL of diluted seed stock. Protein crystallised in conditions containing 200 mM NaCl, 100 mM Tris pH 8.5 amd 25% PEG 3350.

Protein crystal was harvested and flash-frozen in liquid nitrogen. X-ray diffraction data were collected at Diamond Light Source at beamline i24 using wavelength 0.6199 Å. The processing was performed with autoPROC^3^ and data reduction was performed using AIMLESS^4^. Further processing and structure determination was performed using CCP4 software suite^5^. Molecular replacement was performed using Phaser^6^ with reference model PDB ID 4WNO using only peptide chain atoms from the reference structure. Structure refinement was performed using REFMAC5^7^ and Phenix Refine^8^ and manual model building was performed in Coot^9^. The ALP-1 structure was truncated to avoid overfitting and only model the part of the ligand with well-defined electron density. After several cycles of manual building and refinement the final structure model was visualised in ChimeraX^10^.

## Synthesis Experimental

All chemicals unless otherwise stated, were commercially available, at least 95% pure and used without further purification. All reactions were carried out under nitrogen atmosphere with anhydrous solvents, unless aqueous reagents were used or otherwise stated. Commercially available dry solvents were used. Water was purified by an Milli-Q IQ 7010 system. All other solvents were used as supplied (analytical or HPLC grade) without prior purification. Microwave reactions were carried out using a Biotage Initiator+ Microwave System. Organic layers were dried over Na_2_SO_4_. Brine refers to a saturated aqueous solution of NaCl. Concentration of solvents *in vacuo* was achieved by rotary evaporation using a diaphragm pump. Normal phase TLC was carried out on pre-coated silica plates (Kieselgel 60 F254, BDH) with visualisation *via* UV light (254 nm) or basic potassium permanganate solution. Flash column chromatography was performed using a Teledyne Isco Combiflash NextGen 300+ with prepacked Redisep normal phase disposable columns. Reverse phase chromatography was carried out on a Teledyne Isco Combiflash NextGen 300+ using either RediSep C18 reversed phase columns or Biotage SNAP-C18 Columns. Preparative HPLC was performed on a Waters Preparative HPLC System with a Waters XBridge C18 column (100 mm x 19 mm; 5 µm particle size) and a gradient of 5–95% MeCN in water over 10 minutes, flow 25 mL/min, with 0.1% formic acid in the aqueous phase.

NMR Spectra were recorded on a Bruker Avance NEO 500 MHz spectrometer in the deuterated solvent stated. Spectra were recorded at 298 K. The field was locked by external referencing to the relevant deuteron resonance. Chemical shifts are reported in ppm and coupling constants (*J*) in Hz. High Resolution Mass Spectra (HRMS) were recorded on a Bruker MicroTOF II focus ESI mass spectrometer connected in parallel to a Dionex Ultimate 3000 RSLC system with a diode array detector and a Waters X-Bridge C18 column (50 mm x 2.1 mm, 3.5 µm particle size). Mass Spectra (MS) and analytical HPLC traces were recorded on a Shimadzu LCMS-2020 or -2050 equipped with a Hypersil Gold column (1.9 µm 50 × 2.1 mm^2^), a photodiode array detector, and an electrospray ionization (ESI) detector. The samples were eluted with a 3 or 5 min gradient of 5−95% MeCN in water containing either 0.1% formic acid or 0.1% ammonia at a flow rate of 0.8 mL/min.

***tert*-Butyl 4-(4-((5-chloro-4-((2-(isopropylsulfonyl)phenyl)amino)pyrimidin-2-yl)amino)phenyl)piperazine-1-carboxylate (1)**

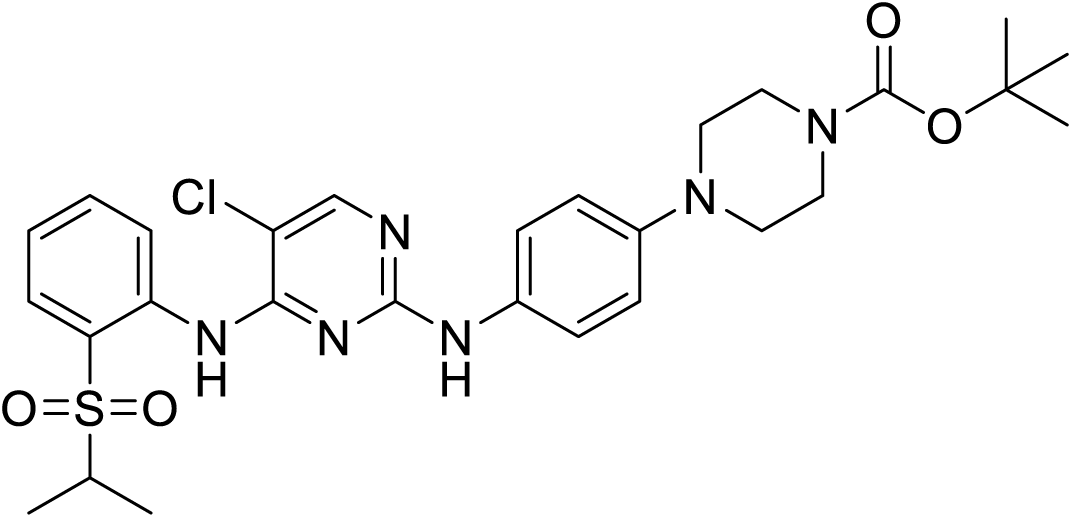

TFA (660 µL, 8.7 mmol) was added to a solution of 2,5-dichloro-*N*-(2-isopropylsulfonylphenyl)pyrimidin-4-amine (2.50 g, 7.22 mmol) and *tert*-butyl 4-(4-aminophenyl)piperazine-1-carboxylate (2.40 g, 8.66 mmol) in *s*-BuOH (25 mL). The reaction mixture was heated to 80 °C for 16 h, then concentrated *in vacuo*. The crude residue was purified by flash column chromatography (CH_2_Cl_2_/MeOH, 0→10%) to afford the title compound (2.76 g, 4.51 mmol, 63%) as a yellow solid. **^1^H NMR** (500 MHz, CDCl_3_) δ 9.85 (s, 1H), 8.55 (dd, J = 8.6, 1.1 Hz 1H), 8.04 (s, 1H), 7.90 (dd, J = 8.0, 1.6 Hz, 1H), 7.54 (ddd, J = 8.7, 7.3, 1.7 Hz, 1H), 7.44 – 7.37 (m, 2H), 7.31 – 7.26 (m, 1H), 6.94 – 6.86 (m, 2H), 3.63 – 3.55 (t, J=5.1 Hz, 4H), 3.22 (hept, J = 6.9 Hz, 1H), 3.10 (t, J = 5.1 Hz, 4H), 1.49 (s, 9H), 1.31 (d, J = 6.9 Hz, 6H); **^13^C NMR** (126 MHz, CDCl_3_) δ 156.5, 155.9, 154.7, 148.0, 137.8, 134.5, 131.4, 124.8, 123.8, 123.6, 122.5, 117.2, 105.6, 80.0, 55.9, 50.0, 28.5, 15.4; **MS** *m/z* (ESI^+^) 587 [M+H]^+^.

**5-Chloro-*N***^4^**-(2-(isopropylsulfonyl)phenyl)-*N***^2^**-(4-(piperazin-1-yl)phenyl)pyrimidine-2,4-diamine (2)**

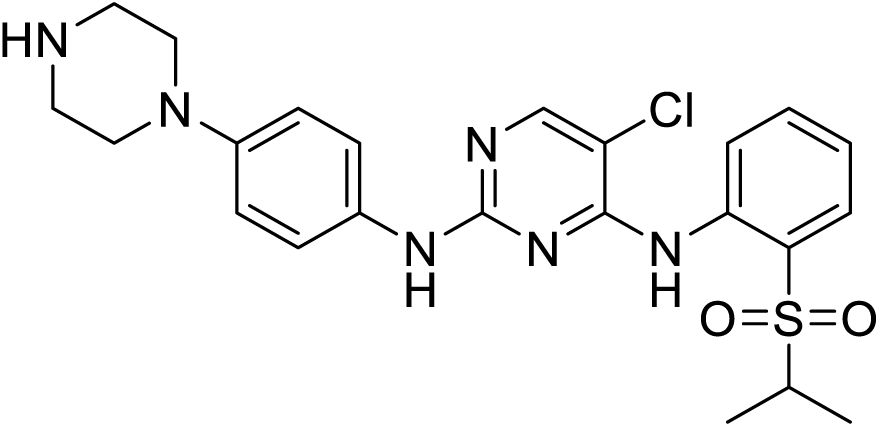

TFA (1.9 mL, 7.66 mmol) was added to a solution of **1** (300 mg, 0.511 mmol) in CH_2_Cl_2_ (3 mL). Reaction mixture stirred at rt for 16 h, then concentrated *in vacuo*. The residue was freebased by SCX column to afford the title compound as a white solid (307 mg, 0.511 mmol, quant.). **^1^H NMR** (500 MHz, CDCl_3_) δ 9.59 (s, 1H), 8.59 (dd, *J* = 8.5, 1.1 Hz, 1H), 8.06 (s, 1H), 7.85 (dd, *J* = 8.0, 1.7 Hz, 1H), 7.77 (s, 1H), 7.50 (ddd, *J* = 8.7, 7.2, 1.7 Hz, 1H), 7.39 – 7.32 (m, 2H), 7.21 – 7.14 (m, 1H), 6.91 – 6.83 (m, 2H), 3.21 (hept, *J* = 6.9 Hz, 1H), 3.09 (dd, *J* = 6.4, 3.4 Hz, 4H), 3.02 (dd, *J* = 6.4, 3.4 Hz, 4H), 2.03 – 1.83 (m, 1H), 1.27 (d, *J* = 6.9 Hz, 6H); **^13^C NMR** (126 MHz, CDCl_3_) δ 158.3, 155.3, 155.1, 148.2, 138.5, 134.5, 131.7, 131.1, 124.1, 123.4, 122.9, 122.3, 116.7, 105.4, 55.6, 51.0, 46.1, 15.3; **MS** *m/z* (ESI^+^) 488 [M+H]^+^.

**2-chloro-1-(4-(4-((5-chloro-4-((2-(isopropylsulfonyl)phenyl)amino)pyrimidin-2-yl)amino)phenyl)piperazin-1-yl)ethan-1-one (1a, KinoGlu 1)**

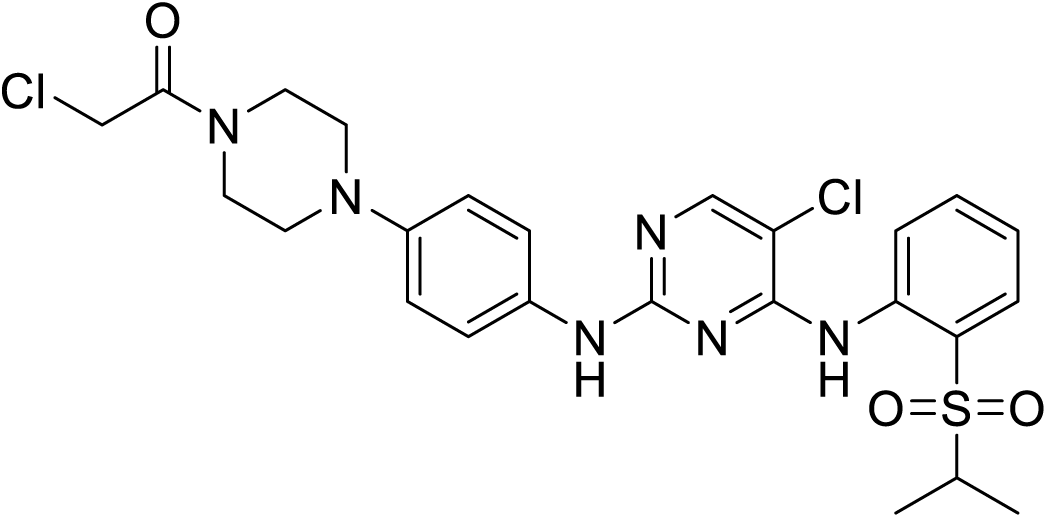

Chloroacetyl chloride (19 µL, 0.21 mmol) was added to a suspension of **2** (103 mg, 0.211 mmol) and DIPEA (110 µL, 0.65 mmol) in DMA (2 mL) Reaction mixture was concentrated *in vacuo* and purified by preparative-HPLC to afford the title compound (80 mg, 0.142 mmol, 67%) as a white solid. **^1^H NMR** (500 MHz, CDCl_3_) δ 9.62 (s, 1H), 8.59 (dd, *J* = 8.6, 1.1 Hz, 1H), 8.11 (s, 1H), 7.89 (dd, *J* = 8.1, 1.7 Hz, 1H), 7.56 (ddd, *J* = 8.6, 7.3, 1.7 Hz, 1H), 7.46 – 7.37 (m, 2H), 7.24 (ddd, *J* = 8.1, 7.3, 1.1 Hz, 1H), 7.18 (br s, 1H), 6.94 – 6.87 (m, 2H), 4.12 (s, 2H), 3.80 (d, *J* = 5.3 Hz, 2H), 3.70 (t, *J* = 5.0 Hz, 2H), 3.28 – 3.17 (m, 3H), 3.14 (t, *J* = 5.3 Hz, 2H), 1.31 (d, *J* = 6.9 Hz, 6H); **^13^C NMR** (126 MHz, CDCl_3_) δ 165.3, 158.1, 155.5, 155.3, 147.1, 138.6, 134.5, 132.8, 131.4, 124.6, 123.5, 123.2, 122.0, 117.7, 106.1, 55.7, 50.5, 50.1, 46.4, 42.3, 40.9, 15.5; **MS** *m/z* (ESI^+^) 563 [M+H]^+^; **HRMS** calcd for C_25_H_28_Cl_2_N_6_O_3_S+H^+^ 563.13934 found 563.14047.

**1-(4-(4-((5-Chloro-4-((2-(isopropylsulfonyl)phenyl)amino)pyrimidin-2-yl)amino)phenyl)piperazin-1-yl)prop-2-en-1-one (1b, KinoGlu 2)**

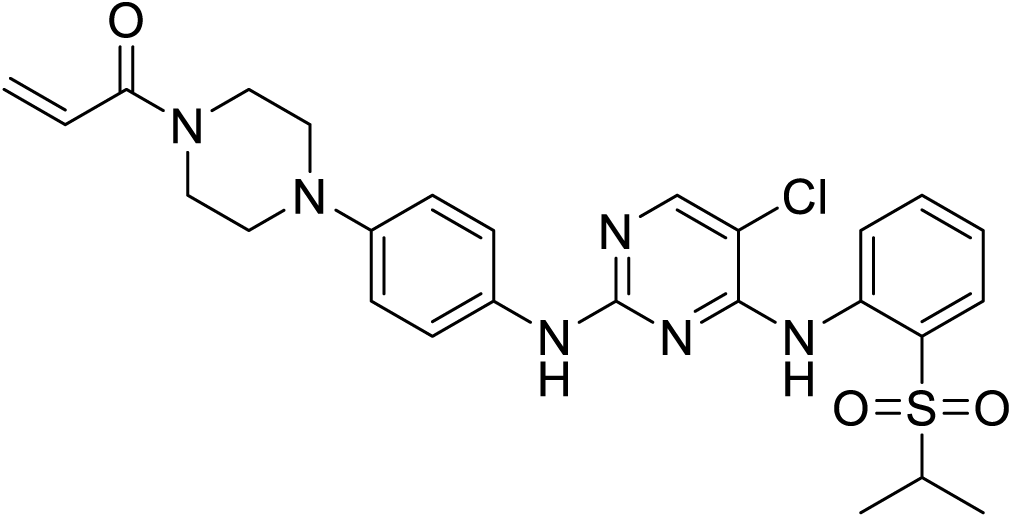

Acryloyl chloride (21 µL, 0.28 mmol), followed by DIPEA (150 µL, 0.86 mmol) were added to a solution of **2** hydrochloride (90 mg, 0.172 mmol) in DMA (2 mL). The reaction mixture was stirred at rt for 2 h, then purified by preparative-HPLC to afford the title compound (44 mg, 0.0821 mmol, 48%) as a white solid. **^1^H NMR** (500 MHz, CDCl_3_) δ 9.62 (s, 1H), 8.59 (d, *J* = 8.4 Hz, 1H), 8.10 (d, *J* = 2.7 Hz, 1H), 7.89 (ddd, *J* = 8.0, 2.7, 1.7 Hz, 1H), 7.58 – 7.52 (m, 1H), 7.44 – 7.39 (m, 2H), 7.25 – 7.18 (m, 1H), 6.90 (dd, *J* = 9.0, 2.2 Hz, 2H), 6.61 (dd, *J*= 16.9, 10.6 Hz, 1H), 6.33 (dd, *J* = 16.9, 1.9 Hz, 1H), 5.73 (dd, *J* = 10.6, 1.9 Hz, 1H), 3.86 (bs, 2H), 3.74 (bs, 2H), 3.28 – 3.17 (m, 1H), 3.14 (t, *J* = 6.9 Hz, 4H), 1.30 (dd, *J* = 6.9, 2.2 Hz, 6H); **^13^C NMR** (126 MHz, CDCl_3_) δ 165.5, 158.2, 155.5, 155.3, 147.3, 138.6, 134.5, 131.4, 128.3, 127.4, 124.5, 123.5, 123.2, 122.1, 117.6, 106.0, 55.7, 50.7, 50.1, 45.9, 42.0, 15.5; **MS** *m/z* (ESI^+^) 541 [M+H]^+^; **HRMS** calcd for C_26_H_29_O_3_N_6_ClS+H^+^ 541.17831 found 541.17940.

**5-Chloro-*N*^4^-(2-(isopropylsulfonyl)phenyl)-*N*^2^-methyl-*N*^2^-(4-(piperazin-1-yl)phenyl)pyrimidine-2,4-diamine (3)**

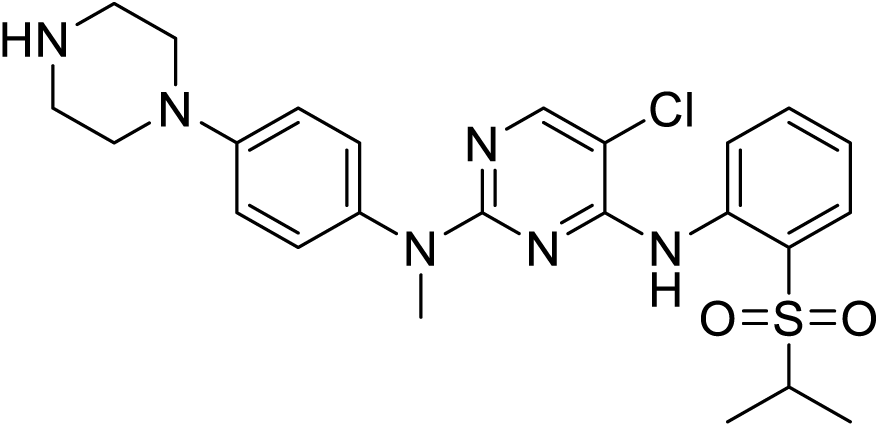

TFA (70 µL, 0.92 mmol) was added to a solution of 2,5-dichloro-*N*-(2-isopropylsulfonylphenyl)pyrimidin-4-amine (265 mg 0.765 mmol) and *tert*-butyl 4-(4-(methylamino)phenyl)piperazine-1-carboxylate (268 mg, 0.918 mmol) in *sec*-BuOH (2.6 mL). The reaction mixture was heated to 80 °C for 16 h, then TFA (700 µL, 9.2 mmol) was added and the reaction mixture heated for a further 16 h. The reaction mixture was concentrated *in vacuo*, then purified by flash column chromatography (CH_2_Cl_2_/MeOH, 0→15%) to afford the title compound as a colourless oil. **^1^H NMR** (500 MHz, CDCl_3_) δ 9.84 (s, 1H), 9.80 (br s, 2H), 8.40 (d, *J* = 8.5 Hz, 1H), 8.07 (s, 1H), 7.84 (dd, *J* = 7.8, 1.7 Hz, 1H), 7.34 (t, *J* = 8.5 Hz, 1H), 7.22 (d, *J* = 8.5 Hz, 2H), 7.17 (t, *J* = 7.8 Hz, 1H), 6.99 (d, *J* = 8.5 Hz, 2H), 3.50 – 3.44 (m, 8H), 3.39 (br s, 4H), 3.21 (hept, *J* = 6.9 Hz, 1H), 1.29 (d, *J* = 6.9 Hz, 6H); **^13^C NMR** (126 MHz, CDCl3) δ 155.2, 148.7, 138.4, 134.5, 131.4, 128.1, 124.0, 123.2, 118.0, 105.3, 55.9, 47.0, 43.6, 39.3, 15.4; **MS** *m/z* (ESI^-^) 499 [M-H]^-^.

**2-Chloro-1-(4-(4-((5-chloro-4-((2-(isopropylsulfonyl)phenyl)amino)pyrimidin-2-yl)(methyl)amino)phenyl)piperazin-1-yl)ethan-1-one (1c)**

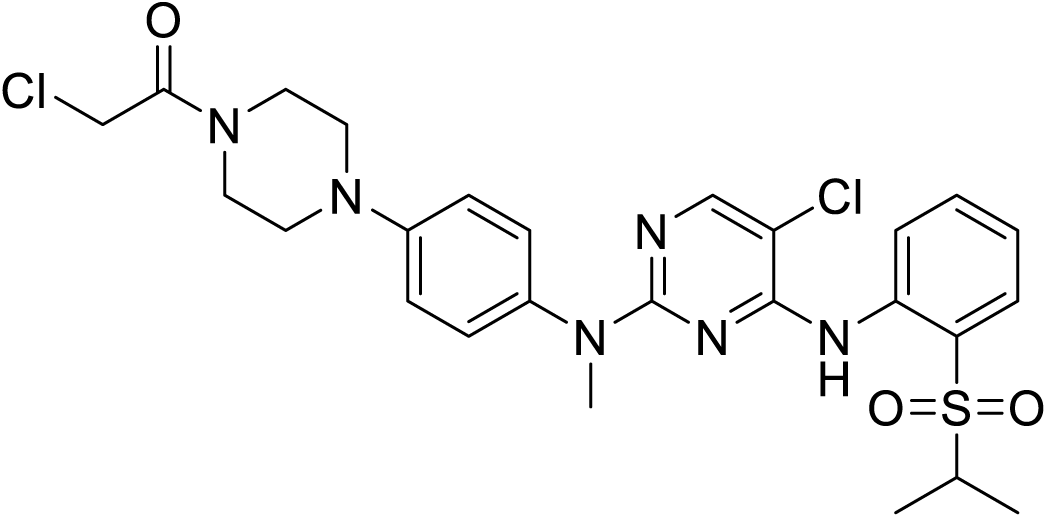

Chloroacetyl chloride (17 µL, 0.22 mmol) was added to a suspension of **3** (100 mg, 0.200 mmol) and DIPEA (100 µL, 0.60 mmol) in DMA (2 mL). Reaction mixture was concentrated *in vacuo* and purified by preparative-HPLC to afford the title compound (25 mg, 0.0433 mmol, 22%) as a white solid. **^1^H NMR** (500 MHz, CDCl_3_) δ 9.66 (s, 1H), 8.44 (d, *J* = 8.7 Hz, 1H), 8.08 (s, 1H), 7.82 (dd, *J* = 7.9, 1.7 Hz, 1H), 7.29 (m, 1H), 7.21 (dt, *J* = 8.8, 2.1 Hz, 2H), 7.14 – 7.06 (m, 1H), 7.01 – 6.94 (m, 2H), 4.13 (s, 2H), 3.82 (t, *J* = 5.2 Hz, 2H), 3.72 (t, *J* = 5.2 Hz, 2H), 3.46 (s, 3H), 3.33 – 3.16 (m, 5H), 1.29 (d, *J* = 6.9 Hz, 6H);. **^13^C NMR** (126 MHz, CDCl_3_) δ 165.3, 160.1, 155.0, 154.8, 149.0, 139.1, 138.7, 134.4, 131.2, 128.0, 123.4, 122.3, 122.3, 117.5, 117.4, 105.0, 55.7, 49.9, 49.6, 46.4, 42.2, 40.9, 39.0, 15.5; **MS** *m/z* (ESI^+^) 577 [M+H]^+^; **HRMS** (ESI^+^) *m/z* calcd for C_26_H_30_O_3_N_6_Cl_2_S+H^+^ 577.15499 found 577.15614.

**1-(4-(4-((5-Chloro-4-((2-(isopropylsulfonyl)phenyl)amino)pyrimidin-2-yl)amino)phenyl)piperazin-1-yl)propan-1-one (1d)**

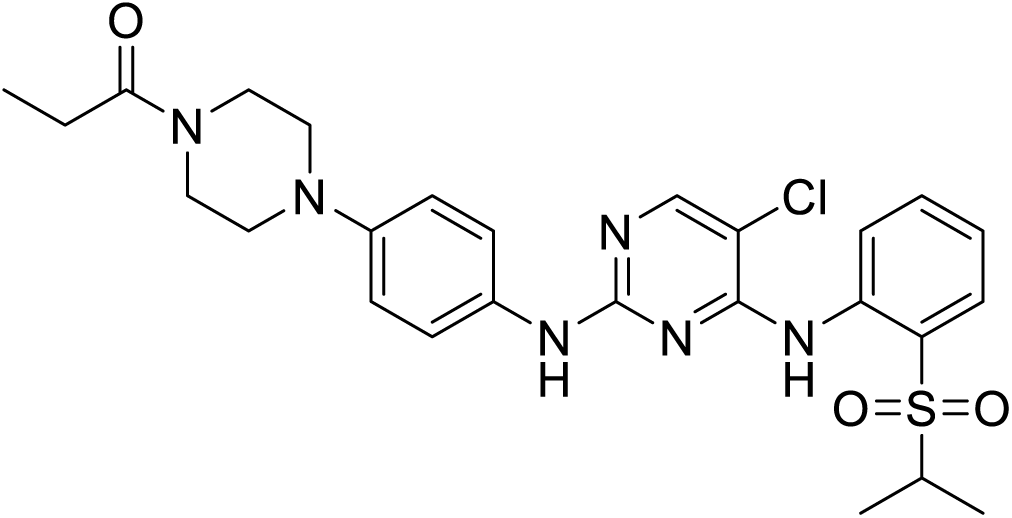

A solution of **2** (82 mg, 0.136 mmol), propionic acid (10 µL, 0.14 mmol), HATU (73 mg, 0.191 mmol) and DIPEA (120 µL, 0.68 mmol) in DMF (1 mL) was stirred at rt for 16 h. Reaction mixture was partitioned between water and EtOAc, then extracted with EtOAc x2. Combined organic layers were washed with brine x3, dried over Na_2_SO_4_ and concentrated *in vacuo*. The crude product was purified by flash column chromatography (heptane/EtOAc, 50→100%) followed by preparative-HPLC to afford the title compound (50 mg, 0.0921 mmol, 67%) as a white solid. **^1^H NMR** (500 MHz, CDCl3) δ 9.62 (s, 1H), 8.59 (d, *J* = 8.2 Hz, 1H), 8.11 (s, 1H), 7.89 (d, *J* = 7.8 Hz, 1H), 7.55 (t, *J* = 8.2 Hz, 1H), 7.41 (d, *J* = 8.6 Hz, 2H), 7.23 (t, *J* = 7.8 Hz, 1H), 7.14 (br s, 1H), 6.90 (d, *J* = 8.6 Hz, 2H), 3.80 (t, *J* = 5.2 Hz, 2H), 3.63 (t, *J* = 5.2 Hz, 2H), 3.23 (hept, *J* = 6.9 Hz, 1H), 3.12 (dt, *J* = 11.2, 5.2 Hz, 4H), 2.40 (q, *J* = 7.4 Hz, 2H), 1.30 (d, *J* = 6.9 Hz, 6H), 1.18 (t, *J* = 7.4 Hz, 3H); **^13^C NMR** (126 MHz, CDCl_3_) δ 172.4, 158.2, 155.5, 155.3, 147.4, 138.6, 134.5, 132.5, 131.4, 124.5, 123.5, 123.2, 122.1, 117.5, 106.0, 55.7, 50.6, 50.2, 45.5, 41.6, 26.6, 15.5, 9.6; **MS** *m/z* (ESI^+^) 543 [M+H]^+^; **HRMS** (ESI^+^) *m/z* calcd for C_26_H_31_O_3_N_6_ClS+H^+^ 543.19396 found 543.19493.

***tert*-Butyl 6-((4-chloro-5-(trifluoromethyl)pyrimidin-2-yl)amino)-3,4-dihydroisoquinoline-2(1*H*)-carboxylate (4)**

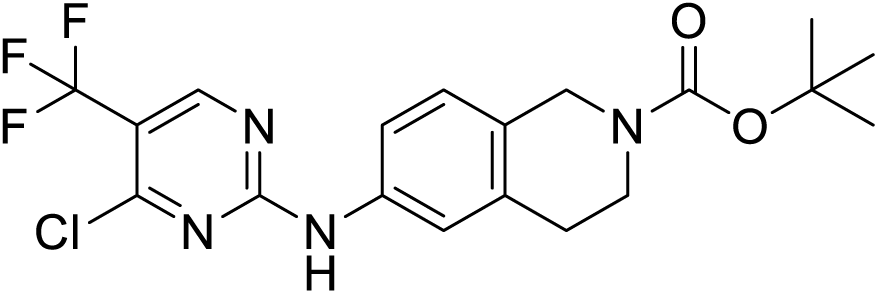

A solution of 2,4-dichloro-5-(trifluoromethyl)pyrimidine (600 mg, 2.77 mmol) and ZnCl_2_ (452 mg, 3.32 mmol) in DCE/*t*-BuOH (1:1) (25 mL) was stirred at 0 °C for 1 h. Then a solution of 6-amino-2-*N*-Boc-1,2,3,4-tetrahydro-isoquinoline (687 mg, 2.77 mmol) and NEt_3_ (460 µL, 3.3 mmol) in DCE/*t*-BuOH (1:1) (5 mL) was added. The solution was stirred at 0 °C for a further 3 h, then concentrated *in vacuo* to afford crude product, which was purified by flash column chromatography (heptane/EtOAc, 0→50%) to afford the title compound (1030 mg, 2.40 mmol, 87%) as a white solid. **^1^H NMR** (500 MHz, DMSO-*d_6_*) δ 10.61 (s, 1H), 8.79 (s, 1H), 7.54 – 7.46 (m, 2H), 7.15 (d, *J* = 8.3 Hz, 1H), 4.46 (s, 2H), 3.54 (t, *J* = 5.9 Hz, 2H), 2.75 (t, *J* = 5.9 Hz, 2H), 1.43 (s, 9H); **^13^C NMR** (126 MHz, DMSO-*d_6_*) δ 160.5, 158.2, 154.0, 136.5, 134.9, 126.6, 121.9, 120.3, 120.2, 118.7, 118.6, 79.0, 28.5, 28.1; **^19^F NMR** (471 MHz, DMSO-*d_6_*) δ -60.07; **MS** *m/z* (ESI^-^) 427 [M-H]^-^.

***tert*-Butyl 6-((4-(cyclopropylamino)-5-(trifluoromethyl)pyrimidin-2-yl)amino)-3,4-dihydroisoquinoline-2(1*H*)-carboxylate (5)**

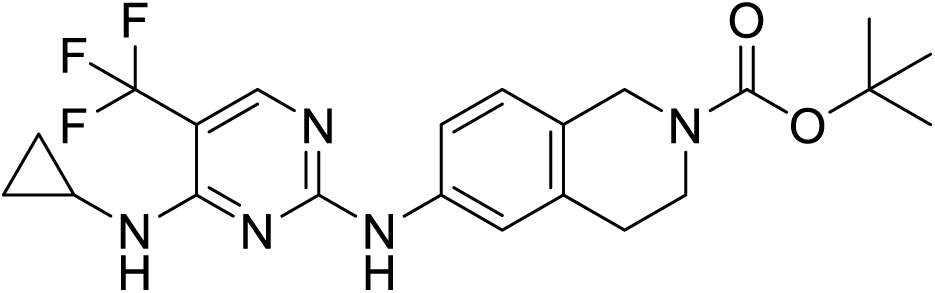

A solution of **4** (1.03 g, 2.40 mmol), cyclopropylamine (400 µL, 5.8 mmol) and DIPEA (500 µL, 5.8 mmol) in DMF (10 mL) was heated to 100 °C under microwave irradiation for 15 min. The reaction mixture was concentrated *in vacuo* to afford crude product, which was then purified by flash column chromatography (heptane/EtOAc, 0→50%) to afford the title compound (956 mg, 2.13 mmol, 89%). **^1^H NMR** (500 MHz, CDCl3) δ 8.16 (d, *J* = 0.9 Hz, 1H), 7.61 (br s, 1H), 7.53 – 7.40 (m, 2H), 7.05 (d, *J* = 8.3 Hz, 1H), 5.37 (br s, 1H), 4.54 (s, 2H), 3.65 (s, 2H), 2.88 (ttd, *J* = 7.0, 3.8, 2.3 Hz, 1H), 2.82 (t, *J* = 5.9 Hz, 2H), 1.49 (s, 9H), 0.94 – 0.86 (m, 2H), 0.69 – 0.61 (m, 2H); **^13^C NMR** (126 MHz, CDCl_3_) rotameric mixture δ 161.0, 160.3, 155.1, 154.8 (q, *J* = 5.0 Hz), 137.6, 135.4, 128.2, 126.7, 126.1, 124.0, 121.8, 119.6, 118.0, 99.1 (q, *J* = 32.4 Hz), 79.9, 45.8, 45.1, 41.9, 40.7, 29.4, 28.6, 24.4, 7.6; **^19^F NMR** (471 MHz, CDCl_3_) δ -61.27 (s, 1F); **MS** *m/z* (ESI^+^) 450 [M+H]^+^.

***N*^4^-Cyclopropyl-*N*^2^-(1,2,3,4-tetrahydroisoquinolin-6-yl)-5-(trifluoromethyl)pyrimidine-2,4-diamine (6)**

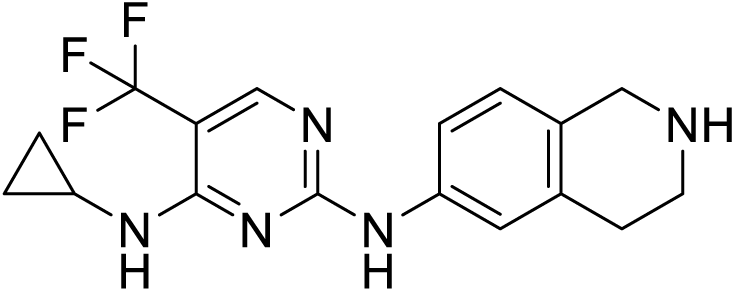

TFA (670 µL, 8.7 mmol) was added to a solution of **5** (167 mg, 0.372 mmol) in CH_2_Cl_2_ (1.5 mL). Reaction mixture stirred at rt for 2 h, then concentrated *in vacuo*. Residue passed through an SCX column to afford the title compound (130 mg, 0.372 mmol, quant.). **^1^H NMR** (500 MHz, CDCl_3_) δ 8.14 (s, 1H), 8.10 (br s, 1H), 7.53 (s, 1H), 7.42 (dd, *J* = 8.3, 2.3 Hz, 1H), 6.94 (d, *J* = 8.3 Hz, 1H), 5.40 (s, 1H), 3.97 (s, 2H), 3.12 (t, *J* = 6.0 Hz, 2H), 2.86 (dddd, *J* = 9.6, 9.6, 3.8, 3.2 Hz, 1H), 2.76 (t, *J* = 6.0 Hz, 2H), 2.32 (br s, 2H), 0.93 – 0.83 (m, 2H), 0.66 – 0.59 (m, 2H); **^13^C NMR** (126 MHz, CDCl_3_) δ 161.0, 160.2, 154.6 (q, *J* = 5 Hz), 137.4, 135.1, 130.4, 126.4, 125.0 (q, *J* = 270 Hz), 120.2, 117.7, 98.6 (q, *J* = 32 Hz), 47.9, 43.8, 29.4, 24.2, 7.4; **^19^F NMR** (471 MHz, CDCl_3_) δ -61.20; **MS** *m/z* (ESI^+^) 350 [M+H]^+^.

**2-Chloro-1-(6-((4-(cyclopropylamino)-5-(trifluoromethyl)pyrimidin-2-yl)amino)-3,4-dihydroisoquinolin-2(1*H*)-yl)ethan-1-one (2a, ALP-1)**

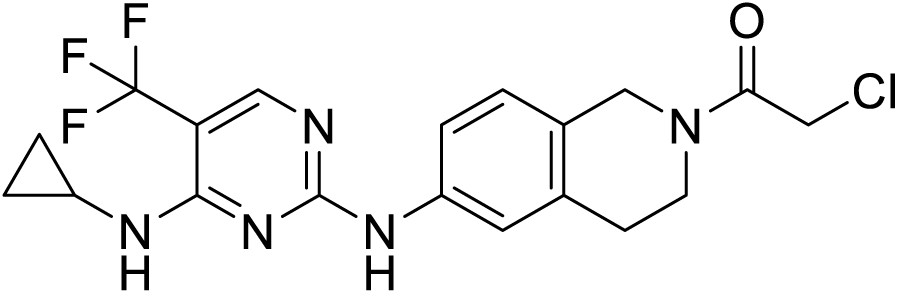

Chloroacetyl chloride (21 µL, 0.27 mmol) was added to a solution of **6** (112 mg, 0.242 mmol) and DIPEA (130 µL, 0.73 mmol) in DMA (2.5 mL). Reaction mixture stirred at rt for 16 h, then concentrated *in vacuo*. The crude product was purified by preparative-HPLC to afford the title compound (64 mg, 0.149 mmol, 62%) as a white solid. **^1^H NMR** (500 MHz, CDCl_3_) 2:3 rotameric mixture δ 8.15 (s, 1H), 7.68 (s, 0.4H), 7.62 (s, 0.6H), 7.55 – 7.47 (m, 1H), 7.42 – 7.34 (m, 1H), 7.08 (dd, *J* = 11.8, 8.3 Hz, 1H), 5.38 (s, 1H), 4.71 (s, 1.2H), 4.66 (s, 0.8H), 4.16 (m, 2H), 3.84 (t, *J* = 5.9 Hz, 0.8H), 3.75 (t, *J* = 5.9 Hz, 1.2H), 2.95 (t, *J* = 5.9 Hz, 1.2H), 2.92 – 2.84 (m, 1.8H), 0.96 – 0.87 (m, 2H), 0.70 – 0.62 (m, 2H); **^13^C NMR** (126 MHz, CDCl_3_) δ 165.7, 165.6, 160.9, 160.3, 154.8, 154.7, 138.4, 138.0, 134.3, 127.1, 126.5, 119.6, 119.1, 118.4, 47.7, 44.6, 44.0, 41.5, 41.4, 40.6, 29.8, 28.7, 24.4, 7.6; **^19^F NMR** (471 MHz, CDCl_3_) δ -61.39; **MS** *m/z* (ESI^+^) 426 [M+H]^+^; **HRMS** (ESI^+^) *m/z* calcd for C_19_H_19_ON_5_ClF_3_+H^+^ 426.13030 found 426.13080.

***tert*-Butyl (3-(cyclobutanecarboxamido)propyl)carbamate (7)**

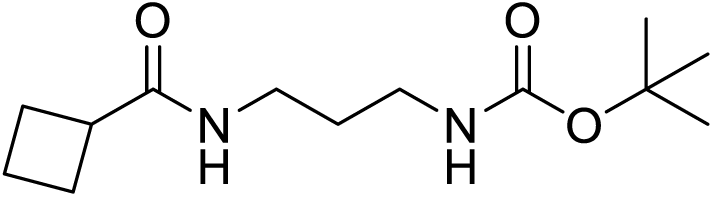

Oxalyl chloride (2.62 mL. 30.6 mmol), followed by DMF (1 mL) were added to a solution of cyclobutanecarboxylic acid (2.55 g, 25.5 mmol) in CH_2_Cl_2_ (160 mL) at 0 °C. The reaction mixture was stirred at 0 °C for 20 min, then at rt for 1 h. The reaction mixture was concentrated *in vacuo* to afford the crude acid chloride intermediate, which was used immediately without further purification. A solution of 563 mg of crude intermediate, 3-(tert-butoxycarbonylamino)propylammonium chloride (1.00 g, 4.75 mmol) and NEt_3_ (1.98 mL) in CH_2_Cl_2_ was stirred at rt for 16 h. The reaction mixture was then washed with brine, concentrated *in vacuo* and purified by flash column chromatography (CH_2_Cl_2_/MeOH, 0→10%) to afford the title compound (469 mg, 1.83 mmol, 39% over two steps) as a colourless oil. **^1^H NMR** (500 MHz, CDCl_3_) δ 6.15 (br s, 1H), 5.04 (br s, 1H), 3.27 (q, *J* = 6.3 Hz, 2H), 3.13 (q, *J* = 6.3 Hz, 2H), 2.99 (p, *J* = 8.6 Hz, 1H), 2.30 – 2.19 (m, 2H), 2.18 – 2.06 (m, 2H), 1.99 – 1.88 (m, 1H), 1.88 – 1.80 (m, 1H), 1.58 (p, *J* = 6.3 Hz, 2H), 1.41 (s, 9H); **^13^C NMR** (126 MHz, CDCl_3_) δ 175.6, 156.7, 79.3, 40.1, 37.1, 35.8, 30.4, 28.5, 25.5, 18.3; **MS** *m/z* (ESI^+^) 257 [M+H]^+^.

***N*-(3-Aminopropyl)cyclobutanecarboxamide hydrochloride (8)**

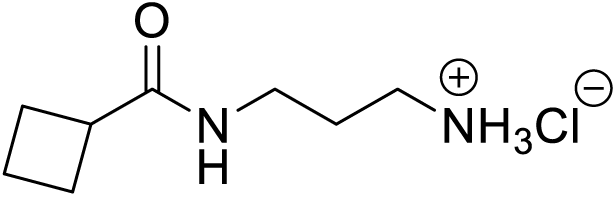

A solution of **7** (469 mg, 1.83 mmol) in HCl/dioxane (4M, 4.60 mL, 183 mmol) was stirred at rt for 2 h. The reaction mixture was concentrated *in vacuo* to afford the title compound (352 mg, 1.83 mmol, quant.) as a white solid. **^1^H NMR** (500 MHz, MeOD) δ 3.28 (q, *J* = 6.6 Hz, 2H), 3.14 (p, *J* = 8.6 Hz, 1H), 2.93 (t, *J* = 6.6 Hz, 2H), 2.30 – 2.19 (m, 2H), 2.19 – 2.11 (m, 2H), 2.05 – 1.94 (m, 1H), 1.90 – 1.81 (m, 3H); **^13^C NMR** (126 MHz, MeOD) δ 178.7, 40.6, 38.2, 36.8, 28.7, 26.3, 19.0; **MS** *m/z* (ESI^+^) 157 [M+H]^+^.

***N*-(3-((5-Bromo-2-chloropyrimidin-4-yl)amino)propyl)cyclobutanecarboxamide (9)**

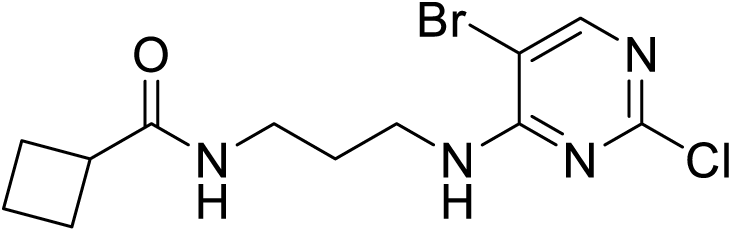

DIPEA (2.53 mL, 14.5 mmol), followed by **8** (352 mg, 1.83 mmol) was added dropwise to a solution of 5-bromo-2,4-dichloro-pyrimidine (416 mg, 1.83 mmol) in EtOH (10 mL) at -10 °C. The reaction mixture was heated to 50 °C for 5 h, then concentrated *in vacuo*. The residue was taken up in EtOAc, washed with water, then brine. Organics were dried over Na_2_SO_4_, then concentrated *in vacuo*. The residue was purified by flash column chromatography (heptane/EtOAc, 0→100%) to afford the title compound (477 mg, 1.37 mmol, 75%). **^1^H NMR** (500 MHz, CDCl_3_) 8.09 (s, 1H), 6.64 (t, *J* = 6.4 Hz, 1H), 5.98 (br s, 1H), 3.54 (q, *J* = 6.4 Hz, 2H), 3.31 (q, *J* = 6.4 Hz, 2H), 3.06 (p, *J* = 8.6 Hz, 1H), 2.38 – 2.23 (m, 3H), 2.22 – 2.10 (m, 2H), 2.05 – 1.91 (m,1H), 1.91 – 1.79 (m, 1H), 1.72 (p, *J* = 6.4 Hz, 2H); **^13^C NMR** (126 MHz, CDCl_3_) δ 176.3, 159.9, 159.3, 156.5, 103.3, 40.1, 37.6, 35.8, 29.8, 25.6, 18.3; **MS** *m/z* (ESI^+^) 349 [M+H]^+^.

***N*-(3-((2-Chloro-5-cyclopropylpyrimidin-4-yl)amino)propyl)cyclobutanecarboxamide (10)**

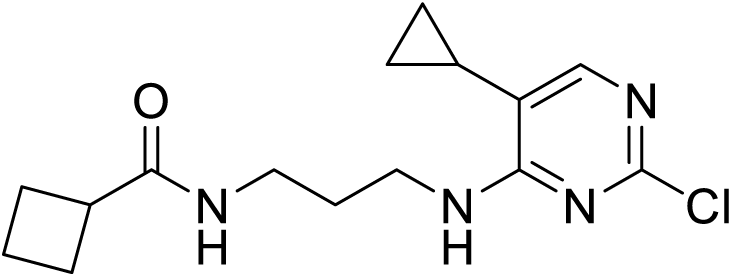

A degassed solution of **9** (477 mg, 1.37 mmol), cyclopropylboronic acid (141 mg, 1.65 mmol), K_2_CO_3_ (569 mg, 4.12 mmol) and Pd(ddpf)Cl_2_ (100 mg, 0.137 mmol) in DME (6mL) and water (4 mL) was stirred at 85 °C for 16 h. Reaction mixture was partitioned between EtOAc and water, the aqueous layer was extracted with EtOAc x2. Combined organics were washed with brine and dried over Na_2_SO_4_ and concentrated *in vacuo*. Crude product purified by flash column chromatography (CH_2_Cl_2_/MeOH; 0→10%) to afford the title compound (259 mg, 0.839 mmol, 61%) as a brown oil. **^1^H NMR** (500 MHz, CDCl_3_) δ 7.73 (d, *J* = 1.2 Hz, 1H), 6.44 (t, *J* = 6.4 Hz, 1H), 6.22 (t, *J* = 6.4 Hz, 1H), 3.54 (q, *J* = 6.4 Hz, 2H), 3.29 (q, *J* = 6.4 Hz, 2H), 3.06 (p, *J* = 8.6 Hz, 1H), 2.33 – 2.21 (m, 2H), 2.19 – 2.11 (m, 2H), 2.02 – 1.79 (m, 2H), 1.74 – 1.66 (m, 2H), 1.48 (ttd, *J* = 8.2, 5.4, 1.2 Hz, 1H), 0.99 (dtd, *J* = 8.2, 5.4, 1.2 Hz, 2H), 0.56 – 0.52 (m, 2H); **^13^C NMR** (126 MHz, CDCl_3_) δ 176.1, 163.7, 158.5, 153.9, 117.6, 40.1, 36.9, 35.6, 30.0, 25.6, 18.3, 7.6, 4.8; **MS** *m/z* (ESI^+^) 309 [M+H]^+^.

***tert*-Butyl 6-((4-((3-(cyclobutanecarboxamido)propyl)amino)-5-cyclopropylpyrimidin-2-yl)amino)-3,4-dihydroisoquinoline-2(1*H*)-carboxylate (11)**

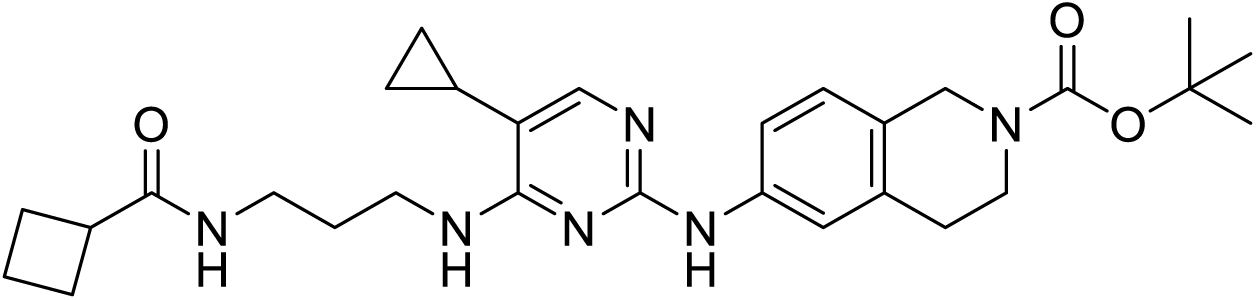

A solution of **10** (207 mg, 0.670 mmol) and 6-Amino-2-*N*-Boc-1,2,3,4-tetrahydro-isoquinoline (250 mg, 1.01 mmol) in AcOH (4 mL) was heated to 110 °C under microwave irradiation for 10 min. Reaction mixture was basified with 2M NaOH (aq.), then partitioned between CH_2_Cl_2_ and water. Aqueous layer extracted with CH_2_Cl_2_ x2. Organics dried over Na_2_SO_4_ and concentrated *in vacuo*. The crude residue was purified by flash column chromatography (heptane/EtOAc; 0→100%) to afford the title compound (216 mg, 0.415 mmol, 62%) as a brown solid. **^1^**H NMR (500 MHz, CDCl3) δ 7.71 (d, *J* = 1.1 Hz, 1H), 7.53 – 7.38 (m, 2H), 7.36 (d, *J* = 8.4 Hz, 1H), 7.00 (d, *J* = 8.4 Hz, 1H), 5.94 – 5.87 (m, 2H), 4.51 (s, 2H), 3.62 (br s, 2H), 3.57 (q, *J* = 6.2 Hz, 2H), 3.29 (q, *J* = 6.3 Hz, 2H), 2.79 (t, *J* = 5.8 Hz, 2H), 2.71 (p, *J* = 8.8 Hz, 1H), 2.20 (pd, *J* = 8.8, 2.4 Hz, 2H), 2.06 – 1.96 (m, 2H), 1.92 – 1.77 (m, 2H), 1.77 – 1.71 (m, 2H), 1.47 (s, 9H), 1.45 – 1.41 (m, 1H), 0.88 (m, 2H), 0.51 – 0.44 (m, 2H); **^13^C NMR** (126 MHz, CDCl_3_) δ 175.6, 162.6, 158.6, 155.0, 153.8, 139.0, 135.2, 126.6, 118.6, 117.2, 110.3, 79.8, 53.5, 40.7, 39.9, 37.1, 35.9, 30.2, 29.4, 28.6, 25.5, 18.2, 7.6, 4.6; **MS** *m/z* (ESI^+^) 521 [M+H]^+^.

***N*-(3-((5-Cyclopropyl-2-((1,2,3,4-tetrahydroisoquinolin-6-yl)amino)pyrimidin-4-yl)amino)propyl)cyclobutanecarboxamide (12)**

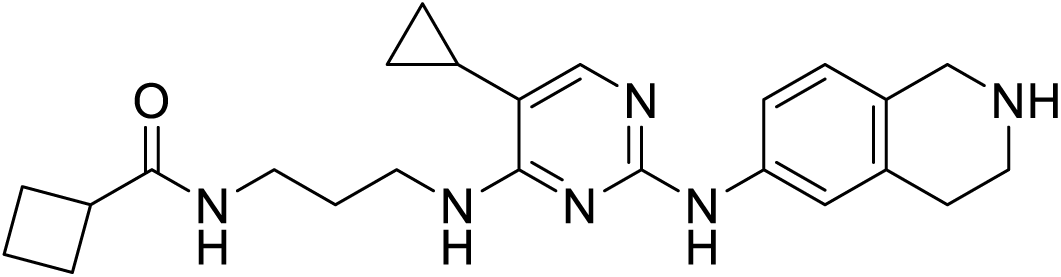

TFA (200 µL, 2.6 mmol) was added to a solution of **11** (101 mg, 0.194 mmol) in CH_2_Cl_2_ (1 mL). Reaction mixture stirred at rt for 3 h, then concentrated *in vacuo*. The crude residue was freebased by SCX column to afford the title compound (85 mg, 0.194 mmol, quant.) as a brown oil. **^1^H NMR** (500 MHz, CDCl3) δ 7.66 (d, *J* = 1.0 Hz, 1H), 7.52 (s, 1H), 7.36 (d, *J* = 2.4 Hz, 1H), 7.29 (dd, *J* = 8.3, 2.3 Hz, 1H), 6.88 (d, *J* = 8.3 Hz, 1H), 6.12 (t, *J* = 6.3 Hz, 1H), 5.91 (t, *J* = 6.3 Hz, 1H), 3.92 (s, 2H), 3.53 (q, *J* = 6.2 Hz, 2H), 3.37 (s, 2H), 3.25 (q, *J*= 6.3 Hz, 2H), 3.08 (t, *J* = 6.0 Hz, 2H), 2.88 (s, 2H), 2.76 – 2.71 (m, 3H), 2.30 – 2.07 (m, 3H), 2.04 – 1.94 (m, 2H), 1.92 – 1.62 (m, 4H), 1.45 – 1.37 (m, 1H), 0.90 – 0.80 (m, 2H), 0.48 – 0.40 (m, 2H); **^13^C NMR** (126 MHz, CDCl_3_) δ 175.7, 162.5, 158.5, 153.4, 138.7, 134.8, 128.7, 126.4, 119.3, 116.9, 110.1, 50.1, 47.7, 43.8, 39.7, 37.1, 35.8, 30.0, 29.3, 25.4, 18.2, 7.5, 4.6 **MS** *m/z* (ESI^+^) 421 [M+H]^+^.

***N*-[3-[[2-[[2-(2-Chloroacetyl)-3,4-dihydro-1H-isoquinolin-6-yl]amino]-5-cyclopropyl-pyrimidin-4-yl]amino]propyl]cyclobutanecarboxamide (2b)**

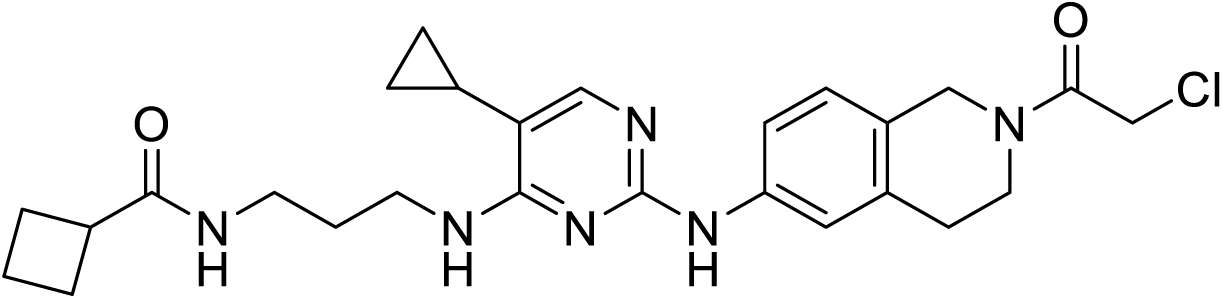

Chloroacetyl chloride (15 µL, 0.19 mmol) followed by DIPEA (89 µL, 0.51 mmol) was added to a solution of **12** (78 mg, 0.171 mmol) in DMA (1.6 mL). Reaction mixture stirred at rt for 16 h, then purified by preparative-HPLC to afford the title compound (12 mg, 0.0241 mmol, 12%) as an off-white solid. **^1^H NMR** (500 MHz, CDCl_3_) half formate salt, rotameric mixture δ 8.67 (s, 0.5H), 7.58 (d, *J* = 2.2 Hz, 0.4H), 7.55 (dd, *J* = 8.3, 2.3 Hz, 0.6H), 7.49 (d, *J* = 7.1 Hz, 1H), 7.47 – 7.41 (m, 1H), 7.12 – 7.00 (m, 2H), 5.90 (q, *J* = 6.8 Hz, 1H), 4.68 (s, 1.2H), 4.63 (s, 0.8H), 4.16 (s, 0.8H), 4.15 (s, 1.2H), 3.81 (t, *J* = 6.0 Hz, 0.8H), 3.72 (t, *J* = 5.9 Hz, 1.2H), 3.57 (q, *J* = 6.1 Hz, 2H), 3.35 (qd, *J* = 6.8, 3.2 Hz, 2H), 2.99 – 2.89 (m, 2.2H), 2.84 (t, *J* = 6.0 Hz, 0.8H), 2.30 – 2.18 (m, 2H), 2.17 – 2.06 (m, 3H), 2.01 – 1.89 (m, 1H), 1.89 – 1.81 (m, 1H), 1.80 – 1.69 (m, 2H), 1.52 (td, *J* = 8.2, 4.2 Hz, 1H), 1.02 – 0.94 (m, 2H), 0.53 – 0.43 (m, 2H); **^13^C NMR** (126 MHz, CDCl3) half formate salt, rotameric mixture δ 176.5, 170.8, 165.7, 165.6, 163.0, 162.9, 160.4, 155.2, 145.8, 138.6, 138.2, 135.1, 134.3, 126.9, 126.8, 126.4, 126.0, 119.6, 119.3, 118.4, 118.2, 111.1, 111.1, 47.6, 44.6, 44.0, 41.6, 41.4, 40.7, 40.0, 37.5, 35.9, 29.8, 29.7, 28.7, 25.6, 18.3, 7.5, 5.0; **MS** *m/z* (ESI^+^) 498 [M+H]^+^; **HRMS** (ESI^+^) *m/z* calcd for C_26_H_33_O_2_N_6_Cl+H^+^ 497.24263 found 497.24345.

**1-[6-[[4-(Cyclopropylamino)-5-(trifluoromethyl)pyrimidin-2-yl]amino]-3,4-dihydro-1*H*-isoquinolin-2-yl]prop-2-en-1-one (2c)**

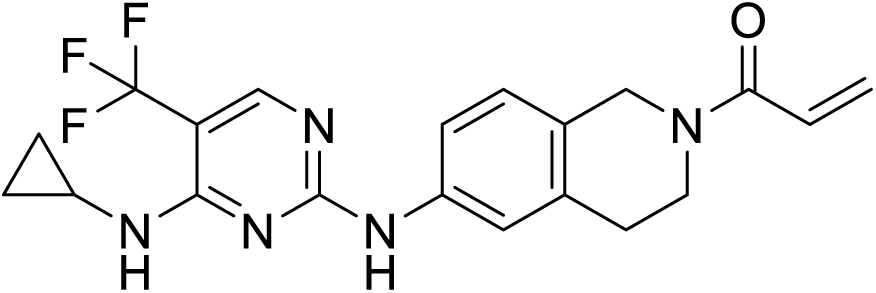

Acryloyl chloride (2.9 µL, 0.036 mmol) followed by DIPEA (16 µL, 0.091 mmol) were added to a solution of **6** (14 mg, 0.0302 mmol) in DMF (0.32 mL). Reaction mixture stirred at rt for 16 h, then purified by preparative-HPLC to afford the title compound (4.26 mg, 0.0106 mmol, 35%) as a white solid. **^1^H NMR** (500 MHz, CDCl_3_) rotameric mixture δ 8.15 (s, 1H), 7.68 (s, 0.4H), 7.60 (s, 0.6H), 7.52 (d, *J* = 8.2 Hz, 0.4H), 7.47 (d, *J* = 8.2 Hz, 0.6H), 7.29 (br s, 1H), 7.12 (d, *J* = 8.3 Hz, 0.6H), 7.07 – 7.02 (d, *J* = 8.3 Hz, 0.4H), 6.65 (dt, *J* = 17.1, 9.8 Hz, 1H), 6.34 (d, *J* = 17.1 Hz, 1H), 5.73 (dd, *J* = 9.8, 1.8 Hz, 1H), 5.37 (s, 1H), 4.76 (s, 1.2H), 4.69 (s, 0.8H), 3.89 (t, *J* = 6.0 Hz, 0.8H), 3.78 (t, *J* = 5.9 Hz, 1.2H), 2.93 – 2.83 (m, 3H), 1.84 (br s, 1H), 0.96 – 0.85 (m, 2H), 0.68 – 0.62 (m, 2H); **^13^C NMR** (126 MHz, CDCl_3_) rotameric mixture δ 165.9, 165.7, 160.8, 160.2, 154.6, 154.6, 138.0, 137.7, 135.6, 134.6, 128.0, 127.9, 127.1, 126.4, 119.5, 119.0, 118.2, 117.9, 47.2, 44.3, 43.5, 40.0, 29.9, 28.8, 24.2, 7.5; **MS** *m/z* (ESI^+^) 404 [M+H]^+^; **HRMS** (ESI^+^) *m/z* calcd for C_20_H_20_ON_5_F_3_+H^+^ 404.16927 found 404.16948.

**(*E*)-1-[6-[[4-(Cyclopropylamino)-5-(trifluoromethyl)pyrimidin-2-yl]amino]-3,4-dihydro-1H-isoquinolin-2-yl]-4-(4-methoxyphenyl)but-2-ene-1,4-dione (2d)**

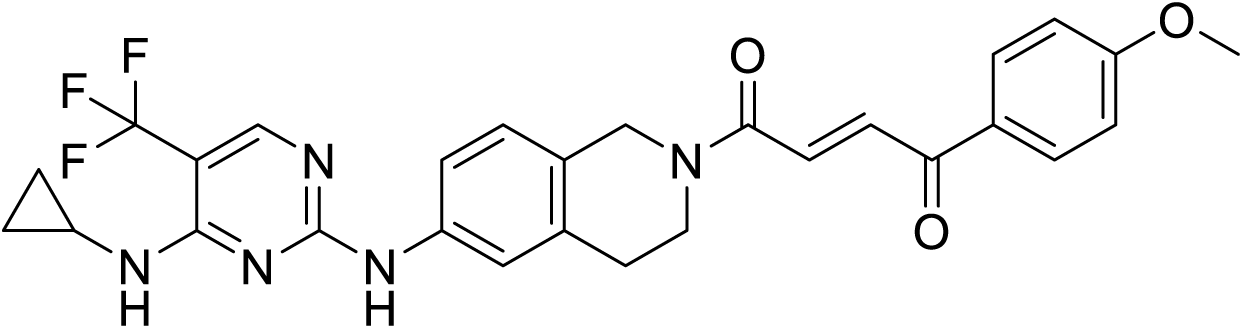

A solution of **6** (113 mg, 0.245 mmol), (*E*)-4-(4-methoxyphenyl)-4-oxo-but-2-enoic acid (61 mg, 0.294 mmol), HATU (168 mg, 0.441 mmol) and DIPEA (130 µL, 0.73 mmol) in DMF (2.4 mL) was stirred at rt for 16 h, then purified by preparative-HPLC to afford the title compound (30 mg, 0.0558 mmol, 23%) as a red solid. **^1^H NMR** (500 MHz, CDCl_3_) δ 8.15 (s, 1H), 8.06 – 7.95 (m, 3H), 7.82 (s, 1H), 7.69 – 7.63 (m, 1H), 7.60 – 7.48 (m, 2H), 7.08 (dd, *J* = 30.2, 8.4 Hz, 1H), 6.95 (dd, *J* = 8.9, 2.0 Hz, 2H), 5.40 (s, 1H), 4.81 (s, 1H), 4.76 (s, 1H), 3.93 (t, *J* = 6.0 Hz, 1H), 3.90 – 3.81 (m, 5H), 2.95 – 2.83 (m, 3H), 0.94 – 0.85 (m, 2H), 0.67 – 0.61 (m, 2H); **^13^C NMR**(126 MHz, CDCl_3_) δ 187.84, 187.81, 164.6, 164.5, 164.2, 160.9, 160.3, 154.7 (q, *J* = 5.0 Hz), 138.4, 138.1, 135.3, 134.4, 134.4, 132.1, 132.0, 131.4, 130.1, 128.2, 127.3, 127.0, 126.5, 126.4, 126.0, 123.9, 121.7, 119.6, 119.2, 118.4, 118.2, 114.2, 99.1 (q, *J* = 32.3 Hz), 55.6, 47.4, 44.7, 43.8, 40.5, 30.0, 28.8, 24.3, 7.5; **^19^F NMR** (471 MHz, CDCl_3_) δ -61.26; **MS** *m/z* (ESI^+^) 538 [M+H]^+^; **HRMS** (ESI^+^) *m/z* calcd for C_28_H_26_O_3_N_5_F_3_+H^+^ 538.20605 found 538.20713.

***N*^4^-Cyclopropyl-5-(trifluoromethyl)-*N*^2^-(2-vinylsulfonyl-3,4-dihydro-1*H*-isoquinolin-6-yl)pyrimidine-2,4-diamine (2e)**

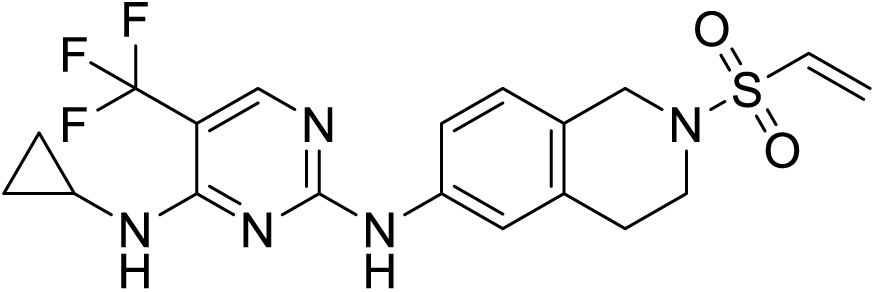

2-Chloroethanesulfonyl chloride (33 µL, 0.31 mmol) in CH_2_Cl_2_ (3.5 mL) was added dropwise to a solution of **6** (122 mg, 0.349 mmol) and NEt_3_ (120 µL, 0.86 mmol) in CH_2_Cl_2_ (3.5 mL). Reaction mixture stirred at 0 °C for 2 h, then concentrated *in vacuo* and purified by preparative-HPLC to afford the title compound (30 mg, 0.0683 mmol, 20%). **^1^H NMR** (500 MHz, CDCl_3_) δ 8.15 (s, 1H), 7.62 (d, *J* = 2.3 Hz, 1H), 7.49 (dd, *J* = 8.4, 2.3 Hz, 1H), 7.45 (br s, 1H), 7.01 (d, *J* = 8.4 Hz, 1H), 6.44 (dd, *J* = 16.6, 9.8 Hz, 1H), 6.30 (d, *J* = 16.6 Hz, 1H), 6.00 (d, *J* = 9.8 Hz, 1H), 5.38 (s, 1H), 4.36 (s, 2H), 3.51 (t, *J* = 6.0 Hz, 2H), 2.93 (t, *J* = 6.0 Hz, 2H), 2.90 – 2.84 (m, 1H), 0.95 – 0.85 (m, 2H), 0.68 – 0.60 (m, 2H); **^13^C NMR** (126 MHz, CDCl3) δ 160.9, 160.3, 154.7, 138.2, 133.9, 133.2, 128.4, 126.8, 126.3, 126.0, 123.9, 119.6, 118.2, 46.9, 43.4, 29.1, 24.4, 7.6; **MS** *m/z* (ESI^+^) 440 [M+H]^+^; **HRMS** (ESI^+^) *m/z* calcd for C_19_H_20_O_2_N_5_F_3_S+H^+^ 440.13626 found 440.13689.

***tert*-Butyl 6-(2,2,2-trifluoro-*N*-methylacetamido)-3,4-dihydroisoquinoline-2(1*H*)-carboxylate (13)**

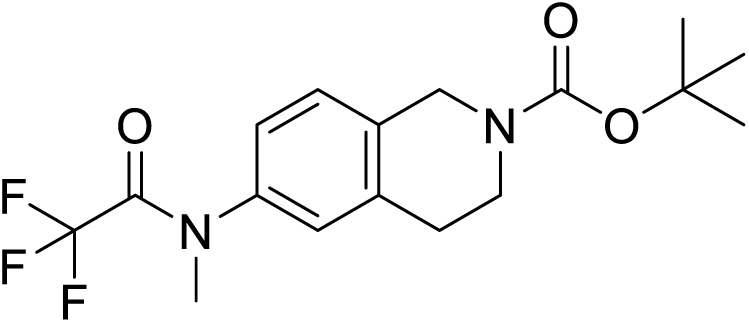

NEt_3_ (560 µL, 4.0 mmol) was added to a solution of 6-Amino-2-*N*-Boc-1,2,3,4-tetrahydro-isoquinoline (500 mg, 2.01 mmol) and trifluoroacetic anhydride (320 µL, 2.4 mmol) in CH_2_Cl_2_ (4 mL) the reaction mixture was stirred at rt for 30 min then concentrated *in vacuo*. The crude trifluoroacetamide intermediate was dissolved in DMF (5 mL) and potassium carbonate (1150 mg, 8.30 mmol), followed by methyl iodide (260 µL, 4.2 mmol) were added. The reaction mixture was heated to 70 °C for 16 h. Reaction mixture cooled to rt and partitioned between EtOAc and water. Organic layer washed with water x2, followed by brine, dried over Na_2_SO_4_ then concentrated *in vacuo*. The crude residue was purified by flash column chromatography (heptane/EtOAc, 0-50%) to afford the title compound (595 mg, 1.55 mmol, 82% over two steps) as a white solid. **^1^H NMR** (500 MHz, CDCl_3_) δ 7.16 (d, *J* = 8.2 Hz, 1H), 7.08 – 6.99 (m, 2H), 4.60 (s, 2H), 3.66 (t, *J* = 6.0 Hz, 2H), 3.33 (s, 3H), 2.84 (t, *J* = 6.0 Hz, 2H), 1.50 (s, 9H); **^13^C NMR** (126 MHz, CDCl_3_) rotameric mixture δ 157.1, 156.8, 154.8, 138.8, 136.3, 134.8, 127.5, 125.1, 117.5, 115.2, 80.1, 45.7, 45.1, 41.4, 40.2, 39.8, 28.9, 28.5; **^19^F NMR** (471 MHz, CDCl_3_) δ -66.97.

***tert*-Butyl 6-(methylamino)-3,4-dihydroisoquinoline-2(1*H*)-carboxylate (14)**

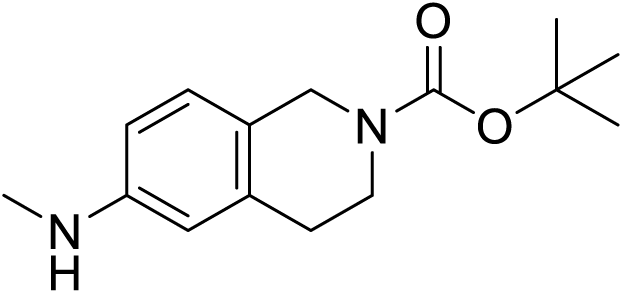

A solution of **13** (514 mg, 1.43 mmol) and potassium carbonate (397 mg, 2.87 mmol) in methanol (4.5 mL) and water (1.5 mL) was heated to 70 °C for 1 h. Reaction mixture extracted with EtOAc x2 and combined organics washed with brine, dried over Na_2_SO_4_ and concentrated *in vacuo* to afford the title compound (365 mg, 1.39 mmol, 97%) as a colourless oil. **^1^H NMR** (500 MHz, CDCl_3_) δ 6.92 (d, *J* = 8.3 Hz, 1H), 6.48 (dd, *J* = 8.3, 2.5 Hz, 1H), 6.38 (d, *J* = 2.4 Hz, 1H), 4.47 (s, 2H), 3.63 (br s, 2H), 2.82 (s, 3H), 2.76 (t, *J* = 6.1 Hz, 2H), 1.49 (s, 9H); **^13^C NMR** (126 MHz, CDCl_3_) δ 155.1, 148.0, 135.8, 127.1, 122.6, 111.8, 111.4, 79.6, 45.6, 44.8, 42.1, 40.9, 31.0, 29.5, 28.6; **MS** *m/z* (ESI^+^) 264 [M+H]^+^.

***tert*-Butyl 6-((4-chloro-5-(trifluoromethyl)pyrimidin-2-yl)(methyl)amino)-3,4-dihydroisoquinoline-2(1*H*)-carboxylate (15)**

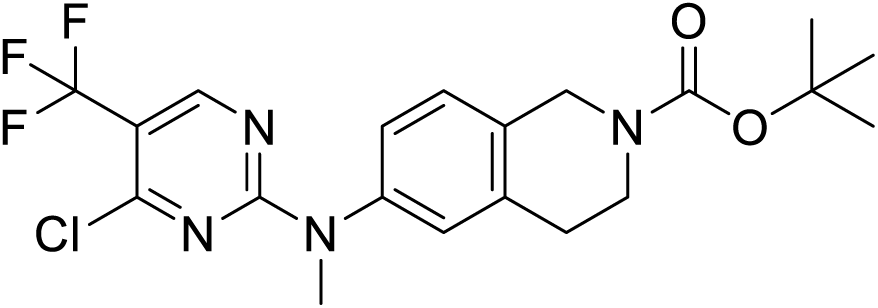

A solution of 2,4-dichloro-5-(trifluoromethyl)pyrimidine (326 mg, 1.50 mmol) and ZnCl_2_ (246 mg, 1.80 mmol) in DCE/*t*-BuOH (1:1) (12.5 mL) was stirred at 0 °C for 1 h. Then a solution of **14** (394 mg, 1.50 mmol) and NEt_3_ (250 µL, 1.8 mmol) in DCE/*t*-BuOH (1:1) (2.5 mL) was added. The solution was stirred at 0 °C for a further 3 h, then concentrated *in vacuo* to afford crude product, which was purified by flash column chromatography (heptane/EtOAc, 0→50%) to afford the title compound (635 mg, 1.43 mmol, 95%) as a white solid. **^1^H NMR** (500 MHz, CDCl_3_) δ 8.42 (s, 1H), 7.18 (d, *J* = 8.2 Hz, 1H), 7.09 (dd, *J* = 8.2, 2.2 Hz, 1H), 7.05 (d, *J* = 2.2 Hz, 1H), 4.60 (s, 2H), 3.68 (br s, 2H), 3.54 (s, 3H), 2.85 (t, *J* = 5.9 Hz, 2H), 1.50 (s, 9H); **^13^C NMR** (126 MHz, CDCl_3_) rotameric mixture δ 162.3, 159.2, 157.1, 157.1, 154.9, 142.0, 136.3, 127.6, 126.6, 124.5, 124.2, 122.0, 119.9, 112.0, 111.7, 111.5, 80.1, 46.0, 43.3, 41.7, 39.5, 28.6; **MS** *m/z* (ESI^+^) 443 [M+H]^+^.

***tert*-Butyl 6-((4-(cyclopropylamino)-5-(trifluoromethyl)pyrimidin-2-yl)(methyl)amino)-3,4-dihydroisoquinoline-2(1*H*)-carboxylate (16)**

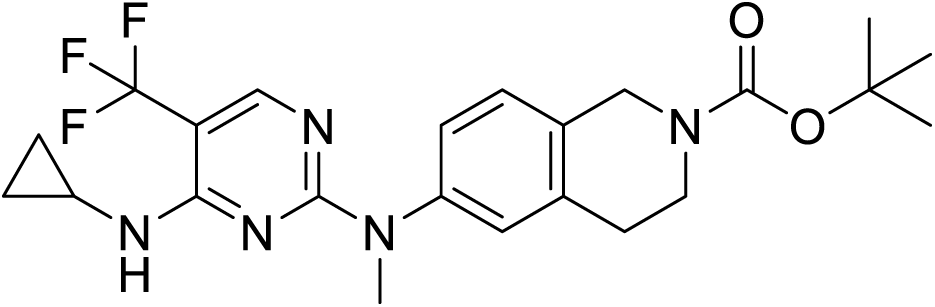

A solution of **15** (620 mg, 1.40 mmol), cyclopropylamine (230 µL, 3.4 mmol) and DIPEA (590 µL, 3.4 mmol) in DMF (9 mL) was heated to 100 °C under microwave irradiation for 15 min. The reaction mixture was concentrated *in vacuo* to afford crude product, which was then purified by flash column chromatography (heptane/EtOAc, 0→50%) to afford the title compound (600 mg, 1.30 mmol, 92%). **^1^H NMR** (500 MHz, CDCl_3_) δ 8.08 (s, 1H), 7.19 – 7.06 (m, 3H), 5.20 (br s, 1H), 4.57 (s, 2H), 3.65 (br s, 2H), 3.53 (s, 3H), 2.88 – 2.72 (m, 4H), 1.49 (s, 10H), 0.80 – 0.72 (m, 2H), 0.58 – 0.51 (m, 2H); **^13^C NMR** (126 MHz, CDCl_3_) δ 162.8, 159.8, 155.0, 154.8 (q, *J* = 5.0 Hz), 143.4, 135.5, 131.6, 126.9, 126.5, 125.0, 124.4, 97.9, 97.6, 79.9, 46.0, 41.8, 40.6, 38.8, 28.6, 24.0, 7.3; **MS** *m/z* (ESI^+^) 465 [M+H]^+^.

***N*^4^-Cyclopropyl-*N*^2^-methyl-*N*^2^-(1,2,3,4-tetrahydroisoquinolin-6-yl)-5-(trifluoromethyl)pyrimidine-2,4-diamine (17)**

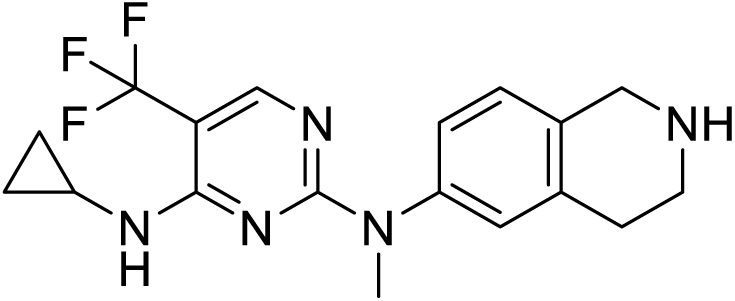

TFA (670 µL, 8.7 mmol) was added to a solution of **16** (600 mg, 1.30 mmol) in CH_2_Cl_2_ (6.5 mL). Reaction mixture stirred at rt for 2 h, then concentrated *in vacuo*. Residue passed through an SCX column to afford the title compound (420 mg, 1.16 mmol, 89%) as a colourless oil. **^1^H NMR** (500 MHz, CDCl_3_) δ 8.08 (s, 1H), 7.09 (dd, *J* = 8.1, 2.3 Hz, 1H), 7.06 – 6.98 (m, 2H), 5.23 (br s, 1H), 4.00 (s, 2H), 3.52 (s, 3H), 3.12 (t, *J* = 6.0 Hz, 2H), 2.82 – 2.71 (m, 3H), 2.29 (br s, 1H), 0.79 – 0.69 (m, 2H), 0.56 – 0.49 (m, 2H); **^13^C NMR** (126 MHz, CDCl_3_) δ 162.7, 159.7, 154.7 (q, *J* = 5.0 Hz), 143.0, 135.3, 133.5, 127.2, 126.7, 124.4, 97.5 (q, *J* = 31.8 Hz), 48.0, 43.6, 38.6, 29.2, 23.9, 7.1; **^19^F NMR** (471 MHz, CDCl_3_) δ -60.85; **MS** *m/z* (ESI^+^) 364 [M+H]^+^.

**2-Chloro-1-(6-((4-(cyclopropylamino)-5-(trifluoromethyl)pyrimidin-2-yl)(methyl)amino)-3,4-dihydroisoquinolin-2(1*H*)-yl)ethan-1-one (3a)**

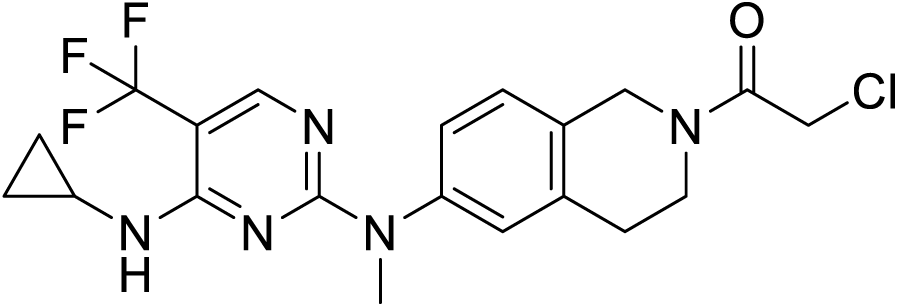

Chloroacetyl chloride (31 µL, 0.39 mmol) was added to a solution of **17** (128 mg, 0.352 mmol) and DIPEA (180 µL, 1.1 mmol) in DMF (3.4 mL). Reaction mixture stirred at rt for 16 h, then concentrated *in vacuo*. The crude product was purified by preparative-HPLC to afford the title compound (63 mg, 0.143 mmol, 41%) as an off-white solid. **^1^H NMR** (500 MHz, CDCl_3_) 3:2 rotameric mixture δ 8.08 (s, 1H), 7.24 – 7.11 (m, 3H), 5.22 (s, 1H), 4.74 (s, 1.2H), 4.69 (s, 0.8H), 4.16 (d, *J* = 4.0 Hz, 2H), 3.85 (t, *J* = 6.0 Hz, 0.8H), 3.76 (t, *J* = 6.0 Hz, 1.2H), 3-54 (d, *J* = 3.3 Hz, 3H), 2.98 (t, *J* = 6.0 Hz, 1.2H), 2.89 (t, *J* = 6.0 Hz, 0.8H), 2.81 – 2.73 (m, 1H), 1.35 – 1.21 (m, 3H), 0.88 (t, *J* = 6.9 Hz, 1H), 0.75 (t, *J* = 6.4 Hz, 2H), 0.59 – 0.50 (m, 2H); **^13^C NMR** (126 MHz, CDCl_3_) rotameric mixture δ 165.7, 165.5, 159.8, 154.7 (q, *J* = 5.0 Hz), 144.2, 143.8, 135.4, 134.4, 130.3, 129.6, 127.3, 127.0, 126.6, 126.6, 126.5, 125.5, 125.3, 124.3, 97.9 (q, *J* = 32.1 Hz), 47.8, 44.7, 43.8, 41.5, 41.3, 40.5, 38.71, 38.68, 32.0, 29.6, 29.1, 28.5, 24.0, 22.8, 14.2, 7.3; **^19^F NMR** (471 MHz, CDCl_3_) δ -60.9; **MS** *m/z* (ESI^+^) 440 [M+H]^+^; **HRMS** (ESI^+^) *m/z* calcd for C_20_H_21_ClF_3_N_5_O+H^+^ 440.14595 found 440.14621.

**(*E*)-1-(6-((4-(cyclopropylamino)-5-(trifluoromethyl)pyrimidin-2-yl)(methyl)amino)-3,4-dihydroisoquinolin-2(1*H*)-yl)-4-(4-methoxyphenyl)but-2-ene-1,4-dione (3b)**

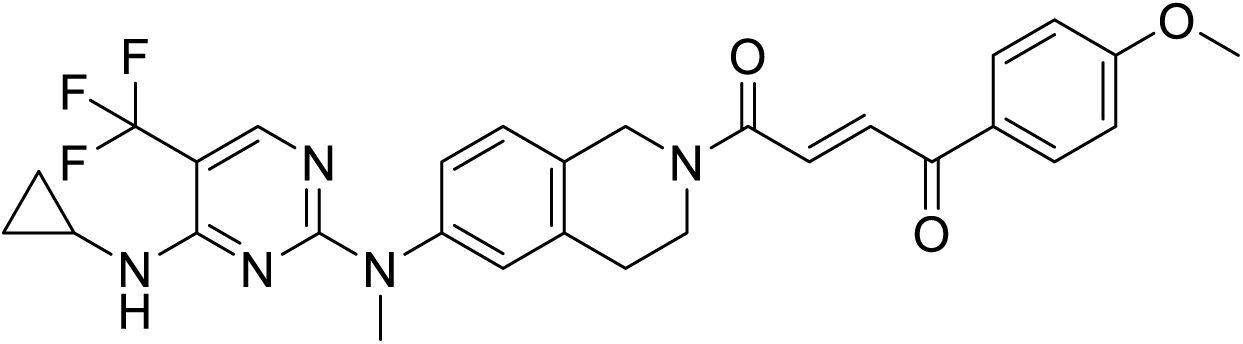

A solution of **17** (88 mg, 0.242 mmol), (*E*)-4-(4-methoxyphenyl)-4-oxo-but-2-enoic acid (60 mg, 0.291 mmol), HATU (166 mg, 0.436 mmol) and DIPEA (130 µL, 0.73 mmol) in DMF (1.2 mL) was stirred at rt for 16 h, then purified by preparative-HPLC to afford the title compound (91 mg, 0.0558 mmol, 23%) as an off-white solid. **^1^H NMR** (500 MHz, CDCl_3_) δ 8.09 – 8.02 (m, 3H), 7.98 (dd, *J* = 14.9, 6.4 Hz, 1H), 7.56 (dd, *J* = 14.9, 8.5 Hz, 1H), 7.22 – 7.11 (m, 3H), 7.00 – 6.92 (m, 2H), 5.24 (s, 1H), 4.84 (s, 1H), 4.80 (s, 1H), 3.93 (t, *J* = 6.0 Hz, 1H), 3.89 – 3.83 (m, 4H), 3.53 (s, 3H), 2.98 – 2.89 (m, 2H), 2.80 – 2.72 (m, 1H), 0.80 – 0.69 (m, 2H), 0.59 – 0.49 (m, 2H); **^13^C NMR** (126 MHz, CDCl_3_) δ 187.7, 187.7, 164.6, 164.4, 164.2, 162.5, 162.5, 159.7, 154.6 (q, *J* = 5.0 Hz), 144.0, 143.7, 135.4, 134.5, 134.4, 132.0, 131.9, 131.3, 130.5, 130.0, 129.6, 128.5, 127.2, 126.8, 126.6, 126.5, 126.4, 125.4, 125.2, 124.2, 122.1, 114.1, 97.7 (dq, *J* = 31.8, 6.0 Hz), 55.6, 47.5, 44.7, 43.6, 40.3, 38.6, 38.6, 29.7, 28.5, 23.9, 7.2; **^19^F NMR** (471 MHz, CDCl_3_) δ -60.9; **MS** *m/z* (ESI^+^) 553 [M+H]^+^; **HRMS** *m/z* calcd for C_29_H_28_O_3_N_5_F_3_+H^+^ 552.22170 found 552.22430.

**1-(6-((4-(cyclopropylamino)-5-(trifluoromethyl)pyrimidin-2-yl)amino)-3,4-dihydroisoquinolin-2(1H)-yl)propan-1-one (3c)**

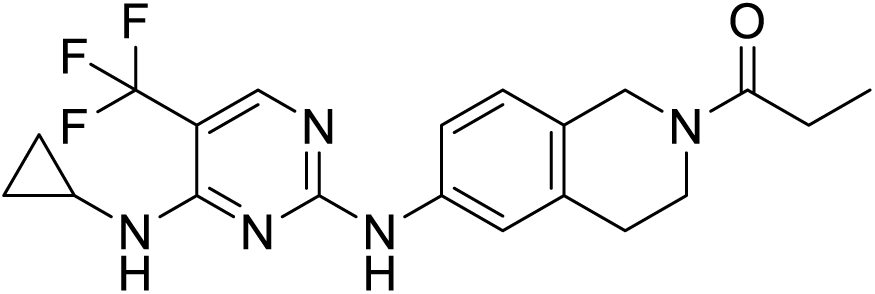

Propionyl chloride (12 µL, 0.14 mmol) was added to a solution of **6** (45 mg, 0.129 mmol) and DIPEA (67 µL, 0.39 mmol) in DMF (1.5 mL). Reaction mixture stirred at rt for 16 h, then concentrated *in vacuo*. The crude product was purified by preparative-HPLC to afford the title compound (12 mg, 0.0296 mmol, 23%) as a white solid. **^1^H NMR** (500 MHz, CDCl3) 3:2 rotameric mixture δ 8.15 (s, 1H), 7.67 (s, 0.4H), 7.59 (s, 0.6H), 7.55 – 7.34 (m, 2H), 7.07 (dd, *J* = 26.3, 8.3 Hz, 1H), 5.38 (s, 1.2H), 4.70 (s, 0.8H), 4.58 (s, 1H), 3.83 (t, *J* = 5.9 Hz, 1H), 3.67 (t, *J* = 5.9 Hz, 1H), 2.85 (m, 3H), 2.43 (dq, *J* = 7.3, 3.4 Hz, 2H), 1.18 (q, *J* = 7.3 Hz, 3H), 0.94 – 0.85 (m, 2H), 0.68 – 0.61 (m, 2H); **^13^C NMR** (126 MHz, CDCl_3_) rotameric mixture δ 172.9, 172.8, 161.0, 160.3, 154.7 (q, *J* = 5.2 Hz), 138.1, 137.7, 135.9, 134.7, 128.3, 127.1, 126.5, 126.1, 123.9, 119.7, 119.1, 118.3, 117.9, 47.1, 44.0, 43.2, 39.7, 30.0, 29.0, 27.1, 27.0, 24.4, 9.6, 9.5, 7.6; **^19^F NMR** (471 MHz, CDCl_3_) δ -61.29; **MS** *m/z* (ESI^+^) 406 [M+H]^+^; **HRMS** *m/z* calcd for C_20_H_22_ON_5_F_3_+H^+^ 406.18492 found 406.18582.

**3-(4-(4-((5-Chloro-4-((2-(isopropylsulfonyl)phenyl)amino)pyrimidin-2-yl)amino)phenyl)piperazine-1-carbonyl)benzenesulfonyl fluoride (S2a)**

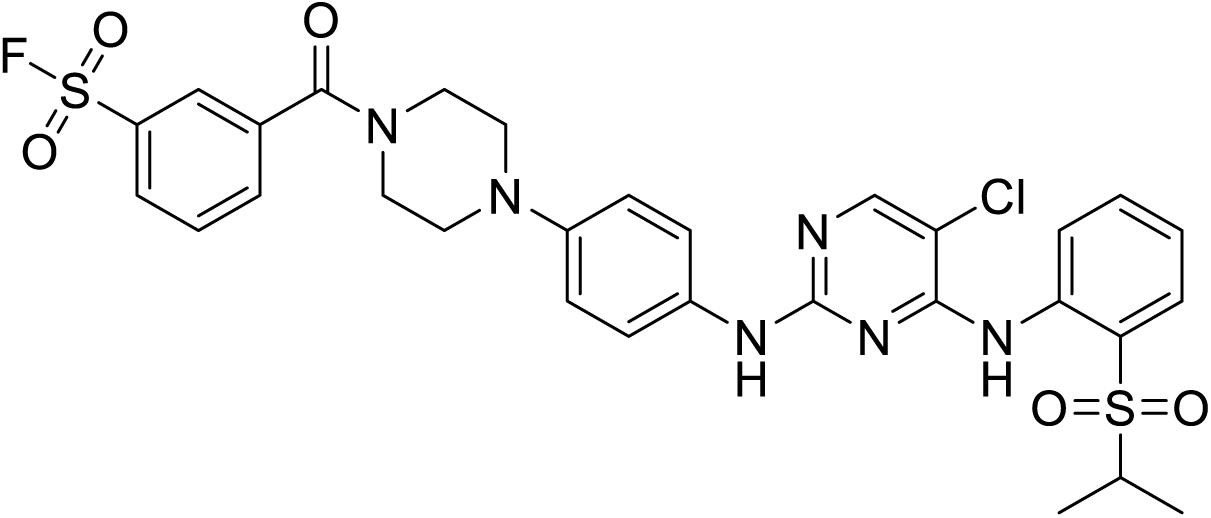

A solution of **2** trifluoroacetamide (191 mg, 0.318 mmol), 3-fluorosulfonylbenzoic acid (50 mg, 0.245 mmol), HATU (158 mg, 0.416 mmol) and DIPEA (110 µL, 0.63 mmol) in DMF (2.4 mL) was stirred at rt for 16 h. Reaction mixture concentrated *in vacuo* and purified by flash column chromatography (heptane/EtOAc, 0→100%) followed by preparative-HPLC to afford the title compound (60 mg, 0.0894 mmol, 37%) as a white solid. **^1^H NMR** (500 MHz, CDCl3) rotameric mixture δ 9.63 (s, 1H), 8.58 (dd, *J* = 8.6, 1.1 Hz, 1H), 8.11 – 8.06 (m, 3H), 7.91 – 7.83 (m, 2H), 7.73 (t, *J* = 7.7 Hz, 1H), 7.55 (m, 1H), 7.45 – 7.39 (m, 3H), 7.25 – 7.18 (m, 1H), 6.93 – 6.86 (m, 2H), 3.97 (br s, 2H), 3.58 (br s, 2H), 3.29 – 3.02 (m, 4H), 1.30 (d, *J* = 6.9 Hz, 6H); **^13^C NMR** (126 MHz, CDCl_3_) rotameric mixture δ 167.4, 165.9, 157.9, 155.5, 154.9, 146.9, 138.5, 137.5, 134.5, 134.3, 133.8, 133.6, 132.9, 131.3, 130.3, 129.6, 127.3, 124.5, 123.5, 123.2, 122.0, 117.7, 116.5, 106.0, 55.7, 50.6, 50.3, 47.8, 42.5, 15.4; **^19^F NMR** (471 MHz, CDCl_3_) δ 66.11; **MS** *m/z* (ESI^+^) 673 [M+H]^+^; **HRMS** (ESI^+^) *m/z* calcd for C_30_H_30_O_5_N_6_ClFS_2_+H^+^ 673.14644 found 673.14844.

**Methyl 4-(*N*-methylsulfamoyl)benzoate (18)**

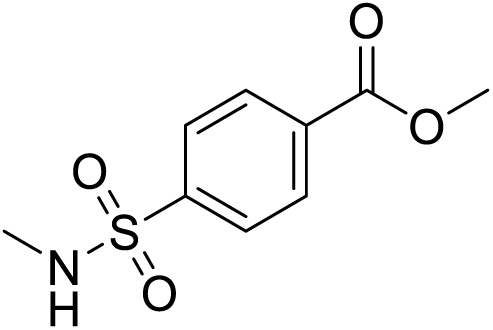

A solution of methyl 4-chlorosulfonylbenzoate (317 mg, 1.35 mmol), 2 M methylamine in THF (1.01 mL, 2.02 mmol) and DIPEA (570 µL, 3.2 mmol) in CH_2_Cl_2_ (6.8 mL) was stirred at rt for 1 h. Reaction mixture concentrated *in vacuo* and purified by flash column chromatography (CH_2_Cl_2_/MeOH, 0→5%) to afford the title compound (244 mg, 1.06 mmol) as a white solid. **^1^H NMR** (500 MHz, CDCl_3_) δ 8.22 – 8.14 (m, 2H), 7.96 – 7.90 (m, 2H), 4.59 (q, *J* = 5.4 Hz, 1H), 3.96 (s, 3H), 2.69 (d, *J* = 5.4 Hz, 3H); **^13^C NMR** (126 MHz, CDCl_3_) δ 165.8, 143.0, 134.1, 130.5, 127.4, 52.8, 29.5.

**Methyl 4-(*N*-methyl-*N*-((triisopropylsilyl)ethynyl)sulfamoyl)benzoate (19)**

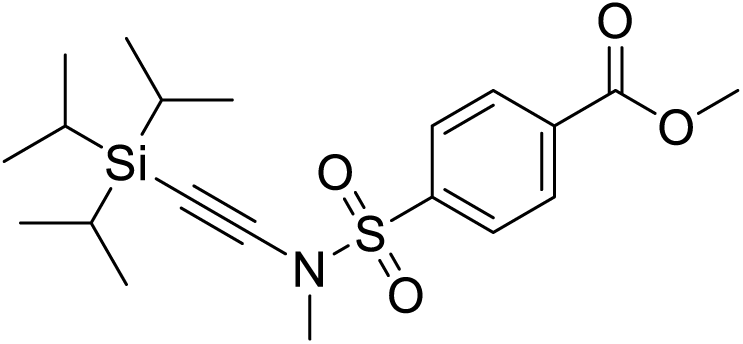

A degassed suspension of **18** (230 mg, 1.00 mmol), (bromoethynyl)triisopropylsilane (290 µL, 1.2 mmol), 1,10-phenanthroline (36 mg, 0.201 mmol), CuSO_4_.5H_2_O (25 mg, 0.100 mmol) and K_2_CO_3_ (277 mg, 2.00 mmol) in toluene (5 mL) was heated to 85 °C for 24 h. Reaction mixture cooled to rt, diluted with CH_2_Cl_2_ and filtered through celite, the filtrate was then concentrated *in vacuo*. Crude residue purified by flash column chromatography (heptane/EtOAc, 0→60%) to afford the title compound (349 mg, 0.852 mmol, 85%) as a white solid. **^1^H NMR** (500 MHz, CDCl_3_) δ 8.23 – 8.18 (m, 2H), 8.01 – 7.96 (m, 2H), 3.98 (s, 3H), 3.11 (s, 3H), 1.04 (app. s, 21H); ^1**3**^**C NMR** (126 MHz, CDCl_3_) δ 165.6, 140.1, 134.9, 130.4, 127.9, 97.3, 68.3, 52.9, 39.6, 18.7, 11.4; **MS** *m/z* (ESI^+^) 410 [M+H]^+^.

**4-(*N*-Methyl-*N*-((triisopropylsilyl)ethynyl)sulfamoyl)benzoic acid (20)**

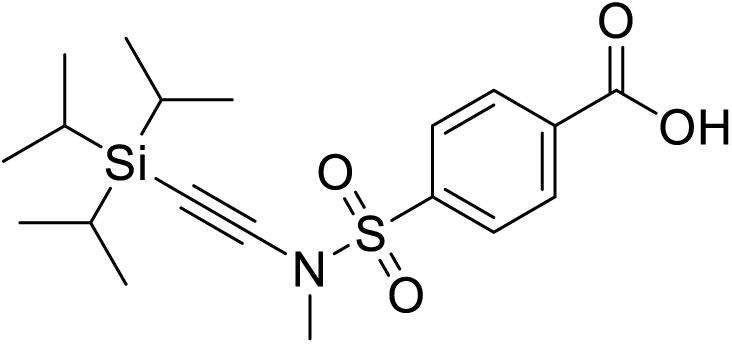

A solution of **19** (335 mg, 0.818 mmol) and LiOH.H_2_O (172 mg, 4.09 mmol) in THF (3.75 mL) and water (1.25 mL) was stirred at rt for 3 h. Reaction mixture acidified to pH 3 with 2 M HCl (aq.) and extracted with EtOAc x3. Combined organics dried over Na_2_SO_4_ and concentrated *in vacuo* to afford the title compound (319 mg, 0.806 mmol, 99%) as a white solid. **^1^H NMR** (500 MHz, CDCl_3_) δ 8.32 – 8.24 (m, 2H), 8.07 – 8.00 (m, 2H), 3.14 (s, 3H), 1.05 (s, 21H); ^13^C NMR (126 MHz, CDCl_3_) δ 170.2, 141.0, 133.8, 131.0, 128.1, 97.2, 68.5, 39.7, 18.7, 11.4; **MS** *m/z* (ESI^-^) 394 [M-H]^-^.

**4-(4-(4-((5-chloro-4-((2-(isopropylsulfonyl)phenyl)amino)pyrimidin-2-yl)amino)phenyl)piperazine-1-carbonyl)-*N*-methyl-*N*-((triisopropylsilyl)ethynyl)benzenesulfonamide (21)**

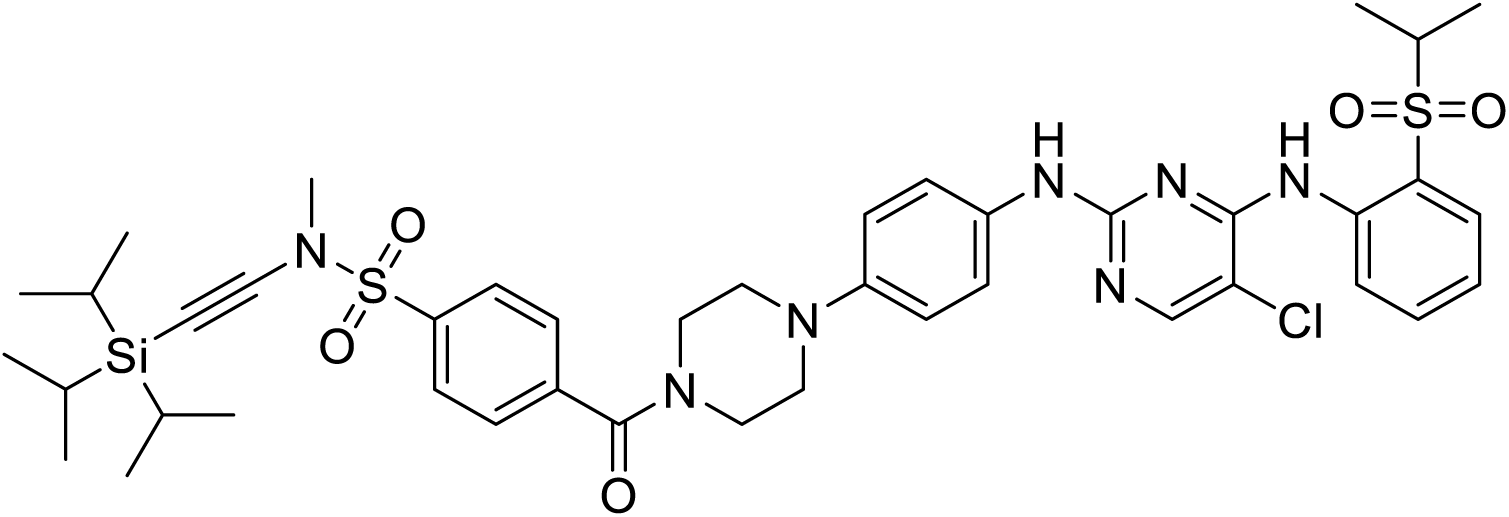

A solution of **2** trifluoroacetamide (218 mg, 0.363 mmol), **20** (172 mg, 0.435 mmol), HATU (194 mg, 0.508 mmol) and DIPEA (190 µL 1.1 mmol) in DMF (3.6 mL) was stirred at rt for 16 h. Reaction mixture concentrated *in vacuo* and purified by flash column chromatography (heptane/EtOAc, 0→100%) to afford the title compound (101 mg, 0.117 mmol, 32%) as an off-white solid. **^1^H NMR** (500 MHz, CDCl_3_) δ 9.64 (s, 1H), 8.59 (d, *J* = 8.4 Hz, 1H), 8.10 (s, 1H), 7.99 (d, *J* = 8.4 Hz, 2H), 7.89 (dd, *J* = 7.9, 1.7 Hz, 1H), 7.61 (d, *J* = 8.4 Hz, 2H), 7.55 (t, *J* = 7.1 Hz, 1H), 7.46 – 7.40 (m, 2H), 7.34 (br s, 1H), 7.23 (t, *J* = 7.1 Hz, 1H), 6.93 – 6.87 (m, 2H), 3.97 (br s, 2H), 3.54 (br s, 2H), 3.23 (hept, *J* = 7.1 Hz, 2H), 3.14 – 3.03 (m, 5H), 1.30 (d, *J* = 7.1 Hz, 6H), 1.04 (s, 21H); ^13^C NMR (126 MHz, CDCl_3_) δ 168.5, 158.0, 155.5, 155.1, 147.0, 140.9, 138.5, 137.5, 134.5, 132.9, 131.4, 128.3, 127.8, 124.5, 123.5, 123.2, 122.0, 117.7, 106.0, 97.4, 68.2, 55.7, 50.8, 50.3, 47.7, 42.3, 39.6, 18.7, 15.4, 11.4; **MS** *m/z* (ESI^+^) 864 [M+H]^+^.

**4-(4-(4-((5-chloro-4-((2-(isopropylsulfonyl)phenyl)amino)pyrimidin-2-yl)amino)phenyl)piperazine-1-carbonyl)-*N*-ethynyl-*N*-methylbenzenesulfonamide (S2b)**

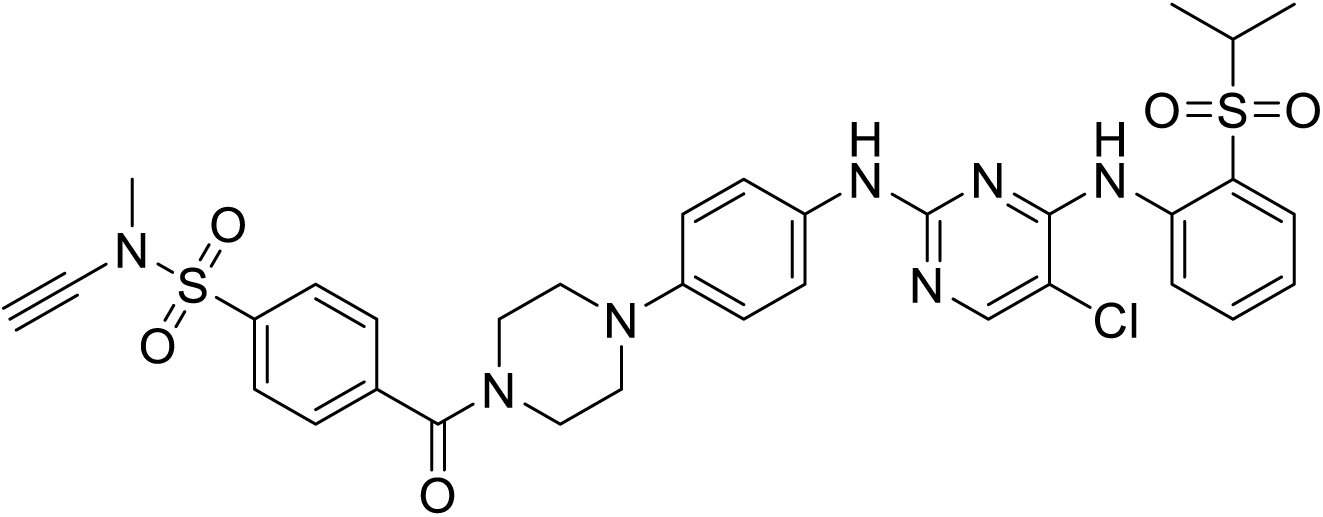

1 M TBAF in THF (180 µL, 0.18 mmol) was added to a solution of **21** (101 mg, 0.117 mmol) in THF (6 mL). Reaction mixture stirred at rt for 3 h, then concentrated *in vacuo*. Residue purified by flash column chromatography (heptane/EtOAc, 0→100%) to afford the title compound (61 mg, 0.0861 mmol, 74%) as a white solid. **^1^H NMR** (500 MHz, CDCl_3_) rotameric mixture δ 9.62 (s, 1H), 8.59 (dd, *J* = 8.6, 1.1 Hz, 1H), 8.12 (s, 1H), 8.03 – 7.97 (m, 2H), 7.90 (dd, *J* = 7.9, 1.7 Hz, 1H), 7.68 – 7.61 (m, 2H), 7.57 (ddd, *J* = 8.6, 7.3, 1.7 Hz, 1H), 7.46 – 7.40 (m, 2H), 7.25 – 7.21 (m, 1H), 7.00 (s, 1H), 6.94 – 6.87 (m, 2H), 3.98 (br s, 2H), 3.56 (br s, 2H), 3.23 (hept, *J* = 6.8 Hz, 2H), 3.14 – 3.06 (m, 5H), 1.31 (d, *J* = 6.8 Hz, 6H); **^13^C NMR** (126 MHz, CDCl_3_) rotameric mixture δ 168.4, 158.1, 155.5, 155.4, 147.0, 141.1, 138.6, 137.5, 134.5, 132.9, 131.4, 128.3, 128.0, 124.6, 123.5, 123.2, 121.9, 117.8, 106.2, 58.1, 55.7, 50.8, 50.3, 47.7, 42.4, 39.1, 15.5; **MS** *m/z* (ESI^+^) 708 [M+H]^+^; **HRMS** (ESI^+^) *m/z* calcd for C_33_H_34_O_5_N_7_ClS_2_+H^+^ 708.18241 found 708.18403.

**(4-(4-((5-Chloro-4-((2-(isopropylsulfonyl)phenyl)amino)pyrimidin-2-yl)amino)phenyl)piperazin-1-yl)(1-hydroxy-1,3-dihydrobenzo[*c*][1,2]oxaborol-6-yl)methanone (S2c)**

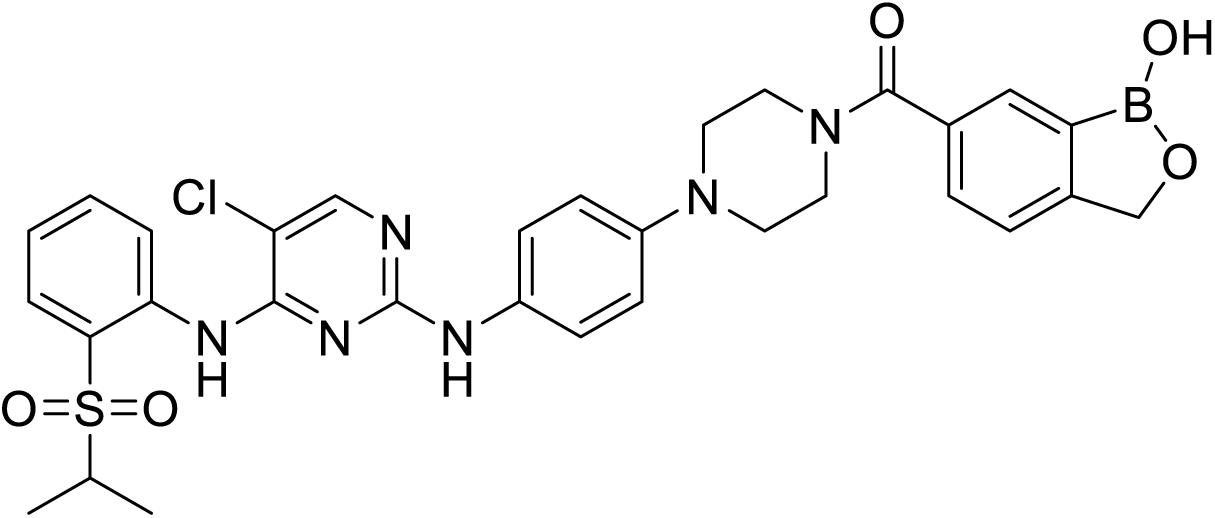

A solution of **2** (100 mg, 0.205 mmol), 1-hydroxy-1,3-dihydrobenzo[*c*][1,2]oxaborole-6-carboxylic acid (38 mg, 0.205 mmol), EDC hydrochloride (63 mg, 0.329 mmol) and DMAP (1 mg, 0.0082 mmol) in CH_2_Cl_2_ (1 mL) was stirred at rt for 16 h. The crude product was purified by preparative-HPLC to afford the title compound (72 mg, 0.111 mmol, 54%) as a white solid. **^1^H NMR** (500 MHz, CDCl3) rotameric mixture δ 9.68 (s, 1H), 8.57 (dd, *J* = 8.5, 1.1 Hz, 1H), 8.07 (s, 1H), 7.95 (s, 1H), 7.90 (dd, *J* = 8.0, 1.7 Hz, 1H), 7.76 (br s, 1H), 7.59 – 7.51 (m, 2H), 7.45 – 7.38 (m, 3H), 7.28 – 7.23 (m, 1H), 6.91 – 6.86 (m, 2H), 5.12 (s, 2H), 3.98 (br s, 2H), 3.63 (br s, 2H), 3.31 – 2.97 (m, 5H), 1.31 (d, *J* = 6.9 Hz, 6H); **^13^C NMR** (126 MHz, CDCl_3_) rotameric mixture δ 171.1, 165.3, 157.8, 155.7, 154.3, 147.3, 138.4, 134.6, 134.3, 132.6, 131.4, 130.0, 129.9, 124.8, 123.8, 123.5, 122.2, 121.5, 117.6, 105.8, 71.1, 55.8, 50.5, 48.1, 42.6, 15.5; **MS** *m/z* (ESI^+^) 647 [M+H]^+^; **HRMS** (ESI^+^) *m/z* calcd for C_31_H_32_O_5_N_6_BClS+H^+^ 647.20092 found 647.20303.

**4-(4-(4-((5-Chloro-4-((2-(isopropylsulfonyl)phenyl)amino)pyrimidin-2-yl)amino)phenyl)piperazine-1-carbonyl)-2-hydroxybenzaldehyde (S2d)**

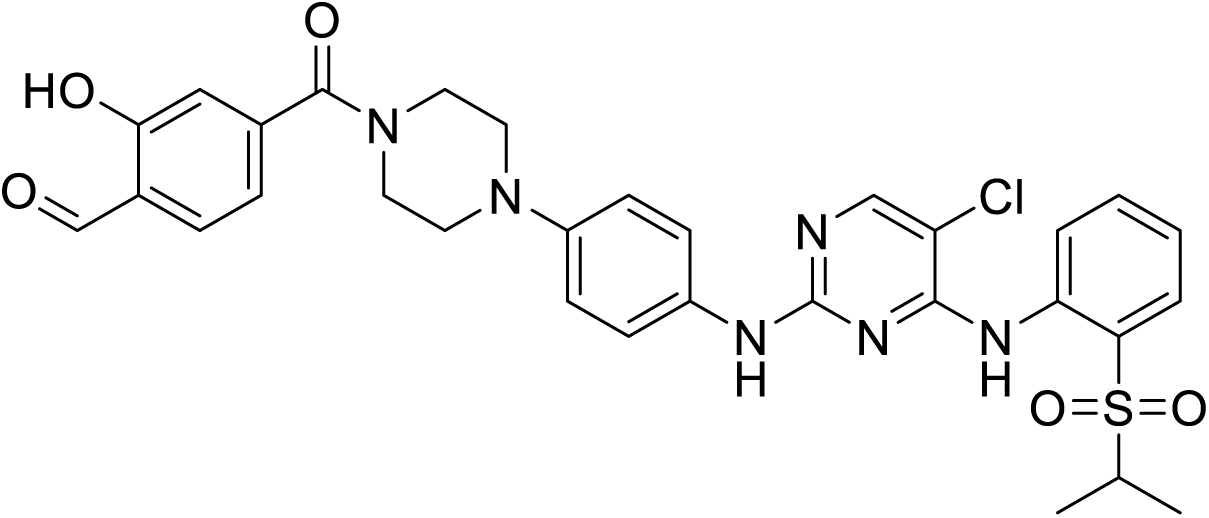

A solution of **2** (43 mg, 0.0883 mmol), 4-formyl-3-hydroxy-benzoic acid (15 mg, 0.0883 mmol) and EDC hydrochloride (22 mg, 0.115 mmol) in CH_2_Cl_2_ (0.5 mL) was stirred at rt for 16 h. The crude product was purified by preparative-HPLC to afford the title compound (32 mg, 0.0501 mmol, 57%) as a white solid. **^1^H NMR** (500 MHz, CDCl_3_) rotameric mixture δ 9.94 (s, 1H), 9.63 (s, 1H), 8.58 (d, *J* = 8.4 Hz, 1H), 8.10 (s, 1H), 7.90 (dd, *J* = 8.0, 1.7 Hz, 1H), 7.65 (d, *J* = 7.8 Hz, 1H), 7.56 (ddd, *J* = 8.4, 7.3, 1.7 Hz, 1H), 7.46 – 7.39 (m, 2H), 7.26 – 7.22 (m, 2H), 7.07 (dd, *J* = 7.8, 1.4 Hz, 1H), 7.04 – 7.00 (m, 1H), 6.93 – 6.87 (m, 2H), 3.95 (br s, 2H), 3.56 (br s, 2H), 3.29 – 3.17 (m, 3H), 3.08 (br s, 2H), 1.31 (d, *J* = 6.9 Hz, 6H); **^13^C NMR** (126 MHz, CDCl_3_) δ 196.3, 168.5, 161.7, 158.0, 155.5, 155.1, 147.1, 143.9, 138.5, 134.5, 134.4, 132.8, 131.4, 124.6, 123.6, 123.3, 122.0, 121.1, 118.4, 117.8, 116.2, 106.1, 55.8, 50.8, 50.4, 47.5, 42.2, 15.5; **MS** *m/z* (ESI^+^) 635 [M+H]^+^; **HRMS** (ESI^+^) *m/z* calcd for C_31_H_31_O_5_N_6_ClS+H^+^ 635.18379 found 635.18616.

**NMR Spectra**

